# RNAxplorer: Harnessing the Power of Guiding Potentials to Sample RNA Landscapes

**DOI:** 10.1101/2020.07.03.186882

**Authors:** Gregor Entzian, Ivo Hofacker, Yann Ponty, Ronny Lorenz, Andrea Tanzer

## Abstract

**Motivation:** Predicting the folding dynamics of RNAs is a computationally difficult problem, first and foremost due to the combinatorial explosion of alternative structures in the folding space. Abstractions are therefore needed to simplify downstream analyses, and thus make them computationally tractable. This can be achieved by various structure sampling algorithms. However, current sampling methods are still time consuming and frequently fail to represent key elements of the folding space.

**Method:** We introduce `RNAxplorer`, a novel adaptive sampling method to efficiently explore the structure space of RNAs. `RNAxplorer` uses dynamic programming to perform an efficient Boltzmann sampling in the presence of guiding potentials, which are accumulated into pseudo-energy terms and reflect similarity to already well-sampled structures. This way, we effectively steer sampling towards underrepresented or unexplored regions of the structure space.

**Results:** We developed and applied different measures to benchmark our sampling methods against its competitors. Most of the measures show that `RNAxplorer` produces more diverse structure samples, yields rare conformations that may be inaccessible to other sampling methods and is better at finding the most relevant kinetic traps in the landscape. Thus, it produces a more representative coarse graining of the landscape, which is well suited to subsequently compute better approximations of RNA folding kinetics.

**Availability:** https://github.com/ViennaRNA/RNAxplorer/

## 1 Introduction

Over the past two decades, our understanding of the roles and functions of RNAs has fundamentally changed. With the advent of next-generation sequencing a plethora of non-coding RNAs were discovered, along with specific expression patterns that support a diversity of functions within cellular compartments and molecular mechanisms (Djebali *et al.*, 2012; ENCODE Project Consortium, 2012). Accordingly, genome-wide bioinformatics studies (Eddy, 1999; Saito *et al.*, 2009) have confirmed the dense population of the intergenic space with transcripts, and comparative genomics approaches have revealed evolutionary conservation of structured ncRNAs. Even protein coding mRNAs often rely on specific structural arrangements to control their own splicing, transcription, translation, or degradation, where structure elements often serve as recognition sites for binding partners such as proteins. Modeling the structure(s) of RNA is therefore an important step towards understanding their function.

At the secondary structure level, efficient dynamic programming (DP) algorithms enable the computation of various RNA structural properties at thermodynamic equilibrium. Software suits such as `RNAstructure` (Reuter and Mathews, 2010), `UNAFold` (Markham and Zuker, 2008), or the `ViennaRNA` package (Lorenz *et al.*, 2011), enable the computation of minimum free energy (MFE), base pairing probabilities, consensus structures, RNA-RNA interactions and beyond using the Turner nearest neighbor model (Turner and Mathews, 2009).

However, RNA folding is a dynamic process that already starts during transcription. While an RNA molecule tends to adopt a stable structural conformation, i.e. one that decreases its free energy, along the way it may be trapped in local minima. Depending on the height of (energy) barriers to escape such local minima, an RNA may only explore a negligible fraction of its conformation space, and never reach its ground state within its life time. Concrete instances of kinetics, where the thermodynamic ground state is not the final state, notoriously include RNAs whose function is mediated by co-transcriptional folding, e.g. transcriptional riboswitches. Examples of such riboswitches are discussed in Breaker (2012) and most of our benchmark sequences (see Table S1) belong to this class. Since experimental methods are limited in the number of alternative structures they can verify, computational models are essential for studying riboswitches in detail. For instance, experimentally derived NMR structures (Helmling *et al.*, 2017) have been used to model concentration-dependent metabolite binding/unbinding kinetics (Wolfinger *et al.*, 2018). Furthermore, kinetic methods are invoked in rational design of artificial riboswitches (Günzel *et al.*, 2020). Folding dynamics can also be modelled on the level of tertiary structures, e.g. for G-Quadruplexes (Stadlbauer *et al.*, 2016). Due to its computational complexity, however, only small substructures can be analyzed on a coarse grained level.

A general framework for studying kinetics relies on an abstraction of the folding process as a Continuous-Time Markov Chain (CTMC) over a discrete conformational space. Properties of the CTMC can be derived from stochastic simulations of single trajectories within the folding landscape (Flamm *et al.*, 2000). However, many trajectories are then needed to estimate population densities, i.e. the probabilities/concentrations associated with most relevant conformations, hindering the kinetics analysis for RNAs beyond modest lengths. For these reasons, recent popular methods rely on a coarse-graining of the folding landscape, in which a subset of representative conformations is first identified, followed by the numerical resolution of the differential equation describing the time-resolved evolution of the population densities. Figure 1 illustrates the general principle of such a prediction workflow. The choice of a suitable coarse-graining is critical in order to allow for the omission of large parts of the conformational space, while at the same time maintaining key states in the RNA landscape for subsequent accurate approximation of RNA folding kinetics. Available approaches for coarse-graining include flooding strategies (Entzian and Raden, 2019; Wolfinger *et al.*, 2004), whose enumerative nature makes them unsuitable for RNAs beyond 100 nt. For longer RNAs, methods combining sampling with a reconstruction of the CTMC, such as the Basin Hopping Graph (Kucharík *et al.*, 2014), currently represent the only realistic option.

**Figure 1:**
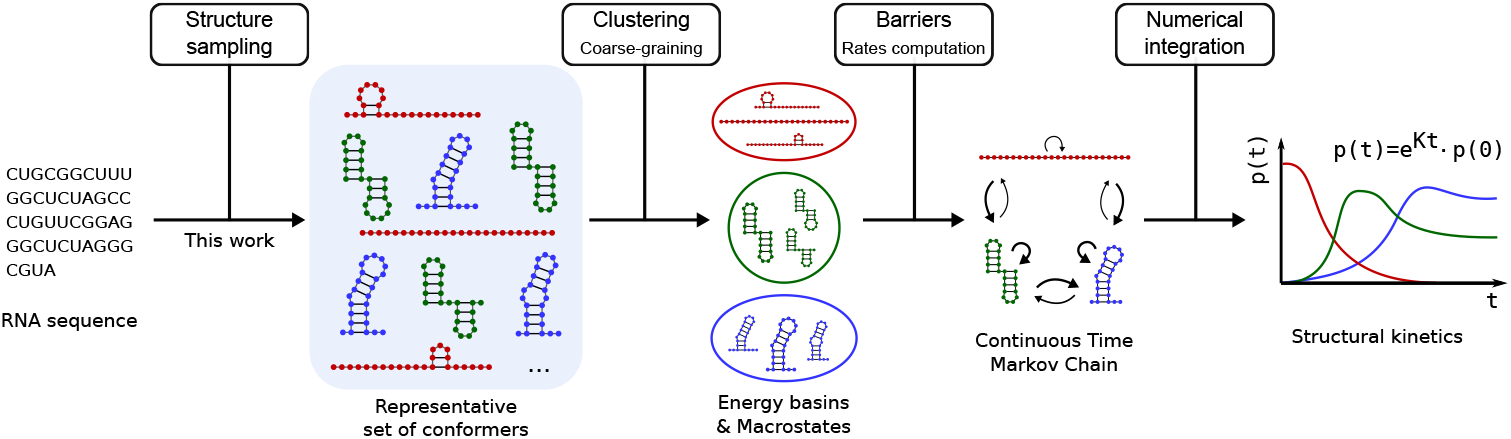
Typical workflows for RNA folding kinetics start with a sequence, for which a representative subset is sampled. This is followed by a coarse graining, rate computation and the final kinetics computation as a Markov Process.

To identify important (meta) stable secondary structures within folding landscapes, the dominant approach usually resorts to structure sampling followed by a clustering step, as introduced by Ding *et al.* (2005). However, classified DP approaches have been proven useful to yield structure representatives from partitions of the ensemble that share a common feature, for instance their abstract shape (Giegerich *et al.*, 2004) or their base-pair distance to one or two reference structures (Freyhult *et al.*, 2007; Lorenz *et al.*, 2009). Other DP algorithms reduce the state space *ab initio* to draw (random) samples that constitute locally optimal structures, i.e. where no structural neighbor has lower free energy (Kucharík *et al.*, 2014; Li and Zhang, 2011; Lorenz and Clote, 2011; Michálik *et al.*, 2017).

However, the accuracy of virtually all the aforementioned methods is hindered by a strong bias towards low-free energy structures. This situation leaves such methods to overlook important regions of the folding landscapes, or induces unreasonable computational costs due to precomputations (Michálik *et al.*, 2017), lack of diversity, forcing further rounds of sampling (Kucharík *et al.*, 2014), or the downstream reconstruction of the coarse-grained CTMC model. Indeed, the clustering of structures, and computation of (pairwise) transition rates between the structures are the computationally most demanding steps. Computing such pairwise transition rates requires approximating the energy barrier between two secondary structures, a NP-hard problem even under simplistic assumptions (Maňuch *et al.*, 2009). Consequently, the structure sampling step is the most crucial, as a good balance between the size of the sample set and the coverage of important parts of the energy landscape are required.

In this work, we present a novel method to construct accurate approximations of kinetics landscapes. To this end, we iteratively utilize an efficient DP algorithm to compute the partition function (McCaskill, 1990) including pseudo-energies (Lorenz *et al.*, 2016), subsequently draw random samples using stochastic backtracking (Ding and Lawrence, 2003) and, iteratively, refine guiding potentials to (dis-)favor particular substructures, similar to the local elevation ideas introduced in *metadynamics* (Huber *et al.*, 1994). In the context of RNA secondary structures guiding potentials have previously been used in path finding (Dotu *et al.*, 2010). Our strategy provides a fast and effective means to discover local minima that may be far away from the ground state in terms of free energy but represent important landmarks of the energy landscape due to their impact on folding dynamics.

## 2 Methods

Formally, given an RNA sequence *σ* of length *n*, a secondary structure *s*(*σ*) = {(*i, j*) | (*σ*[*i*]*, σ*[*j*]) ∈ *BP*)} is a set of base pairs (*i, j*) compatible with *σ*. Interacting nucleotides are usually restricted to the canonical Watson-Crick pairs (A, U) and (G, C) and the Wobble pair (G, U), i.e. *BP* = {(A, U), (U, A), (G, C), (C, G), (G, U), (U, G)}. The generally accepted definition of secondary structures also excludes pseudo-knots and assumes a minimum of three unpaired bases between any two pairing bases due to sterical reasons. A detailed definition is given in Supp. Sec. 1.1.

The ensemble of all secondary structures compatible with an RNA *σ* defines its conformation space Ω (*σ*) = {*s*(*σ*)}. Note, that in the following, we always assume a fixed sequence *σ* and will therefore only use Ω instead of Ω(*σ*) for the sake of convenience. In conjunction with (i) a move set 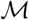 that specifies elementary transitions to transform one structure *s_i_* into one of its neighbors *s_j_*, and (ii) the energy function 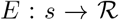 that assigns each structure *s* ∈ Ω a real numbered value, one obtains the notion of the energy landscape 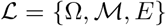. Over the past decades, different move sets 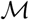 have been used (Flamm *et al.*, 2000; Xayaphoummine *et al.*, 2003), mostly to restrict the size of their induced neighborhood. The most commonly utilized move set is the difference of exactly one base pair between neighboring structures.

Local minima are defined via steepest descent trajectories *γ^∞^*(*s*) of subsequent single base pair moves. These trajectories are called gradient walks and always end in a local minimum. Structures for which a gradient walk ends in the same local minimum belong to the same gradient basin of attraction 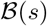. Performing gradient walks for all structures results in a unique partitioning of the state space. This is often used as a most natural coarse graining in RNA folding kinetics simulations (Wolfinger *et al.*, 2004). Definitions of gradient walks can slightly differ in resolving ambiguity. We refer to the definition used by Entzian and Raden (2019), which employs lexico-graphical order to break ties between structures with equal energy, such that the mapping of a structure to its basin representative structure becomes unique.

Moreover, gradient basins and the minimal saddle points connecting them can be used to conveniently visualize and compare high-dimensional energy landscapes as barrier trees or disconnectivity graphs (DG) (Becker and Karplus, 1997; Flamm *et al.*, 2002). However, computing the barrier tree for a particular RNA sequence typically relies on exhaustive enumeration of Ω which becomes impractical for sequence lengths of about 100 *nt* or longer, as Ω grows exponentially with the length (Waterman, 1978).

### Equilibrium ensemble properties

Most RNA secondary structure prediction methods borrow a key concept of statistical mechanics, namely that structures *s* in thermodynamic equilibrium are Boltzmann distributed, hence *p*(*s*) α exp(−*βE*(*s*)) with *β* := 1*/kT* for *k* the Boltzmann constant and *T* the temperature. For a particular RNA sequence this immediately suggests an obvious structure representative: the one with minimal free energy (MFE), i.e. *s*_MFE_ = arg min_*s*∈Ω_ *E*(*s*) since it has the highest probability among all other structures of the conformation space. Efficient DP algorithms exist that compute *s*_MFE_ in 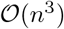 time and 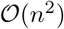 memory for sequences of length *n* (Zuker and Stiegler, 1981). A small change in this DP concept leads to an efficient method to compute the partition function *Z* = Σ_*s*∈Ω_ exp(−*βE*(*s*)), with the same asymptotic complexities (McCaskill, 1990). Using *Z* many thermodynamic equilibrium properties can be derived, e.g. probabilities

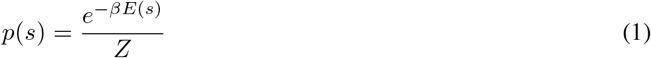

for any structure *s* or *p_ij_* = Σ_*s*|∈(*i,j*)∈*s*_ *p*(*s*) for base pairs (*i, j*). The DP algorithm to compute *Z* can also be adapted to perform Boltzmann sampling, i.e. to draw structures *s* randomly from the ensemble according to their probability *p*(*s*). This can be regarded as (random) backtracing in the DP matrices with worst case time complexity of 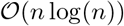 per sample (Ding and Lawrence, 2003; Ponty, 2008).

### 2.1 The RNAxplorer Method

The `RNAxplorer` method approximates RNA energy landscapes using an iterative scheme which samples random structures using guiding potentials. To mitigate oversampling, we introduce a focused approach based on (directed) guiding potentials, i.e. pseudo-energy terms that supplement the free-energy, and steer the sampling away from a (set of) structure(s). Pseudo energy terms are accumulated after each iteration to avoid a concentration of samples within low free-energy basins, thus ensuring maximal coverage of the landscape. This allows a finer level of control over the redistribution of the emission probabilities than previous alternatives, such as the temperature elevation method introduced by Kucharík *et al.* (2014) (see Supp. Mat. 1.3).

#### Sampling with base pairs-associated guiding potentials

Given a pseudo energy *E*_Ψ_(*s*), our sampling procedure considers a pseudo-energy function 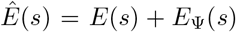 where *E*(*s*) is the classic Turner free-energy and *E*_Ψ_(*s*) is a guiding potential defined below. Our goal is then to sample from the distribution

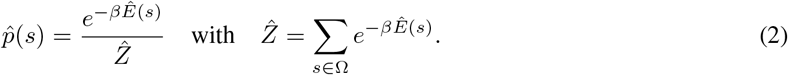

Boltzmann sampling requires the precomputation of the (pseudo) partition function 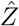, not through exhaustive summation due to the combinatorial explosion of Ω, but rather by using a recursive DP scheme. Thus, in order to benefit from efficient algorithms, we restrict our attention to guiding potentials *E*_Ψ_ such that, for any structure *s*, *E*_Ψ_(*s*) can be written as a sum of contributions associated with derivations of the underlying folding grammar. Sampling under such guiding potentials is generically supported by the soft constraints framework introduced by Lorenz *et al.* (2016).

In particular, let us consider simple, base pairs-associated potentials, which can be decomposed into energy terms *E^i,j^* ∈ ℝ, each associated to a base pair (*i, j*). For any structure *s*, one has

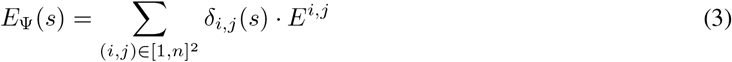

where *δ_i,j_* ∈ (*s*) is the indicator function, taking value 1 if (*i, j*) ∈ *s* and 0 otherwise. Despite their simplicity, *E*_Ψ_ terms can be used to steer the sampling towards/away from one or several reference structure(s).

For instance, the base pairs of a reference structure *s′* can be individually penalized/promoted by setting *E^i,j^* = *δ_i,j_*(*s′*) · *α*, for some arbitrary real *α*, leading to *E*_Ψ_(*s*) = |*s* ⋂ *s′*| · *α*. Setting *α* < 0 will decrease the expected distance between sampled structures to *s′*, while *α >* 0 will increase it. Note that more elaborate guiding potentials can be supported, e.g. through variations of the energy values *α* and/or a combination of using individual base pairs and structures as targets (see Supp. Sec. 2.2 and 2.3).

#### Defining guiding potentials to avoid recurrent structures

In order to steer sampling away from a given structure *s′* that has already been sampled repeatedly, we consider a guiding potential

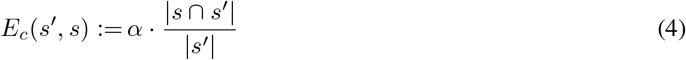

that for each structure *s* ∈ Ω adds individual pseudo-energy penalties, depending on the number of base pairs *s* shares with *s′*.

Moderate penalties arise if the weight factor *α* is chosen close to thermal fluctuations, which is why our method defaults to *α* = *kT* unless stated otherwise. Such potentials could also be used to attract subsequent sampling towards a region of interest within the kinetics landscape, by simply changing the sign of *α*.

Finally, guiding potentials can be modified to capture the base pair distance between *s′* and *s*, i.e. the minimum number of base pairs to insert/remove to transform *s′* structure into *s* (see Supp. Eqn. (9) and (12)). This alternative definition yields comparable, yet slightly inferior results as shown in the Supp. Mat. 2.1.

#### Overall iterative strategy

For each structure *s*, the pseudo-energy *E*_Ψ_(*s*) is initially set to 0, and incrementally updated to accumulate contributions from the dominant structures encountered over the course of sampling.

At each round *m*, a multiset 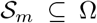 of structures is sampled from a (distorted) Boltzmann distribution. Through gradient descent, each structure 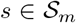 is mapped to its local minimum *γ^∞^*(*s*), used as a representative ŝ for its energy basin. The resulting set of local minima is then analyzed to identify the most over-represented structure, denoted as

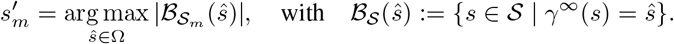

In other words, 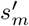 is the local minimum that attracts the most samples in 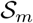. In the (unlikely) case of ties, one of the most highly represented structure is chosen arbitrarily and returned.

The method then updates the pseudo-energy term *E*_Ψ_ for the next iteration, based on the structural features of 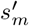, by setting:

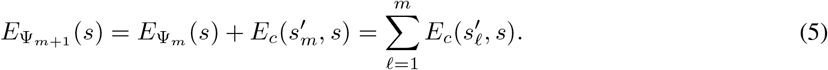

This can be expressed at level of individual base pairs (*i, j*) by setting

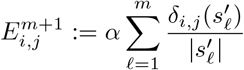

where 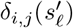 denotes the presence (1) or absence (0) of (*i, j*) in 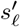. Then, any structure *s* inherits a total pseudo-potential of:

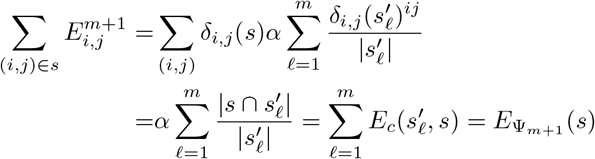

in which one recognizes the intended guiding potential after *m* updates.

The total number of iterations 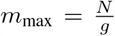 is governed by the fraction of the requested sample size *N* and a user-adjustable granularity *g* that determines the number of samples drawn at once in each round. Unless stated otherwise, we use default values *N* = 10^5^ and *g* = 100.

Additionally, we use a strategy which determines whether *E*_Ψ_ needs an update after each iteration. This avoids unnecessary recomputation of the rather costly partition function (see Sec. 2.1). The general idea is that the depth of a sampling is sufficient if collisions pervasively occur, i.e. most structures are observed multiple times (Sahoo and Albrecht, 2012). Therefore we compare the set 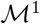 of local minima that are observed only once over all iterations, against 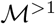, those observed multiple times. Our algorithm then only updates *E*_Ψ_ if the ratio 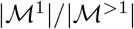 does not exceed a saturation threshold *μ*, set by default to *μ* = 0.1 by analogy to Kucharík *et al.* (2014).

### 2.2 Quality Assessment

Whether or not a landscape 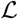 is adequately approximated by a set of structures strongly depends on the requirements of downstream analysis and thus is difficult to generalize. In the following we discuss the use of general measures for structure diversity, distance class based diversity measures and the coverage of basins and energy barriers in 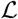. The approximated shape of 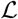 is important for the overall dynamical behavior of subsequent folding simulations. We, therefore, also analyze sample sets for the presence of certain key structures.

#### General measures

Typical measures that express the diversity of a set of structures are the number of unique local minima, the Density of States (DOS) (Cupal *et al.*, 1997) and the weighted mean base pair distance. Density of States is simply the number of structures per energy level. The weighted mean pairwise distance is defined as the sum over all base pair distances between structures *s* and *t* multiplied by their probabilities *p*(*s*) and *p*(*t*). For details we refer to Supp. Mat. 5.1.

#### Distance classes

The partitioning of Ω into distance classes with respect to one or many reference structures leads to a projection of the high-dimensional state space. Such a projection still captures the structural diversity of the ensemble, but can now be easily visualized and characterized. Following the lines of Lorenz *et al.* (2009), with two fixed references ŝ_1_, ŝ_2_, each structure *s* ∈ Ω is assigned to its corresponding class 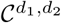 where *d*_1_ = *d*_BP_(s, ŝ_1_) and *d*_2_ = *d*_BP_(s, ŝ_2_). Each class can then be represented by

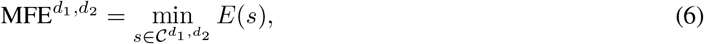

or the corresponding ensemble free energy

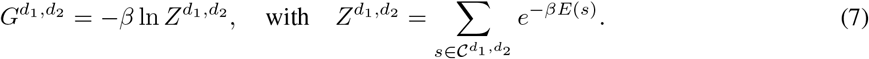

Finally, the resulting projections can be conveniently visualized in Cartesian coordinates and dimensions *d*_1_ and *d*_2_, for instance in the form of a heat map (see Fig. 2 or Fig. 5).

**Figure 2:**
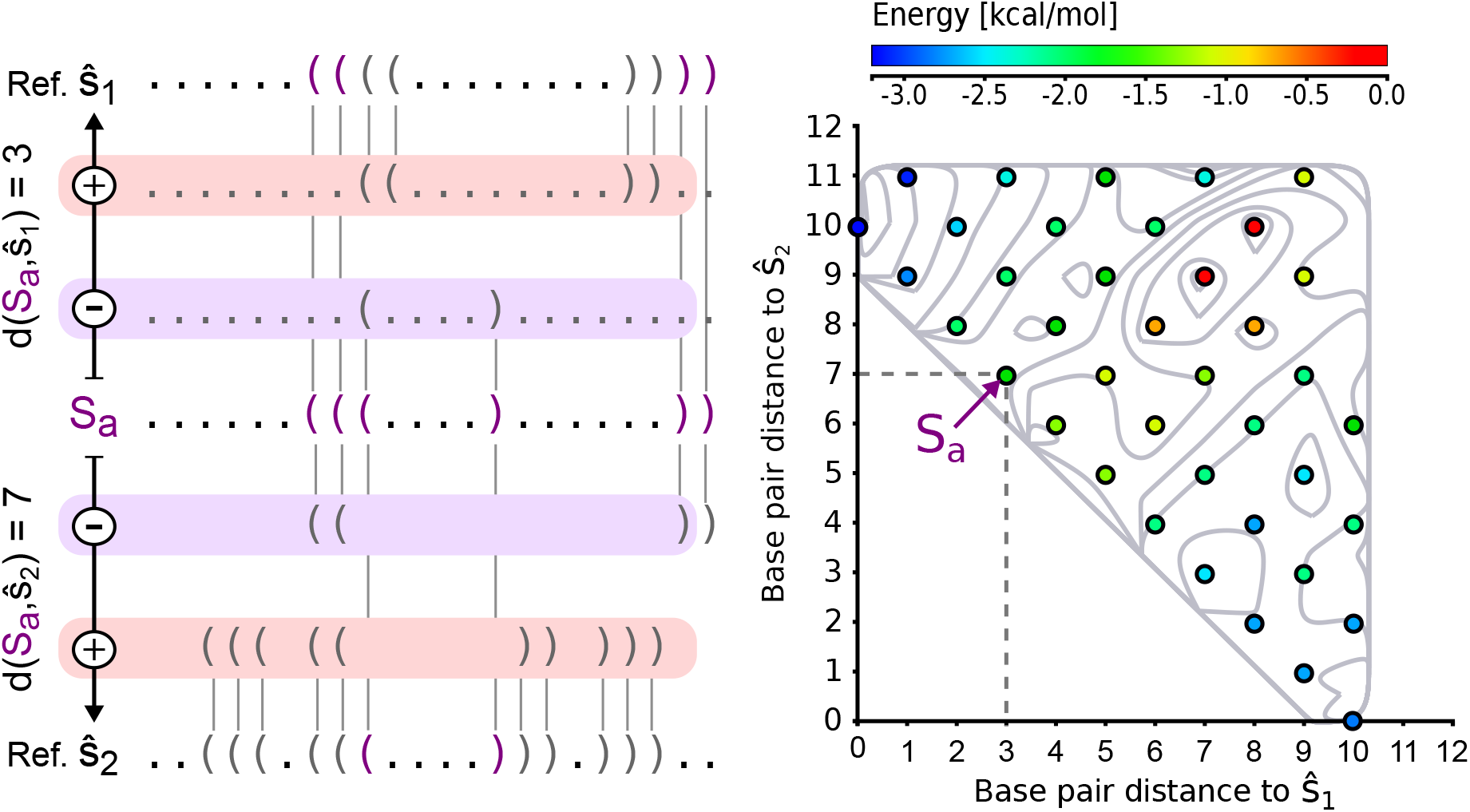
Toy example of a 2D projection. Each colored circle corresponds to a valid base pair distance to both reference structures. The color corresponds to the energy of the MFE structure representative of this spot. The background consists of isolines which are created from an interpolation of the free energy values that correspond to the MFE structures of each circle. Structure *S_a_* has base pair distance 3 to the reference ŝ_1_, because one base pair has to be removed and two added in order to change ŝ_1_ into *S_a_*. The distance to ŝ_2_ is 7 because two base pairs must be removed and 5 inserted to change *S_a_* into ŝ_2_.

As a proxy of diversity, we count the number of distance classes 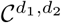 that are adequately covered by a sample set 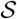. We assume a class to be covered, if any sampled structure *s* mapped to 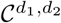 is within an energy margin *ϑ* around 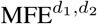 of the full ensemble, i.e. 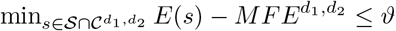, cf. Supp. Sec. 5.5.

#### Energy barriers

Comparing folding simulations produced by different tools can be challenging, because the inherently different coarse graining of each sampling method results in different representations of both fast fluctuations at the beginning of the simulation and slow folding components close to thermodynamic equilibrium. To assess the quality of our sampling strategies within the RNA folding kinetics workflow (Fig. 1) we need to evaluate our samples in terms of providing a basis for calculating folding rates. For this, samples must not only cover the lowest energy states of 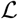, but also refolding events with large energy barriers, i.e. those associated with slow rates that effectively determine the long time behavior of folding dynamics (Becker and Karplus, 1997; Flamm *et al.*, 2002). For that purpose, structures of a sample set can be mapped into a barrier tree representation of the full ensemble. We then compute the fraction of leaves covered by, and the highest energy barriers associated with the structures within each sample set. For details, we refer to Supp. Sec. 5.3.

### 2.3 Implementation

From the implementation perspective, we use the constraint framework of the `ViennaRNA` Package that allows us to specify guiding potentials *E*_Ψ_ as separate functions which are then integrated into the prediction algorithms (Lorenz *et al.*, 2016). This allows us to dynamically adapt *E*_Ψ_ without the need to re-implement the computation of 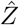 and the subsequent Boltzmann sampling. We implemented the novel guiding potential-based sampling approach described in Section 2.1 using the programming language C, as part of the executable program `RNAxplorer`. For the iterative sampling method, the user can choose between two guiding potentials, the base pair associated potential, described in section 2.1, or an alternative based on base pair distance, described in Supp. Sec. 2.1. To deviate from the default parameters, the granularity *g*, the total sample size *N*, the saturation threshold *μ*, as well as the weighting factor *α* are user-adjustable. The `RNAxplorer` program offers additional guiding potential-based structure sampling modes, e.g. one that explicitly penalizes or rewards a pre-defined set of reference structures and structures in its vicinity. Furthermore, it implements different heuristics to compute (optimal) transition paths to eventually determine saddle points required to assess transition rates, and finally, provides gradient walk methods to coarsen the sampled state space. The program also comes with a Python script that enables hierarchical clustering of secondary structures, see Supp. Sec. 2.3.

## 3 Results

In the following we assess the quality and applicability of our novel sampling method by comparing its results against other widely-used RNA secondary structure sampling methods. For that purpose, we first collected a set of 9 benchmark sequences for which landscape approximations were made. They have a minimum length of 110 nt, a maximum length of 233 nt and a median length of 130 nt. The exact sequences can be found in Table S1 of the supplementary material. The methods and tools we compare our approach against are (i) *uniform sampling* with `RNAsubopt` achieved using Boltzmann sampling at extremely high temperatures (10^6^ °C), (ii) regular Boltzmann sampling with `RNAsubopt` (*-p* command line option), (iii) Boltzmann sampling of locally optimal structures with RNAlocopt (Lorenz and Clote, 2011), (iv) non-redundant sampling of saturated structures with RNANR (Michálik *et al.*, 2017), (v) the temperature elevation scheme of RNAlocmin (Kucharík *et al.*, 2014), and (vi) a set of locally stable structures, generated by RNASLOpt (Li and Zhang, 2011). Note, that RNASLOpt differs from all the others in that it is deterministic and always exhaustively enumerates locally optimal structures (LOpts) in a pre-defined energy band above the MFE. The width *ɛ* of this band can be specified in discrete steps of kcal/mol or percentages. This, unfortunately, prohibits one to explicitly set the number of output structures in advance. Therefore, in some of the analysis below, we either determined the minimal width *ɛ* that results in at least the number of required samples in a pre-processing step, or we simply omit its use altogether. All programs were used with default parameters unless stated otherwise.

### 3.1 Time and Memory Consumption

First, we prepared a set of artificially generated random sequences with equal probabilities for each of the four RNA nucleotides to assess the runtime and memory requirements for all programs in our comparison. To that end, we generated 10 sequences with lengths of 50–300 nt in steps of 50 nt. For each of the resulting 60 sequences the 6 different tools were instructed to (randomly) draw 1, 000 structures from the respective ensembles. For the iterative methods implemented in `RNAlocmin` and `RNAxplorer`, the number of iterations was set to 100. Programs were compiled with `GCC` 8.2.1 and all computations were performed on a `Linux` workstation with Intel^®^ Core™ i7-7700K CPU running at 4.20 GHz and 32 GB of RAM.

As expected, the standard Boltzmann sampling strategies of `RNAsubopt` with default parameters as well as *uniform sampling* were the fastest methods tested (Fig. 3) and required the least amount of memory. The next best tool in terms of both, runtime and memory requirements, is our new heuristic `RNAxplorer`, followed by `RNAlocopt` and `RNAlocmin`. While runtimes of `RNAxplorer` and `RNAsubopt` are within the same order of magnitude, `RNAlocopt` and `RNAlocmin` are by two orders of magnitude slower. The exponential runtime asymptotics of `RNANR` and `RNASLOpt` render them the slowest for longer sequences. Note, that we were not able to produce results (within 3 days) for sequence longer than 150 nt (`RNANR`) and 200 nt (`RNASLOpt`) due to the limited memory of our testing machine. However, for shorter RNA sequences up to 150 nt, these two programs are still faster than `RNAlocmin` (Fig. 3). Further runtime and memory benchmarking results are available in Supp. Sec. 4.

**Figure 3:**
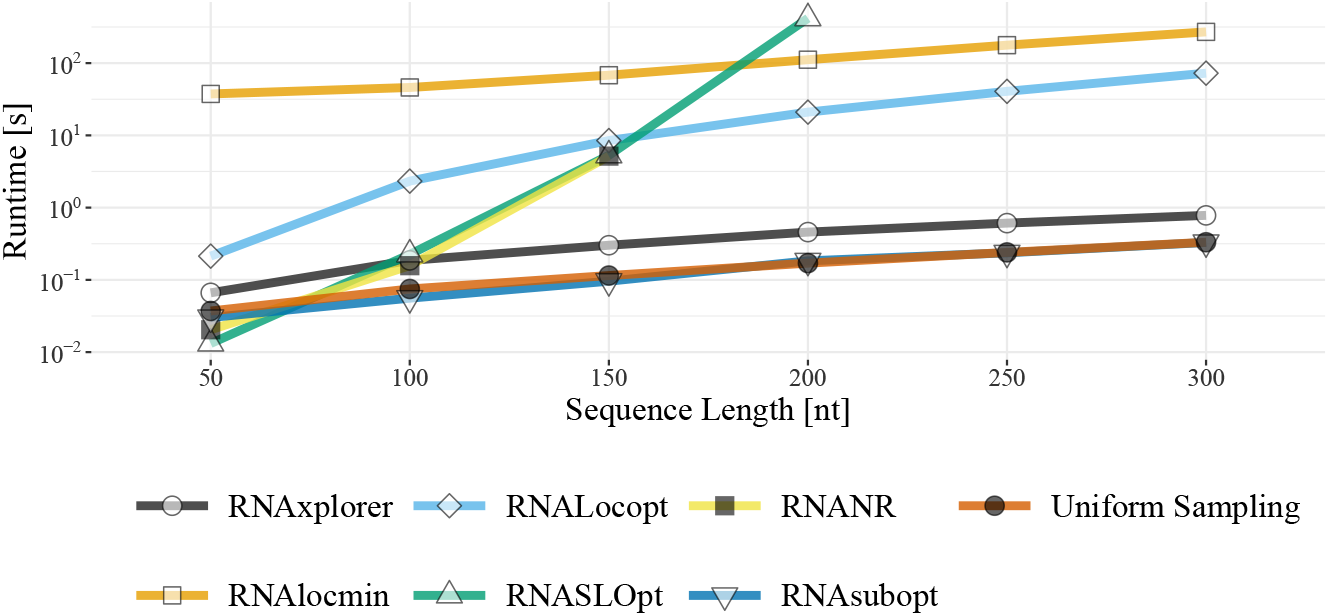
Runtime comparison. Runtimes observed for a sample of 1,000 structures for RNAs with lengths from 50 to 300 nucleotides, averaged over 10 randomly generated sequences. For `RNASLOpt`, we precomputed a *ɛ* value in order to obtain at least 1,000 structures. For `RNAxplorer` and `RNAlocmin` the number of iterations was set to 100.

### 3.2 Structure Sample Diversity

For each method we assessed sampling quality in terms of diversity of structures obtained. First we calculated standard measures to provide an overall description of the samples produced. Sampling redundancy in terms of (i) number of unique local minima and (ii) mean base pair distance both turn out favorable for `RNAxplorer` and `RNANR` (see Supp. Sec. 5.1 and Supp. Fig. S5). The energy spectrum of the samples expressed as Density of States shows that most methods are prone to over-sample the low free energy regime, while `RNAxplorer` also captures structures at higher energy levels, e.g. structures around the meta-stable state of SV-11 appear as second peak (Supp. Fig. S6). However, these results are not sufficient to evaluate the methods regarding their suitability within the folding kinetics workflow. In the following we therefore focus on a newly developed measure to investigate the spatial resolution of the sample sets based on distance classes.

#### Coverage of distance classes

Our main question was whether (i) the samples spread over a large number of representative structures with fundamentally different base pair patterns, or (ii) the samples mainly reflect representatives of structurally similar clusters. For that purpose, we use distance classes 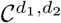 (cf. Fig. 2), where we partitioned the sample sets according to their distance to (i) the MFE structure and (ii) the most stable structure that does not share any base pair with the MFE structure. Note, that the latter can be obtained from a constrained MFE prediction where all base pairs of the actual MFE structure are prohibited. For each class we computed the MFE and ensemble free energy to compare them against exact values as computed with `RNA2Dfold` (Lorenz *et al.*, 2009).

Such projections into lower dimensions provide easy to assess visual impressions of the sample diversity, as shown in Figure 5. However, here we use them to count how many 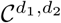 were covered by the different sampling methods. To alleviate the impact of randomness during the sample generating process, we averaged the results for each experiment over 10 independent runs. Figure 4 summarizes the results over all benchmark sequences as a function of sample size and two thresholds *ϑ*_1_ = 0 kcal/mol and *ϑ*_2_ = 5 kcal/mol.

**Figure 4:**
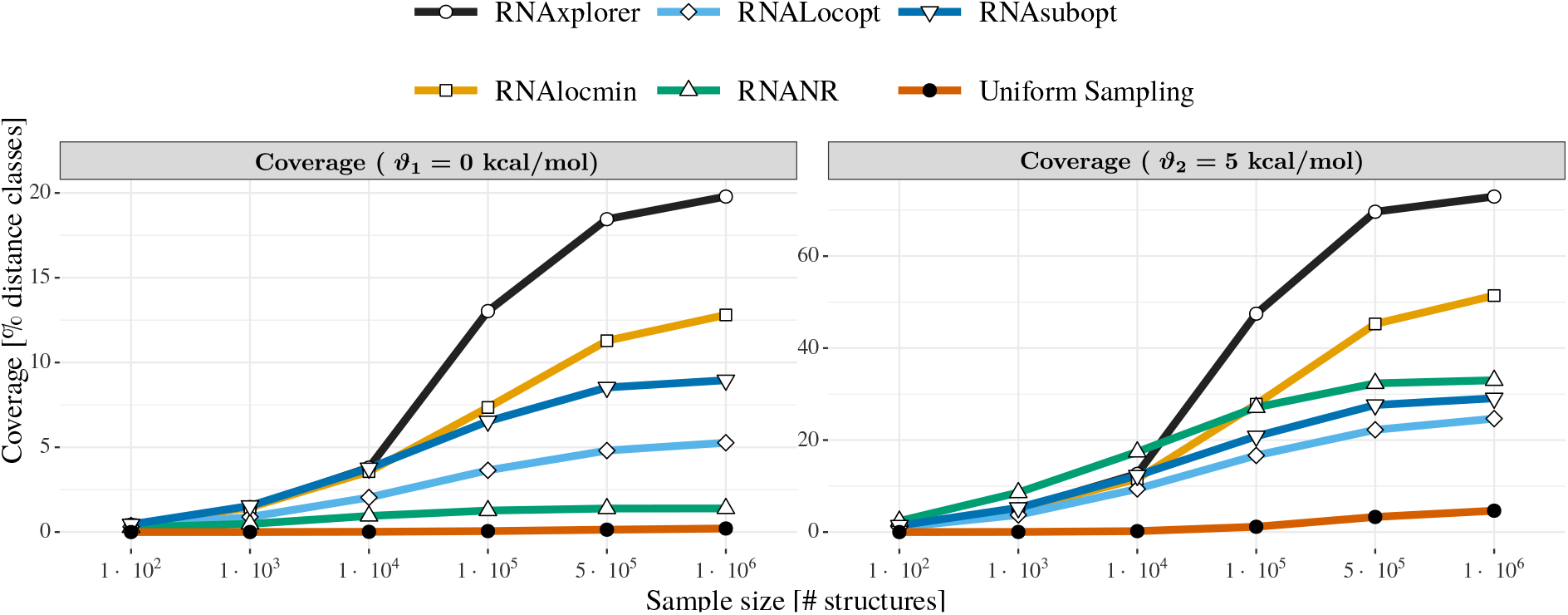
Distance class coverage as a function of sample size. Shown are the fractions of distance classes 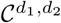 covered by at least one local minimum. The local minima are derived from the sample set and have to be energetically close to the respective 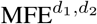. The data averages over all 9 benchmark sequences, 10 independent runs per tool and margins *ϑ*_1_ = 0 kcal/mol (left plot), and *ϑ*_2_ = 5 kcal/mol (right plot).

**Figure 5:**
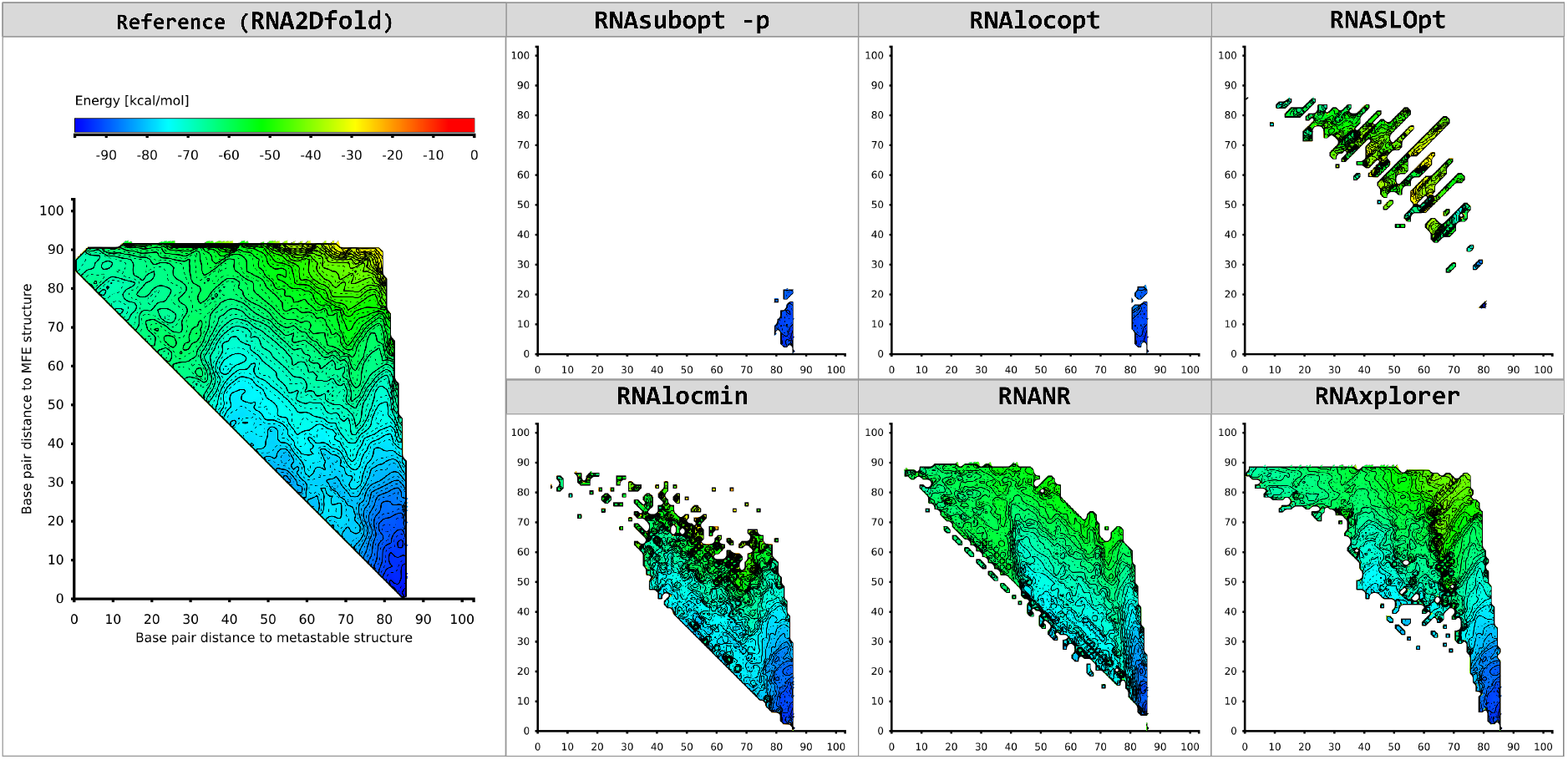
2D projections of local minima as obtained from different methods for the riboswitch SV-11 Q beta replicase template (Biebricher and Luce, 1992). Reference structures for the projection are the MFE and metastable structure. The left most column depicts the *ground truth* as computed by `RNA2Dfold`, chosen here as a reference for comparison. The remaining panels show the results for Boltzmann sampling (`RNAsubopt` −p), local optima sampling (`RNAlocopt`), `RNASLOpt`, variable temperature sampling (`RNAlocmin`), non-redundant sampling (`RNANR`), and repellant sampling (`RNAxplorer`), which required 6.75s, 21.81s, 115.99s, 487.73s, 4285.81s, and 27.87s to produce the sample sets, respectively. The sample size for each tool is 10^6^ (except for `RNASLOpt` which always yields less structures even with exhaustive enumeration, i.e. artificially high *ɛ*).

`RNAxplorer` clearly outperforms the other methods even for small sample sizes. With increasing sample size the coverage quickly rises and is always higher compared to the other methods. Only for `RNAlocmin` the coverage rises similarly fast with increasing sample size. The next best tools are `RNAsubopt` and `RNAlocopt` (*ϑ*_1_) and `RNANR` (*ϑ*_2_). As expected, *uniform sampling* covers just a tiny, almost constant fraction even for very large sample sizes of 10^6^ structures. For `RNANR` the diversity is very sequence dependent which is depicted in Figure S16. Since `RNANR` could not be applied to 3 of the 9 benchmark sequences (SAM riboswitch of `metE`, lysine riboswitch of `lysC` and TPP riboswitch of `thiamine` gene) due to its demanding memory requirements (more than 200GB), the average for this tool as shown in Figure 4 only consists of the remaining 6 sequences. Results for the individual benchmark sequences can be found in Figure S16. The analog measure based on partition functions is shown in Figure S18 for individual sequences and in Figure S19 as average over all sequences. For more details on the coverage measure see Supp. Sec. 5.5.

### 3.3 Suitability for RNA Folding Kinetics

Using the `barriers` program (Flamm *et al.*, 2002) we generated barrier trees for our benchmark set of random sequences using exhaustive structure enumeration up to 15 kcal/mol above the MFE with `RNAsubopt`. Coarse graining of the barrier tree was set to a minimal energy barrier of 3 kcal/mol between neighboring basins. We then mapped the local minima generated by each sampling method into the respective barrier trees to determine how many of the 100 largest energy barriers could be found based on the samples. The results were further averaged over 10 rounds of sampling to alleviate the impact of randomness in the sample sets.

As shown in Table 1, for 100 nt long sequences all tools already find a large amount of the highest energy barriers even for small sample sizes such as 10^3^. At the same time, the number of recovered basins is as low as 1 — 2%. `RNANR` in general recovers more basins than the other tools for sequence lengths of 70 nt or longer. For sample sizes of 10^5^ structures, the tools `RNAxplorer`, `RNAlocmin`, `RNAlocopt`, and `RNAsubopt` perform equally good in finding the highest energy barriers. In contrast, both `RNAxplorer` and `RNAlocmin` stand out in the number of recovered basins with 22.25% and 15.48%, respectively, compared to less than 10% achieved by the other methods. In terms of run time, `RNAxplorer` is much faster than `RNAlocmin` with an average of just 3.72 s compared to 73.61 s. Details and remaining results for this analysis are available in Supp. Sec. 5.3.

**Table 1:**
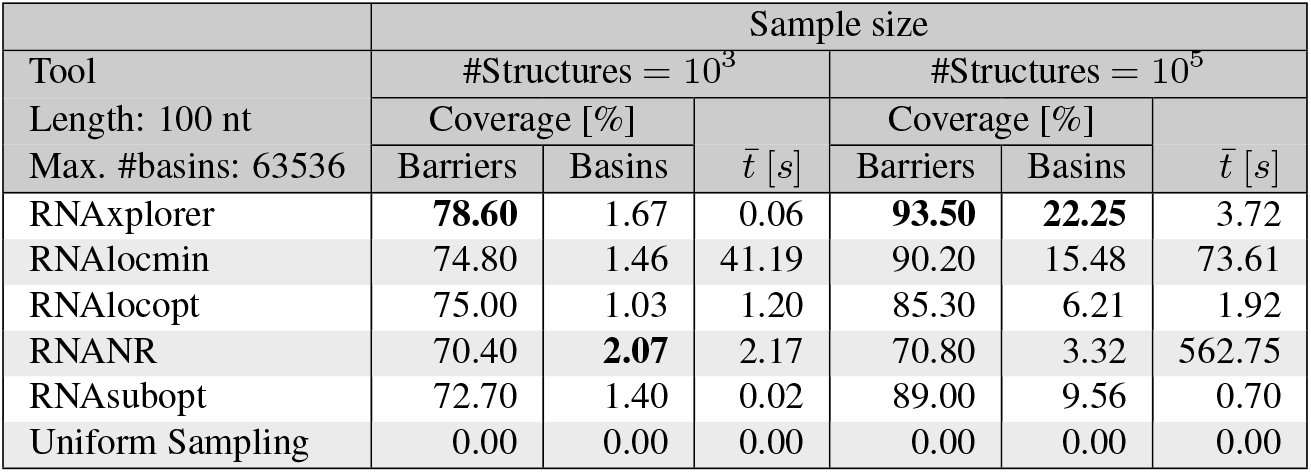
Coverage of barrier trees for ten 100 nt long sequences, using sample sizes of 10^3^ and 10^5^, respectively. Values in ‘barriers’ columns show coverage of the 100 highest saddle points associated to the deepest left and right minima in the barriers tree. For colums ‘basin’ the percentatge of total basins covered is shown. 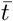 columns report average runtime in seconds.

## 4 Conclusion and Discussion

In this paper we have introduced `RNAxplorer`, a tool based on an RNA secondary structure sampling method with guiding potentials to approximate the underlying energy landscape. Its very small foot print in terms of memory and computation time requirements enables it to be applied to RNAs with sequence lengths beyond those that can be handled with other, comparable approaches. Our tool creates diverse structure samples with low as well as high free energy, that seem to nicely encompass those relevant for subsequent folding kinetics simulations. This has been shown in a benchmark analysis for biologically relevant and randomly generated RNAs using various quality measures. Thus, our novel sampling method may enable the investigation of the folding dynamics of longer RNAs than possible with state-of-the-art tools.

Efficient implementation, simple strategy and utilization of features of the `ViennaRNA Package` in general and soft constraints in particular make `RNAxplorer` one of the fastest structure sampling methods available. Memory consumption is minimal and mostly attributed to storing the list of structures obtained and the DP matrices of the partition function computations. As a consequence, unlike other tools in our benchmark, `RNAxplorer` yields representative samples within reasonable time frames even for RNAs with lengths of 300 nt or beyond.

The main contribution to the asymptotic time complexity of our new approach is the number of times new guiding potentials are added, as they each require additional 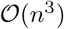 time to re-compute the partition function. For sequences of length *n* and a total number of structures *N* to sample, the upper limit on the runtime becomes 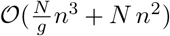. Choosing *g ≈ n* ensures that the folding and sampling part of the program have approximately equal costs, leading to a worst-case asymptotic complexity in 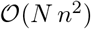. Moreover, most sampling rounds do not satisfy the saturation criterion, so a typical run of `RNAxplorer` requires much less than 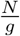 recomputations of the partition function, further reducing its practical computational demand.

While RNAxplorer has a number of tunable parameters, these parameters have default values that should work well for almost any application. Users can adjust these parameters manually, but are invited to proceed with caution. Indeed, setting the weight factor *α* to a low value, yields structures that are approximately Boltzmann distributed, and mainly populates the MFE basin. On the other hand, *α* should not be much larger than *kT* to ensure that the Boltzmann distributions of consecutive iterations have sufficient overlap.

### Coverage of distance classes

The coverage of distance classes is a combined measure which consists of important local minima and structural diversity based on 2D projections. For small sample sizes (less than 10^4^), `RNANR`, `RNAlocmin` and `RNAlocopt`, could be alternative methods with comparable quality. For larger sample sizes (10^5^ – 10^6^), `RNAxplorer` clearly outperforms the other methods. Although only at most 35% of local minima are uniquely sampled, the samples are diverse and `RNAxplorer` covers more than 70% of the projected landscapes with low energy structures, for 20% of the landscape we even identify the local minimum (Fig. 4).

Studying the topology of a landscape projection can help to choose the most efficient sampling strategy for a given problem. If, for instance, only one local minimum is present, simple Boltzmann sampling might suffice. In cases with additional metastable states, guiding potentials are the method of choice, because they steer the sampling procedure directly to structures far away of the MFE structure. This scenario is exemplified by our data for riboswitch SV-11, (Fig. 5), where only `RNAxplorer` identified the two functional states of the switch. `RNANR` produces a projection with a similar cell coverage, however, it does not find as many local minima as `RNAxplorer`, which is shown by the less intense coloring of cells compared to the reference 2D plot (Fig. 5), the lower coverage of distance classes (Supp. Fig. S16) and the DOS (Supp. Fig. S6). `RNAlocmin` could not find the metastable state, because at later stages, i.e. at higher temperatures, it turns into uniform sampling, which results in mostly high free energy structures and thus misses potentially important local minima. In the 2D projection this can be seen as separate spots in the higher energy areas (Fig. 5).

### Coverage of energy landscapes

The coverage of distance classes already indicates whether important local minima have been sampled. Using `barriers` we test for support of important transitions over high energy barriers by comparing all samples to a ground truth calculated by exhaustive enumeration. Unfortunately, `barriers` is limited to RNAs smaller than 100 nt due to time and memory constraints. For this reason we cannot use our benchmarking set of natural RNAs, but created random sequences in the range of 50 to 100 nt.

`RNAxplorer` covers the largest energy barriers much better than other tools, even for small sample sizes. Although, `RNAlocmin`, `RNAlocopt`, and `RNAsubopt` produce a comparable high number of largest barriers, they cover a much smaller fraction of basins (Table 1). RNAxplorer covers much more basins and thus provides in addition to the major transitions more detailed information on fast refolding processes. With growing sequence length, `RNAxplorer` outperforms the other methods in terms of covered barriers and basins, as well as run time. Thus, we show that `RNAxplorer` yields a very good approximation of the actual state space and is better suited for fast and efficient sampling of long sequences.

### Relationship to continuous energy landscapes

It should be noted, that the application of penalizing pseudo-energy potentials is similar to the concept of meta dynamics simulations on continuous energy landscapes (Laio and Parrinello, 2002), in particular the Local Elevation (LE) method, as used for Monte Carlo protein folding simulations (Huber *et al.*, 1994). However, for the discrete energy landscapes of RNA secondary structures, we can use efficient methods to compute the partition function and to sample from the entire Boltzmann distributed ensemble. Thus, approximations of the landscape can be directly obtained from the samples rather than from time-consuming Monte Carlo simulations. Furthermore, the RNA folding grammar does not allow for the application of Gaussian potentials as required for the LE method, but is rather limited to potentials that linearly depend on particular structural features.

### RNAxplorer in the folding kinetics workflow

To summarize our benchmark study, we have demonstrated that `RNAxplorer` is well suited to serve as sampling tool within the RNA folding kinetics workflow (see Fig. 1). It creates both diverse and low free energy structures much faster than other tools. As a result, it covers most of the basins and largest barriers which are crucial to model the long term folding behavior. Thus, the samples produced by `RNAxplorer` sufficiently capture the relevant areas of the structure space and we think that `RNAxplorer` sets a mile stone in computing the folding kinetics for RNAs of 300 nt and beyond.

