## Supplementary material for "RNAxplorer: Harnessing the Power of Guiding Potentials to Sample RNA Landscapes"

### 1 Concepts and notations

#### 1.1 RNA Secondary Structures

An RNA sequence $\sigma = x_1, \ldots, x_n$ is a string of length $n$ consisting of characters $\sigma[i] = x_i \in \Sigma$ over the RNA alphabet $\Sigma = \{A, C, G, U\}$ denoting the nucleotides of the RNA molecule from 5' to 3' end. A secondary structure $s(\sigma) = \{(i,j) \mid (\sigma[i], \sigma[j]) \in BP\}$ compatible with $\sigma$ then is a set of base pairs $(i,j)$ between two nucleotides at position $i$ and $j$. In the following, we assume a fixed sequence $\sigma$ and write $s$ instead of $s(\sigma)$ for short. Typically, the set of allowed base pairs is limited to the so-called Watson-Crick pairs $(A,U)$ and $(G,C)$, and the Wobble pair $(G,U)$, i.e. $BP = \{(\mathsf{A},\mathsf{U}),(\mathsf{U},\mathsf{A}),(\mathsf{G},\mathsf{C}),(\mathsf{C},\mathsf{G}),(\mathsf{G},\mathsf{U}),(\mathsf{U},\mathsf{G})\}$. Furthermore, any nucleotide $x_i$ is restricted to be part of at most one base pair, the minimal backbone length $|i-j|-1 \geq m$ of a base pair $(i,j)$ is limited by a constant $m = 3$ for sterical reasons, and the most common definition of a secondary structure excludes crossing basepairs, so-called pseudo knots, that arise if for any two base pairs $(i,j)$ and $(k,l)$ the crossing condition $i < k < j < l$ is fulfilled. The dot-bracket notation of a secondary structure $s$ for a sequence of length $n$ is a string of the same length that is composed of matching parentheses and dots. Each base pair $(i,j) \in s$ is represented by a *(* and *)* at positions $i$ and $j$, unpaired bases are represented by the dot character ., e.g. dot-bracket notation for $s = \{(2,10),(3,9),(4,8)\}$ of the sequence `AGGGAUACCC` is

```
.(((...)))
```

The above definition allows one to regard secondary structures as an outerplanar graph, that can be efficiently decomposed into its faces, the loops $L \in s$ of a secondary structure $s$. Physics based structure prediction and evaluation methods assign each loop a free energy contribution $E_L$ and the total free energy of a structure $s$ is estimated as $E(s) \approx \sum_L E_L$. The individual loop energies $E_L$ have been mostly derived from UV-melting experiments and mathematical modeling and follow the so-called Turner Nearest Neighbor energy model [13].

**RNA structure neighborhood** The neighborhood of a structure is the set of all structures that can be created by application of a single move from an underlying move set. In this study, we use the two symmetrical moves of (i) removing an existing and (ii) closing a new base pair. An example of the induced neighborhood for a particular structure is shown in figure S1.

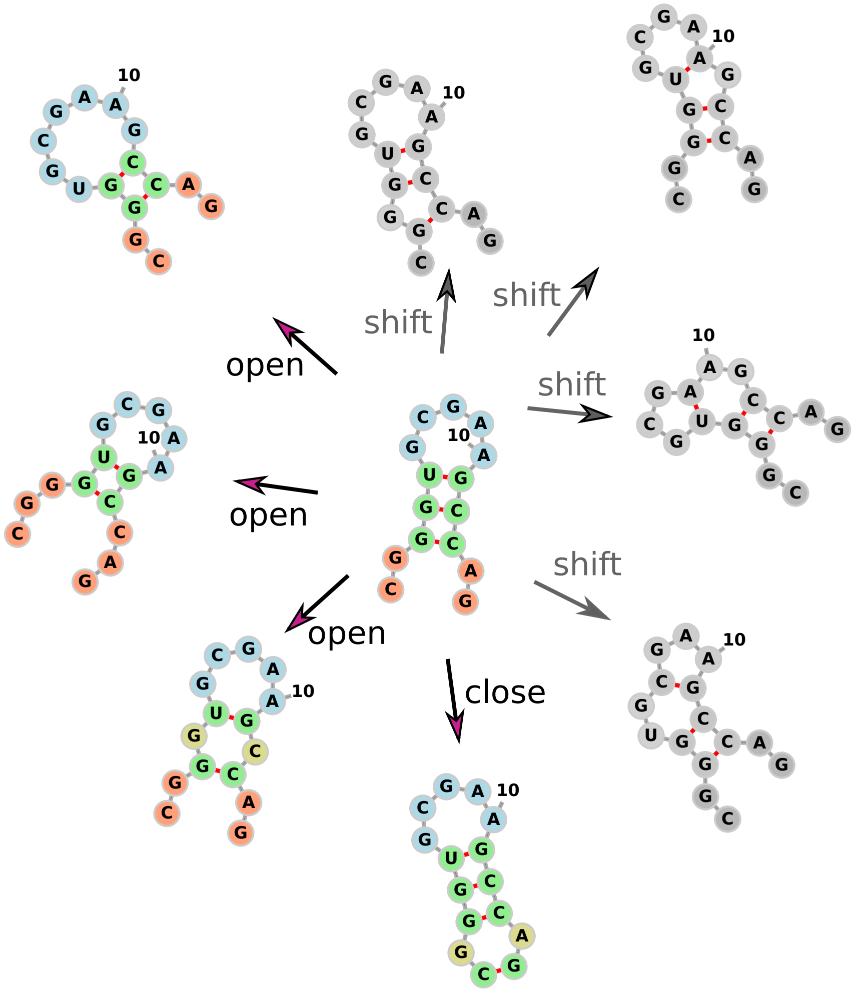

Figure S1: **Neighborhood of a secondary structure.** Shown are all direct structural neighbors of the structure `..(((.....)))..` for the sequence `CGGGUGCGAAGCCAG`. Here we apply a move set that consists of formation of a new base pair (close), removal of an existing base pair (open), and shifting an existing base pair (shift), i.e. changing one of the pairing partners of an existing base pair within the same loop. Note, that the latter was not used in this study, hence the neighbors have been greyed out.

#### 1.2 Base Pair Distance

The base pair distance $d_{\mathrm{BP}}(s,t)$ between any two structures $s$ and $t$ is the minimal number of moves, i.e. opening or closing of individual base pairs, that are required to transform structure $s$ into $t$. This distance measure is symmetric and can be expressed in set form as

$$d_{\mathrm{BP}}(s,t) = |s \cup t| - |s \cap t|. \tag{1}$$

Furthermore, using the indicator function

$$\delta_{ij}(s) = \begin{cases} 1, & \text{if } (i,j) \in s \\ 0, & \text{otherwise} \end{cases} \tag{2}$$

that denotes whether or not base pair $(i,j)$ is part of structure $s$, we can rewrite the base pair distance as

$$d_{\mathrm{BP}}(s,t) = \sum_{i<j} \left(\delta_{ij}(s) + \delta_{ij}(t) - 2\delta_{ij}(s)\delta_{ij}(t)\right). \tag{3}$$

##### 1.3 Sampling with the Temperature elevation method

To increase the structural diversity obtained from Boltzmann sampling, Kucharík et al. [9] developed an iterative scheme where in each round, after a specified number of samples is drawn, the sampling temperature $T$ is artificially increased if sufficient samples at a given energy level have been observed. For that purpose, they use an extra scaling factor $\xi \geq 1$ to effectively level the probability distribution of the ensemble. In particular, the probability of a structure $s$ then becomes

$$p_\xi(s) = \frac{e^{-\beta(\xi)E(s)}}{Z_\xi} \quad \text{with} \quad \beta(\xi) = \frac{1}{\xi kT} \tag{4}$$

and corresponding partition function $Z_\xi = \sum_{s\in\Omega} exp(-\beta(\xi)E(s))$. Consequently, the probability distribution becomes uniform for $\xi \to \infty$, so special care has to be taken when the normalization factor $\xi$ is elevated.

#### 2 RNAxplorer details

##### 2.1 Guiding Potentials and the RNA folding Recursions

The general idea to alter the Boltzmann probability of particular structures $s$ using guiding potentials with respect to a fixed structure $\hat{s}$ is to apply pseudo energies $E_c(\hat{s}, s)$ to their nearest neighbor energies $E(s)$. Hence their total energy becomes $\hat{E}(s) = E(s) + E_c(\hat{s}, s)$ and the corresponding equilibrium probabilities change to

$$\hat{p}(s) = \frac{e^{-\beta\hat{E}(s)}}{\hat{Z}}, \quad \text{with} \quad \hat{Z} = \sum_{s\in\Omega} e^{-\beta\hat{E}(s)}. \tag{5}$$

In practice, however, the partition function $\hat{Z}$ is never computed through exhaustive enumeration, but a dynamic programming (DP) scheme instead. Thus, no structure $s$ will ever appear as a single derivation of the underlying folding grammar. This implies that it is impossible to target individual structures, but only the substructures it is decomposed to [11], which consist of (parts of) loops and base pairs. Consequently, any term $E_c$ that acts on a particular substructure will always affect all structures that share this substructure.

To apply the general idea of guiding potentials in the context of the DP recursions that compute $\hat{Z}$, we need to decompose $E_c$ into its respective loop-dependent components. Each derivation of the folding grammar then receives a part of the contribution that corresponds to the substructures it covers. In particular, pseudo energy contributions have to be added whenever a (sub)structure is split into smaller components, or a base pair is added. We therefore extend the full nearest neighbor energy model for RNA secondary structures as follows

$$\begin{aligned}
\hat{Z}_{i,j} &= \hat{Z}_{i,j-1} \cdot e^{-\beta E_c^1(i,j)} \\
&\quad + \sum_{i \leq u < j} \hat{Z}_{i,u-1} \cdot \hat{Z}^B_{u,j} \cdot e^{-\beta E_c^2(i,j,u)} \\
\hat{Z}^B_{i,j} &= e^{-\beta\left(\mathcal{H}(i,j) + E_c^3(i,j)\right)} \\
&\quad + \sum_{i<p<q<j} \hat{Z}^B_{p,q} \cdot e^{-\beta\left(\mathcal{I}(i,j,p,q) + E_c^4(i,j,p,q)\right)} \\
&\quad + \sum_{i<u<j} \hat{Z}^M_{i+1,u} \cdot \hat{Z}^{M'}_{u+1,j-1} \cdot e^{-\beta\left(a+b+E_c^5(i,j,u)\right)} \\
\hat{Z}^M_{i,j} &= \hat{Z}^M_{i,j-1} \cdot e^{-\beta\left(c+E_c^6(i,j)\right)} \\
&\quad + \sum_{i \leq u < j} \hat{Z}^B_{u,j} \cdot e^{-\beta\left((u-i)\cdot c + b + E_c^7(i,j,u)\right)} \\
&\quad + \sum_{i<u<j} \hat{Z}^M_{i,u-1} \cdot \hat{Z}^B_{u,j} \cdot e^{-\beta\left(b+E_c^8(i,j,u)\right)} \\
\hat{Z}^{M'}_{i,j} &= \hat{Z}^B_{i,j} \cdot e^{-\beta b} \\
&\quad + \hat{Z}^{M'}_{i,j-1} \cdot e^{-\beta\left(c+E_c^9(i,j)\right)}
\end{aligned} \tag{6}$$

where the additional pseudo energy contributing parts $E_c^1, \ldots, E_c^9$ are highlighted in red. Here, $E_c^{3-5}$ handle cases where base pairs are added, while the other terms are only attributed to decompositions into smaller substructures.

The most simple case arises, when a guiding potential

$$E_c^{bp}(\hat{s}, s) = \alpha \cdot |s \cap \hat{s}| \tag{7}$$

only depends on the number of base pairs structure $s$ has in common with another, fixed structure $\hat{s}$. Then, the above loop-dependent pseudo energy contributions are

$$\begin{aligned}
E_c^3(i,j) = E_c^4(i,j,p,q) = E_c^5(i,j,u) &= \begin{cases} \alpha & \text{if} \quad (i,j) \in \hat{s}, \\ 0 & \text{otherwise} \end{cases} \\
E_c^1(i,j) = E_c^2(i,j,u) = E_c^6(i,j) = E_c^7(i,j,u) = E_c^8(i,j,u) = E_c^9(i,j) &= 0.
\end{aligned} \tag{8}$$

The contributions become more elaborate for guiding potentials where the strength of the potential depends on the base pair distance between $s$ and $\hat{s}$, e.g.

$$E_c^d(\hat{s}, s) = \alpha \cdot d_{\text{BP}}(s, \hat{s}). \tag{9}$$

In that case, the individual loop-dependent pseudo energy contributions depend on the inherent distance to $\hat{s}$ any particular decomposition imposes. This generalizes a strategy introduced by [5] and [12] respectively for a single and two reference structures. The overarching principle is to keep track of the base pairs, in the reference structures, that are conclusively ruled out by the choice of a DP derivation. Similar to the work of Lorenz et al. [12], the required distances can be computed in a pre-processing step using additional $\mathcal{O}(n^3)$ time and $\mathcal{O}(n^2)$ memory. The final contributions then become

$$\begin{aligned}
E_c^1(i,j) &= \alpha \cdot d_{\mathrm{BP}}(\hat{s}[i,j], \hat{s}[i,j-1]) \\
E_c^2(i,j,u) &= \alpha \cdot d_{\mathrm{BP}}(\hat{s}[i,j], \hat{s}[i,u-1] \cup \hat{s}[u,j]) \\
E_c^3(i,j) &= \alpha \cdot d_{\mathrm{BP}}(\hat{s}[i,j], \{(i,j)\}) \\
E_c^4(i,j,p,q) &= \alpha \cdot d_{\mathrm{BP}}(\hat{s}[i,j], \{(i,j)\} \cup \hat{s}[p,q]) \\
E_c^5(i,j,u) &= \alpha \cdot d_{\mathrm{BP}}(\hat{s}[i,j], \{(i,j)\} \cup \hat{s}[i+1,u] \cup \hat{s}[u+1,j-1]) \\
E_c^6(i,j) &= \alpha \cdot d_{\mathrm{BP}}(\hat{s}[i,j], \hat{s}[i,j-1]) \\
E_c^7(i,j,u) &= \alpha \cdot d_{\mathrm{BP}}(\hat{s}[i,j], \hat{s}[u,j]) \\
E_c^8(i,j,u) &= \alpha \cdot d_{\mathrm{BP}}(\hat{s}[i,j], \hat{s}[i,u-1] \cup \hat{s}[u,j]) \\
E_c^9(i,j) &= \alpha \cdot d_{\mathrm{BP}}(\hat{s}[i,j], \hat{s}[i,j-1]) \qquad (10)
\end{aligned}$$

where $\hat{s}[i,j] = \{(p,q) \in \hat{s} \mid i \leq p < q \leq j\}$ denotes the set of all base pairs $(p,q)$ formed by $\hat{s}$ on the subsequence $\sigma[i:j]$.

Both of the above examples are cases where a guiding potential becomes stronger the more base pairs structure $s$ shares with $\hat{s}$ (Eqn. 7), or the further apart both structures are in terms of base pair distance (Eqn. 9). However, in many cases it is useful to apply the opposite, i.e. the strongest contribution when $s$ does not share any base pair with $\hat{s}$, or when $s$ is identical to $\hat{s}$, respectively. To accomplish that, one can simply substract the respective distances from a maximum value, e.g. $|\hat{s}|$ or $d_{\max}(\hat{s}) = \max_{s \in \Omega} d_{\mathrm{BP}}(s, \hat{s})$. Additionally, it is convenient to limit the total contribution of $E_c$ to the value of $\alpha$ such that it becomes independent of the number of base pairs in $\hat{s}$ and the length of the sequence. Again, the above mentioned maximum distance value can be used as a scaling factor to accomplish that, and the guiding potentials discussed in this paragraph become

$$E_c^{bp}(\hat{s}, s) = \alpha \cdot \frac{|\hat{s}| - |s \cap \hat{s}|}{|\hat{s}|}, \qquad (11)$$

$$\text{and} \quad E_c^{d}(\hat{s}, s) = \alpha \cdot \frac{d_{\max}(\hat{s}) - d_{\mathrm{BP}}(s, \hat{s})}{d_{\max}(\hat{s})}, \qquad (12)$$

respectively. These guiding potentials are still decomposable to comply with the recursive DP scheme of RNA secondary structure prediction, since any distance measure in the respective loop-dependent contributions then only needs to be substracted from its corresponding maximal value.

Implementation-wise, we used the *soft constraints feature* of the `ViennaRNA Package` [11] to refrain from the tedious task to re-implement the partition function and Boltzmann sampling algorithms. This allowed us to focus on the loop-dependent contributions $E_c^{1-9}$ in our tool `RNAxplorer` which are then simply attached at run-time in a plug-in like manner to the efficient implementations of partition function computation and Boltzmann sampling in the `ViennaRNA Package`. Furthermore, our implementation thus easily handles entire sets of references structures $\hat{s}_1, \ldots, \hat{s}_k$ with individual contributions $\alpha_1, \ldots, \alpha_k$ as required for our adaptive sampling scheme presented in section 2.1 of the main paper.

#### 2.2 Negative vs. Positive Guiding Potentials

As pointed out in the main article, guiding potentials on particular (sub)structures can be favorable or unfavorable in terms of sampling probability whenever $(\alpha < 0)$ or $(\alpha > 0)$, respectively.

Let us first consider the favorable case where $\alpha < 0$. Given a fixed reference structure $\hat{s}$ both, the term

$$E_c(\hat{s}, s) = \alpha \cdot \frac{|s \cap \hat{s}|}{|\hat{s}|} \qquad (13)$$

and Equation (12), lead to an attraction of the sampling process to structures that are close to $\hat{s}$ in terms of the number of base pairs they share, or their effective base pair distance, respectively. On first sight, such an approach seems to make little sense, especially if $p(\hat{s})$ is already large compared to other structures. However, extending $E_c$ to multiple reference structures $\hat{s}_k$ not only increases the individual probabilities $p(\hat{s}_k)$, but at the same time, increases the probability for structures $s$ that are close to *all* the references $\hat{s}_k$. The base pair distance approach in (12) with multiple structures $\hat{s}_k$ can thus be imagined as a fuzzy triangulation in $\mathcal{L}$ that, to the largest extent, increases the probability for structures enclosed by the hyperspace defined by the vertices $\hat{s}_k$.

The second case, $\alpha > 0$, behaves complementary to the above. Instead of attracting the sampling process to the references, it is effectively diverted from them. Here, $E_c$ becomes the largest within the hypersphere defined by the vertices $\hat{s}_k$ and decreases on its outside, i.e. the more distant a structure is from the references the less penalty in terms of $E_c$ it receives. This easily enables one to penalize sets of base pairs, loops, or entire secondary structures.

##### 2.3 Making reference structures equiprobable

Theoretically, one could use a set of reference structures $\hat{s}_1, \ldots, \hat{s}_k$ to triangulate any particular structure $\hat{s} \in \Omega$ and make it the effectively most (least) stable, thus most (un)probable, one. However, this might require exponentially many references $\hat{s}_k$ to be part of $E_c$. Another, probably more useful application of guiding potentials in the context of stochastic sampling is to render any set of (reference) structures $\hat{s}_1, \hat{s}_2, \ldots, \hat{s}_k$ equally probable, i.e. $p(\hat{s}_1) = p(\hat{s}_2) = \ldots = p(\hat{s}_k)$, by constructing a guiding potential

$$E_c(\hat{s}_1, \ldots, \hat{s}_k, s) = \sum_k \alpha_k \cdot d_{\mathrm{BP}}(s, \hat{s}_k) \tag{14}$$

with pseudo energies $\alpha_k$ such that $\hat{E}(\hat{s}_1) = \hat{E}(\hat{s}_2) = \ldots = \hat{E}(\hat{s}_k)$. The respective $\alpha_k$ arise from observing that (i) $\hat{E}(s) = E(s) + E_c(\hat{s}_1, \ldots, \hat{s}_k, s)$ and (ii) both, $E(s)$ and $d_{\mathrm{BP}}(s_i, s_j)$, can be precomputed for any (pair of) reference structure(s) $\hat{s}_k$. This results in a system of linear equations with unknown variables $\alpha_1, \ldots, \alpha_k$ that can be easily solved. For instance, to determine the variables $\alpha_1$ and $\alpha_2$ for two structures $\hat{s}_1, \hat{s}_2$ leads to the following pair of equations

$$\begin{aligned} -E(\hat{s}_1) &= 0 \cdot \alpha_1 + d_{\mathrm{BP}}(\hat{s}_1, \hat{s}_2) \cdot \alpha_2 \\ -E(\hat{s}_2) &= d_{\mathrm{BP}}(\hat{s}_2, \hat{s}_1) \cdot \alpha_1 + 0 \cdot \alpha_2 \end{aligned} \tag{15}$$

and infinitely many solution pairs $\alpha_1, \alpha_2$ that reflect the many choices to actually set the total values $\hat{E}(\hat{s}_1) = \hat{E}(\hat{s}_2)$, thus the effective probabilities $\hat{p}(\hat{s}_1) = \hat{p}(\hat{s}_2)$. In practice, a reasonable choice may rely on setting these probabilities equal to that of an auxiliary proxy $\bar{s}$ which is not part of the $E_c$. This extends the linear system to

$$\begin{aligned} -E(\hat{s}_1) &= 0 \cdot \alpha_1 + d_{\mathrm{BP}}(\hat{s}_1, \hat{s}_2) \cdot \alpha_2 \\ -E(\hat{s}_2) &= d_{\mathrm{BP}}(\hat{s}_2, \hat{s}_1) \cdot \alpha_1 + 0 \cdot \alpha_2 \\ -E(\bar{s}) &= d_{\mathrm{BP}}(\bar{s}, \hat{s}_1) \cdot \alpha_1 + d_{\mathrm{BP}}(\bar{s}, \hat{s}_2) \cdot \alpha_2 \end{aligned} \tag{16}$$

with only one solution for the variables $\alpha_1$ and $\alpha_2$.

Note, that using the guiding potential approach the probabilities $\hat{p}(\hat{s}) = exp(-\beta E(\hat{s}))/\hat{Z}$ themselves can not be set to particular absolute values *a priori*. Only their relations are preserved. This is due to the fact that the approach acts on the level of free energies and a particular probability can only be known once the partition function $\hat{Z}$ that includes all $\hat{E}$ is (re-)computed. In most cases it therefore remains unclear which structures $\hat{s}_1, \ldots, \hat{s}_k$ and what pseudo potentials $\alpha_1, \ldots, \alpha_k$ to choose such that any $\hat{s}$ becomes highly probable. Moreover, depending on the shape of the underlying energy landscape, a large number $k$ might be required to lower the probability of low free energy structures in the vicinity of $\hat{s}$. Otherwise, upon Boltzmann sampling, such unwanted structures can easily be overrepresented in corresponding sample set.

In Figure S2 we show an example using the *SV-11* sequence of our benchmark set. Here, we set the $\alpha_k$ values for each reference structure $\hat{s}_k$ such that $\hat{E}(\hat{s}_k) = E_{\text{MFE}}$, i.e. all reference structures are lifted to the energy level $E_{\text{MFE}}$ of the MFE structure as present in the undistorted landscape. The first two fixed references $\hat{s}_1$ and $\hat{s}_2$ correspond to the MFE and meta-stable state structure, respectively. When sampling from the distorted landscape, the sample set is dominated by structures that are structurally close to $\hat{s}_1$ or $\hat{s}_2$, as seen in Fig. S2(a) and Fig. S2(b) for sample sizes of $10^4$ and $10^6$ structures, respectively. In the latter, one can already observe a number of structures in between the two references. Those are close to both of the references and thus receive fairly large amounts of the energy boni $\alpha_1$ and $\alpha_2$. Adding a third and fourth reference structure $\hat{s}_3$ and $\hat{s}_4$ into the system of equations then already leads to a much larger triangulation effect. The resulting sample sets are depleted of structures close to any of the (three or) four references $\hat{s}_{1-4}$, but enriched in structures that keep the right distance from all of them, as seen in Fig. S2(c) and Fig. S2(d).

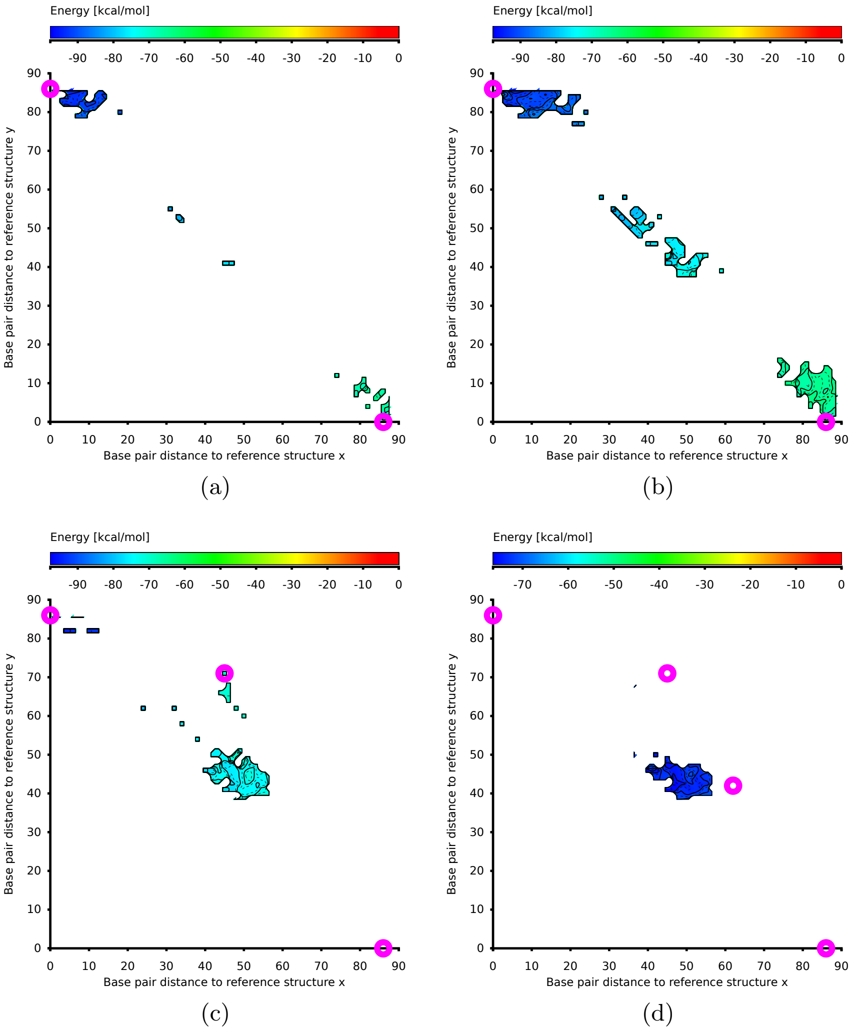

Figure S2: Sampling with up to 4 equiprobable reference structures using the *SV-11* benchmark sequence. The sample set is projected into 2D distance class space using structure $\hat{s}_1$ and $\hat{s}_2$ as references, where $\hat{s}_1$ is the MFE structure (magenta circle upper left), and $\hat{s}_2$ is the meta-stable state (magenta circle lower right). For each sample set we determine the lowest free energy representatives of the corresponding distance classes, whose free energy is color-coded in the plot. Sample sizes of (a) $10^4$ or (b) $10^6$ structures yields samples close to any of the two references. Upon increased sample size, as shown in (b), the sample set also consists of structures that are close to $\hat{s}_1$ and $\hat{s}_2$ at the same time. For sample sizes of $10^5$ structures and a third reference structure $\hat{s}_3$ (magenta circle mid top) as shown in (c), the sampling process is mostly attracted to structures close to all instead of any individual reference structures. With the same sample size but an additional fourth reference structure $\hat{s}_4$ (magenta circle mid right) this triangulation effect becomes even stronger, as shown in (d).

##### 2.4 Clustering Secondary Structures

To optionally reduce the number of output structures obtained from our `RNAxplorer` program, we implemented a clustering post-processing step that effectively reduces each cluster to its representative. For that purpose, we resort to a divisive clustering scheme using the `DIANA` heuristic [8]. Since this divisive algorithm eventually places each structure into a single-element cluster one requires a quality measure that indicates a suitable partition at which further divisions are stopped. In an earlier approach, Ding et al. [3] used the Calinski-Harabasz (CH) index for sample sets of RNA secondary structures that becomes maximal for an optimal set of clusters.

However, for this work, we resort to a much simpler heuristic that only analyzes the diameters $\varnothing(c) = \max_{s_1,s_2 \in c} d_{\mathrm{BP}}(s_1, s_2)$ of the clusters $c \in C$ at each step of the recursive DIANA scheme. We assume the total set of clusters $C$ to be sufficiently compact if either of the following two conditions holds: (i) no cluster diameter exceeds a pre-defined threshold $\tau_1$, i.e. $\nexists c \in C \mid \varnothing(c) > \tau_1$, or (ii) the average diameter $\bar{\varnothing} = \frac{1}{|C|} \sum_{c \in C} \varnothing(c) < \tau_2$ falls below a second pre-defined threshold $\tau_2$. Finally, for each cluster $c$ a representative structure $s_c \in c$ is chosen, e.g. the one with lowest free energy or simply the centroid, and provided as (reduced) output.

### 3 Data

For our benchmarks we compiled two sets of benchmark sequences:

1. Randomly generated sequences with lengths of 50 to 300 nucleotides in steps of 50 nucleotides. For each sequence length, ten sequences were generated with equal probability for all bases of the RNA alphabet $\Sigma = \{A, U, G, C\}$.

2. A set of 9 natural and artificial RNA sequences that are known to exhibit at least two distinct functional structure states. The actual sequences and corresponding references are available in Table S1.

| Name | Organism | Length | Sequence | Ref. |
|---|---|---|---|---|
| adenine riboswitch of *add* gene | V.vulnificus | 113 | GGCUUCAUAUAAUCCUAAUG AUAUGGUUUGGGAGUUUCUA<br>CCAAGAGCCGUAAACUCUUG AUUAUGAAGUCUGUCGCUUU<br>AUCCGAAAUUUUAUAAAGAG AAGACUCAUGAAU | [10] |
| lysine riboswitch of *lysC* gene | B.subtilis | 233 | GAAGAUAGAGGUGCGAACUU CAAGAGUAUGCCUUUGGAGA<br>AAGAUGGAUUCUGUGAAAAA GGCUGAAAGGGGAGCGUCGC<br>CGAAGCAAAUAAAACCCCAU CGGUAUUAUUUGCUGGCCGU<br>GCAUUGAAUAAAUGUAAGGC UGUCAAGAAAUCAUUUUCUU<br>GGAGGGCUAUCUCGUUGUUC AUAAUCAUUUAUGAUGAUUA<br>AUUGAUAAGCAAUGAGAGUA UUCCUCUCAUUGC | [10] |
| 5' leader of MS2 | coli phage | 130 | GGGUGGGACCCCUUUCGGGG UCCUGCUCAACUUCCUGUCG<br>AGCUAAUGCCAUUUUUAAUG UCUUUAGCGAGACGCUACCA<br>UGGCUAUCGCUGUAGGUAGC CGGAAUUCCAUUCCUAGGAG GUUUGACCUG | [6] |
| TPP riboswitch of *thiamine* gene | B.subtilis | 185 | GCAGAACAAUUCAAUAUGUA UUCGUUUAACCACUAGGGGU<br>GUCCUUCAUAAGGGCUGAGA UAAAAGUGUGACUUUUAGAC<br>CCUCAUAACUUGAACAGGUU CAGACCUGCGUAGGGAAGUG<br>GAGCGGUAUUUGUGUUAUUU UACUAUGCCAAUUCCAAACC<br>ACUUUUCCUUGCGGGAAAGU GGUUU | [10] |
| adenine riboswitch of *ydhL* gene | B.subtilis | 110 | AUUAUCACUUGUAUAACCUC AAUAAUAUGGUUUGAGGGUG<br>UCUACCAGGAACCGUAAAAU CCUGAUUACAAAAUUUGUUU<br>AUGACAUUUUUUGUAAUCAG GAUUUUUUUU | [10] |
| cyclic-di-GMP riboswitch of *tfoX* gene | C.desulforudis | 124 | ACCCCGAAAGGGCAAACCGG UACGAAAGUCCGGGACGCAA<br>AGCUACGGGUCCUUAAGUUC CAUGGGGAAUAGGACGGCUG<br>AGCCGCUGGGGUUAUUACUU UCGCGGAGCCGCCCUAUGGG GCGG | [10] |
| SAM riboswitch of *metE* gene | T.tenocongensis | 134 | CUCUUAUCAAGAGAGGUGGA GGGACUGGCCCGAUGAAACC<br>CGGCAACCAGCCUUAGGGCA UGGUGCCAAUUCCUGCAGCG<br>GUUUCGCGUUGAAAGAUGAG AGAUUCUUGUAGUCUCUUCU UUUAGCGAAGGGAC | [10] |
| SV-11 | artificial | 115 | GGGCACCCCCCUUCGGGGGG UCACCUCGCGUAGCUAGCUA<br>CGCGAGGGUUAAAGCGCCUU UCUCCCUCGCGUAGCUAACC<br>ACGCGAGGUGACCCCCCGAA AAGGGGGGUUUCCCA | [1] |
| ground state | | | (((.(((((((((((((((( ((((((((((.(.((((((<br>((((((((..((((...))) )..))))))))))))))).).<br>))))))))))))))))))). .)))))))))..))). | |
| metastable state |  |  | (((((((((((...)))))) )..((((((((((....)))<br>)))))))........)))). ....((((((((........<br>)))))))).((((((((... ..))))))))..... |  |
| guanine riboswitch of *xpt-pubX* gene | B.subtilis | 148 | CACUCAUAUAAUCGCGUGGA UAUGGCACGCAAGUUUCUAC<br>CGGGCACCGUAAAUGUCCGA CUAUGGGUGAGCAAUGGAAC<br>CGCACGUGUACGGUUUUUUG UGAUAUCAGCAUUGCUUGCU<br>CUUUAUUUGAGCGGGCAAUG CUUUUUUU | [10] |

Table S1: **Functional RNAs taken for our benchmark analysis.** Most sequences are bacterial riboswitches previously used in the benchmark of `RNASLOpt` by Li and Zhang [10]. The artificial SV-11 sequence represents an extreme case of a self-induced riboswitch where the two functional states are separated by more than 30 kcal/mol.

| Length | Number | Sequence |
|---|---|---|
| 50 | 1 | GUAAUGUCUAAUCGGGAAAG AAGCAGCAAAGUUAGACUGC CUUGCAGACA |
| 50 | 2 | CGAGGUCGUAGAGUAACCCG CACCUACACCUCAGGUGAUC AACGGUAUUA |
| 50 | 3 | CAUGAUAGUCCUAGGGGGAG CAAUCAUUAUUACAAGAAGU UCCCCGUCCC |
| 50 | 4 | CCAGCGAGCUUUUUGCUUCG GUGUCAUAAAGGGCUCAGGU CAAAUAUCUA |
| 50 | 5 | UUAACAGCGGCACGAAAUUG GCCAUUGUCUAUUGUGAUGG AGGACAUGAU |
| 50 | 6 | AGAGGCUUCUCAUGUCGACC GUCCGCAGGUGCUCAUUACG CUUAAGUCAU |
| 50 | 7 | UCUAAACAACCCGUCAACCA GAUUCAUAGCCCCCAAACCC AAGCAUCCUC |
| 50 | 8 | UUAUACAGAGGCACGACUCC CAGCUUCUAAUUGACCCUUG AGGGCACGCC |
| 50 | 9 | GGUGUAAAAUUUUUGGGUCU AACCUGGCUAAAACAGCUUC GCUCGGCGUU |
| 50 | 10 | GACGAUUCCAUCGUUUGAUG GUCAGAAAAAUGAAAAUAAA GUAAUAGCCC |
| 100 | 1 | UUGGCCCAUAGGCCCCGACC GGGUGUCCCCAGUAGUGGAA CAUCUUUACACCGACUUGUG UACUGUAAGUAACCUCGCGA CGCUGACCGGGGGCGCGGCU |
| 100 | 2 | GCCUCACCACUGAAAGUAUC UUAACCUUUUUAGAGUCGCU CGUACCGCCCGGGAGAGCCU AUUGAUGAUAACGAGCCGUA CUUGUUUGUCGUUUAUAUCG |
| 100 | 3 | AGGCUGCGGUUUCUAAGUAU GGAAAUGCCGCCCGAUCGAC AUUCCGACAGCUUGGGAGAA UCCAGCUACCUGCUUUCGGU UCAUUGUUCUCCCUUCUUAA |
| 100 | 4 | GUGCAAGAGCCGGUGAGAUG AUGAGAAUUGGGUUGCUCCG UAUCAGUUCAAGAGAAGUAA UCUGGCACUGAUUACGUUCU CCACUUUUAUACAAAGAACA |
| 100 | 5 | UCAGCCUAGGGUUUUUCGGU UUCUCGUUCACACAUGGCUA AUGAACAGUGGCGGUGCGUG CUGUAACCGGGUCGGUGUAG UGCGUCUGAUAUGCGAGACG |
| 100 | 6 | ACCGAGGACACGGGUUGGGC UAACAGAAAGAAGUUAACCG AUGCUGCGUUUCGGAUGGUA AUUCCGGUAGCACCUUUGUU AUGCAAAUGUUUACAAAUUA |
| 100 | 7 | UGCUCACGUCCUCAGCAUCA CCUUUGUAUGCUCGAGGAUA AACGAGCGUUACUAGUCAAC CGUUCGGAAUGCUCUGGUAU AUCUCUGACAUAUAGGGUCG |
| 100 | 8 | AACAGGGGGUAAUUCAUUAU ACGACUUCCCUCUAGAAUUA GAACAUCCACAUACCUUUAC UUAAGCCGACCUAUGCUAGA UCUCGGGUCUUUCCCUAUAC |
| 100 | 9 | GAAAGAAACCCCCCUAGUCU GCGCGCGCCUGUUCAGGAAA CUUAGUUUUGUAGUGUAAGG CUUAGAGGGCUCGGCCAACU AACAACUAUUUCGGGAGUUU |
| 100 | 10 | GCUUGAGUCCUCUGACAAGU AAUAAAGCCACCCUAAACGC UCGCCCCCCGACUACGUGCC ACACUUUUAUCCCUUAACUG GACCCAGUUUCAAUGCCAGU |
| 150 | 1 | GACUUCUCGGAAUGAGAAAG CGUCGGCCACACAGCUCAAU CUGGGAAGUUCCGGUAUGCC GUUGCAAGAUCCGGCGUUUG AGGCCCACUGGUGAUUUCCG<br>UUUCUCACUUCCACCGGCGU AUUUUGUAGAGACGCGUGAA UGGGCGAACA |
| 150 | 2 | GUAUCAACUAUAGUUAUCGG AGCGCAGAUUUAUUCCUCAA UGACUCGCAGCUACCUCCGG GGGUGGUAACUAUGAGAUUA GGUAGGUUUCCGAGCACCAU<br>CUCCUUUCAAAUAGCAAUCC UUGUCGAGCGUCUGAUAGCA CUAUGGUUAA |
| 150 | 3 | CAGUGCGGCGAGCUGAUAAC GCGCUAGACCGGAUCGUUAG CCGGAUCGGCCGGGUGGCUG AUUAUACUAAGCUAACUCCA GCAAUUGUCCGUAAAAUGUG<br>AUUUUCAGACAGACGUUUUG UAUGGCUAGACACCUUGGAU GGCGGGUGAU |
| 150 | 4 | ACACGGCUACUUAUAGAUUG GGGCCCAGGCGUCUGCUUAC GAUUCUACUGAGCUGAUGUA CGGUGGAGUUUAGUCUAACU CAAGCUAUGGCAUCGACGGU<br>AGCGCUAAGCAAGUACCUCG UUCGUUGACGAGUCGGUUAA AAGUUUCACC |
| 150 | 5 | UGUGUGCCUGGGCAUCUUUA GACACCCGGUGGCUACGGUU UCUCGUCCGCAAAGAUAGAA CUAGACCGUGUGAUUCCGAC AAAUCCAGCGGCUUGCGUUU<br>GGACGAGAAACUGAAUAGCC UCACGGCAAUCCAUAGUUCA UCUCCAGCCU |
| 150 | 6 | ACGUCCCUUGCUUCCUAGUA GUUUCGGAUACUCAUCCAUA UCCCGUACCUGAUAUCGAAA CCACAAUGGGUAAUGUGUCG CCAGGACACCCGGUUUGCUC<br>UUCUCGAAGACGUACAAUGA CCAAGCGUCUGAUGUUCUUU CAUCUCCCCA |
| 150 | 7 | GGGAGGUGUCCGCGAACCGA UUUAGUGCGCCUACGGCGAC UGGGUCCAGGAGUACUCUCG GUUAUGGAAACUACUGAAAA CGAGAUCAAGUCAUAGUGGU<br>CAGGGAGGACCUGGAUGUCU AGGCAGAGGGAAUGAAGUGU GAAAUAGGUC |
| 150 | 8 | UCUCAGUCAGCGGUGGUAGC UCCUGCACGAGAGGCCCACG AAGUGGCCUAUCUGCUCCUU GCGGUAUGAUGAGACAACCU CCUGGCCGUUGCCACUAUUA<br>GUCGUUUCUGUUGCUAUGUC GUCCACGCUAGACAAGGAAA AGAACCCACG |
| 150 | 9 | CGUACAGAGGUUAUACCUCA ACUAAUCAUCUCAUCUUUUC AAAUAUAAGCUCAUCAGUGA GACUCACGAGUUGAUUCCUC CUUCUUAUGCCAUCGACAAU<br>UUACGGCGGUCGAGCACGGG GCCGUAUUCUGAUUGGAACC AAGAUAAAGU |
| 150 | 10 | CCGCCAGUGUGGCCGGCGAG CUUCUAGAACCGCCCCCCGA AAAACCAAGAAUGACUUUAG UGUAUCAGGUGCAGCCCGGC AAAACGGUAGGUACAAAGGG<br>AUCCGUAGCUCGCCCACCUA GAUUCAUUUUUCCGGUCCCU UAAGUAGUCG |
| 200 | 1 | CCGUUAAAGUAUGAAUAAGG CACUUACUAUACGCCAAGAC CAGGUCUCAUGUAACCCCAA UGACCUGUCGGGUGGGCGCA UAUGGGGUUGCUCAUCCUAU<br>CCCAAGUUUUUAAUCAACAA ACGUCUCUUAGUCUAUUUUG GCUGGGGGCGGCGAAAACUC UAAGGCUAAGUUCGUCCUAA CGGGGUGACAAAUAGGAAUG |
| 200 | 2 | UACCCCCACACGAAGUCGCU UAUCUCUCGGUUUACCAAAU AAUGAACCCGUAUGAUGCGC ACUGACUUCCCCUUUGGUAC CCCACGGGUAGCAUCGCCGA<br>ACAAUGGUGAUCGGUGAAGU CGCUGGUAGGAGUUACCGGU UGAAGAUGUCUUGCGAAGCU AUGCUGCUUGCCGCCAUUAC GUCAAAGGAGCGACGUAUAG |
| 200 | 3 | AGGAGAACUGGGGGUGAUGA GUCCGAUGAGGCGUGAACGA GUCUCAAGAGCUCCGAGGCU UAAUGAAACAGUACGGAAUA CCCGCUUCAGUGUUGGCUUU<br>AAGCCCUUCCCGUCACUCUA GUUUGCAAAUAUGAGGCACA CAGACACUCCCUCCGAGUAC GAGCUCUGUCCUGCGCGAGU UGUCAAGUUACCACCGUCGG |
| 200 | 4 | UGAUGUCUUACAGGCAGGUC UGCUAUCGCUUUCCCGAUCU AUCUCUACUGUUGCUCAGCG AUUGUCACCCACAGACGCAA UGCACACUUCUCGGGCCGGU<br>CAAGACUGAAGGGCACAAAG AGUAUUUUAGGCAUAUCAAA CAGAUUUUAGUUAGUGGAAC UGUACUCCAAUGUCCUCUGA UACUGAGGACGUAACCGAAU |
| 200 | 5 | UCGUAGAUUGCAGUAAAACA CUACACCUUGAUUAAGAAGA AGGGGAAGCGUUAUUUACGG CUCUAUUAUCGCGAUGUUCA GAGUUUUUAUGCACCGGUCA<br>CCCCAUUCGAAGGGACCUGU AAGUCGGAAAGCGCUCGCAA CUUUCGGACUGCGUCGUUCU GUCUGCCCAGUGAGCUCGAA CCCGGCUGUCGGCGAAGCGG |
| 200 | 6 | ACGAUAACUUUUUUACGUUU AUCAGCGUGGAUCAGUAGCG UUAGGAGGCAGGUACUUUUC AUCGCCGGAACAAUGUAGAU CCAGUGAACGCGGUUGUGUA<br>UAAUUUAGGUUGAUGCUCGG GACCGAAAACAGGGCGUCGA AAUGUAGCACAGCACCGCGC AGAUUCUUCUGACAGGUGUC CUAGGUACUAAGGCGGGAGA |
| 200 | 7 | AAUUCCCCAAGAGGGAUAAU GCGUGUCAGAGUCUCUGUGC UUAAGUCUAAGUCCACCAUG UGGCGUAUUUGUUUGAACGU CGGGCUAGUUACGGGAAAUU<br>CGGCUAAUAUGUUGCAAUAA CCACGUACGCGACGAAUUUA GCCACUAAGAUUGCGGCCGC UGCGUGGCCCUGUGCAAGUG GUGAAGGCAAGUAAGGAAAG |
| 200 | 8 | UGUUCUAUGCGUAGUUGUGU GGGCUUAGCAGUUGCAUGGC UCGUGUACUUUAAUUGCAAU CAUCCACUUCGAGUUGUGCG GUAAAGGGGACGCCCUCGUA<br>GGGCGCUUGUAUGGGUAAUG CCUCGUACCCUCCAAGCUUA UUCCAGUAAAGUUGGGGUCG UGGGCACGAUUCACGGUCCU GAGGAUACUACGAAUUGAAG |
| 200 | 9 | UGGAUUCACAAGCAGCCCCC AAUACUAUUUCGUACCAGUG UGGAGUCCUAGUCUUGGUGA ACGGGAAGUGGCAUCUCGGG CCACCGCUAAGCCAAGCACC<br>AGAUCGAGCCAUUAAGCGCC AGUGCCUUCAAUGGUCUUGA CGCAGUGUUAAUGCGACUUG GUAACGGUGCAUGACUAGGC ACUGGGUCUAUUAUUCUGGC |
| 200 | 10 | CCCCUGACUUCGCCAGUUCC CCCGACCUGUUUUUUACAGU CGCACUGGAGUCCUAGAUCU UGCUUGCGAUUAGCCAUUUG GAAUGGGCAUCGUGGGUGCG<br>UCCGGCGACGAGUUUUCUAC UGAAUGGGCGAGUAGUCACU AGAGUGUGGGGAUUCACAGA AUAGAAAACCAAUAACUGGG UGUGUAUGCAGGGAGGCAAU |

Table S2: **Random RNAs taken for our benchmark analysis.** These sequences are generated with probability 1/4 for the characters 'ACGU'.

| Length | Number | Sequence |
|---|---|---|
| 250 | 1 | UUGGAUUCUUCACUUAUCCU UCUCAGCUAGCGACGGGAUC CGGGCAAUGCGUCAGUUUGA ACCGCAUCGUAUCAGGAAAA UUCCCGGUCACGCCCAGUCG<br>AUGUGGAAGUUUUUGAGGCU GCGGUACACCAGUUCUACCU CAAAUGAAUAACAUCUCCAC UGACCACAUGUCUGGCGUGC CCCACCUCACGACAAAACUG<br>CUUAGUCCCUCUUUCCCGGG GCGCGCCGGAUGAAGCUAGC GGGCGCAAGG |
| 250 | 2 | CGGUAUAACAAUUGUUCGUU UGCGGCCAUUGUCUAACUCG CCUUGCUAACGAACCUCUAA CCUUCGUCACGUUGCGCCCA CCGGGUAAGCUGUGGCGUGC<br>CUUCUAGUGCCCCCUCCGGG CGAGAGUGGCACACAAAUCU ACGUUACACGCAAGCCAUUA AACGAACACAUUACAAUGCU CAUAGGUGCCUAGAUGAAAA<br>CACCGAUCCAAUGGCCGUGC UCAACUCACCCUGGCUCAUA AGGUCCCCAA |
| 250 | 3 | GCACGACGAUGAUUUCACUG GUUCUUCCAUUCCACUGUUU ACCUAGCACUAAACCGCGUC AUUUCUGAACGGACGGACCC ACAUUACCUGAACGACAGCG<br>UACGCUUCUCCGCACAGCUU AGGCGGUUUAAUACGAUCGA CUUGCUGCGAGAUGGUGCUG CUCUCAAGAAGAAAGAUUUC CCGGGCCGUCCUGGCGAAAA<br>GAGUGAGAAGCGACUUUCCU ACUGUCGGCAUGAAGUUAGU CAUAUAGUAU |
| 250 | 4 | UUGGAGAGACCCAUCAGCUA UCAGAAAAUCCUCUUACAUU AAAGUGCCCUCCCUUAACUC GGGAGACGCGGGGGAAGGCU GCGUGCAGCCCUUGUGCAAG<br>CUUCUUAAGAAAAGUCAUGC GAACUCAAAAUCCACUUGCA UUUAUACACACCUCAUCGGU GUGGAGCGGGUGCCGCCGUG GCGCCGUUAUUAUAUGGCAA<br>AGUUCGCUUUUAAUUGUGGU CAGCGAAUUUCUCCAAUACC GCGGCACAUG |
| 250 | 5 | AAUCGGCUGUCGCACCGGAG UACUUUGGACCAGUACAAUC GCAAAGGGAUCUGAUAGCAU CGUGACUCAUUUCAUAAGCU GAGAUAAUGAACGUCUAAAU<br>UUAUUUAAUCACCAGUCUUC UGUCAUGAGUCUCUUCGAUC UGAAUCAAACCUAACGUCUG UGGAGAAGGUAAAUAUAGCC GCAUAAACAAAUGCUAGUUG<br>CAAGGUAGAAUUGAUACGGA CUCAGGCGAAGGGGCGAGUC GGAUUCACUU |
| 250 | 6 | UCUCGUUAUAAUACAGUUCG AAUUGUGCACCGGUGCUAAA GUUUCGUGACCUCUGAUAGC GCAAGUGGCGCGCCAGAAUC UCGCUUGGUCGAUUUCCAAC<br>GCUCGCUAUCUAGCUAAGCC ACUUGGCCCAGUCUAGUGCG GCUGAACGAAAGCAAGCGCU AUCUACCCCUGGCGGCAUCA CUCAUGUAUAGGCGCAGCUG<br>GAAACACUCUAAGGCACCGG AAAGUAAAUUAACCGCUCUG CGACAAAUGA |
| 250 | 7 | UGAUCCCCAUUUACUUUGAC ACAUGGCUAGAGAGGGGAGA UACUAGCACUAGUUGCGAUC UUGCAAGGGACACAACCAUG UCAAAUAUAGAUUUUAUAAG<br>UGUGGGCGGGUCGUGGCUCG ACUGAACACAAGACCCUCAA GACAGGUCAGAUGCAUCUUU UCCCAGGCCACGCUGAUCCC AGACUCAUGCGCCUGAUAUC<br>UGGUUGGCCUCGAGGCGAGU UUAAAGGUUUAGACGUUGCU UCAAACAUUA |
| 250 | 8 | GCCUUCUCGAAGAAUAAGAU UAUCCUCAAAUUUGGCUAUA UGCCAAGAAUCUUUCUCCGA ACAAACCAUUACGCCCCCCU UGUUUCUGGAAAGCAUCAUG<br>GAUGCAAAUGCAGAUACGAU GAUCAAGAGACCCACUGCUU UGGUUCUUAAACAAUGCCUU AUGCUAAUACUUCGGUUACC CGCAUUCUCAAACCCAUAAU<br>CUCUGCUCACCCUCCAUGCA UCCCUGGUCGCUGGACGCGU UAGGGCACGA |
| 250 | 9 | ACAUUCGGCCUGGCGACGCG CGUCCCUGGGUAAGAGGCCG CUUUAGACUCAAUAAUCCGC ACGAUUACAGCUAAGAGUUG CAUGAUCUUUUACAUUACCC<br>UACCAUGGGCUAAAACACCA GGGCGAUAUUUAGCCCCGAG CCUAAACGUGCAACACUGGG CGAUCCGGGGUACGGCCCUG GAGCGCACCCGGCACUGACG<br>GGCGUUACUAGCUCGCUAAA UGAUUAAGCCCCUCAUAUUG AUGAUGAGAU |
| 250 | 10 | UUCUGUAGCGAACGGUCGUA GAAAACGUGUAGCUGGCGGG UUAAGUUAGUAACCGCUGUU UCCUCCCUGUGACGAGCAAG CCCGGGCAGUUCCUUUGGCU<br>GUCUAAAGCGCGAGCAUACG CAAGGGAGGGUUCAUAUCGC CCGCUUAGGCCUCCUCCCCA AAACCGGACGCUUUACUAAA AACUGAUGCUGGGCUGUGAU<br>GUCCACACCGUUCUAUGAAG GAUGAUACCGCUGACCGAAU CGUGUCGACU |
| 300 | 1 | AUUGUGACGAGCGUGUACAA CGGUACUAAAAACAAAUAAC UGCACGGCUUCGUAAGAUUU GUCAAAGGCACUGUAAGCGC UAACUAUCCGUACAGUCUAG<br>AGGGGUCCGUUUCAGGUGAA UUGUCUUCACGGUGGGGUGU UGAAGGUUCACCGUUUCUGG CUAGUAUGCAGAUAAGGAAG AAAGCUAUAAACACGAUAUA<br>AGGAAAAGGUAGUUCCUCGU AAAGAACGUGCUAACUAUGG AAAAAGGAGGUGGCAUCGGG GAUAUAAGUCGCUCAGUGCA GUCAGGAGUCGCUGCACUGC |
| 300 | 2 | UCUGAUUUUGCUGACGCACU UUAGACGAUAGGGUUCGAAC ACGAACUAUAACGCAUGGAU CCACCGACGCUAGUUAGAAA CUAUCGUUACUAGACGAUUA<br>GCGCAUCAAAGGGUAAACAG GCCGGAGCACGGUAUAGGGG AAUGUGAGAGCUGCCUUAUG UAUUACCCCAGGCUCCAUCC UAGCUCCCGAUCAACCCGGC<br>AUUCCCAUACCUACUGUAAU GCAAUCCGCGGCCCACCGCC CCAGCGGACCGCCAGGCUCC ACAGAGGUUUUAAGUUAAGA CUGGCCGUAUCGGGGGCGAG |
| 300 | 3 | CACCAGAGAUACCUGCCAGA UCCUAUAUAGUGGGGGACU CAGCCCCGGGGCUACCAUCG AAAAUAAGACAAAACGUCCU UGCCUCCGAUGUUCAUCCGC<br>CCACGCGACUACCUCCUAAA AAUACUGCACUGCGCCUUGA UACUCAUCGAAGGGCUGGAA AGAAAACGAUCGUGUGUCGG CCUAAGGUCUCUCAGCCUAG<br>GGUACCCGGUCCUCACGCCC GUGGAGUCCAGAUAGGUUGA AUUACAAAUAAGGACCACUG UCUCUGCGCGGUGACCCGAA GCGCACAGGCGCUUAGAAGU |
| 300 | 4 | CACGAAGUUAUCUCCUUGCC AGAAGAUAGCGUACCUCGCA AGGCCAAAAGUGUCGGUGUG UUUUGACAUCUCUAUCCCAA GUGGCGUCCUAGGAAACUGA<br>ACGAUAGGCUCAGGGUGGUG CGAAAGUGGGCUUAAACAUU AGGAAGCAGGAAUGAGCCGA GUAUAAAACAUCGCGUAAUC AUCGAGUGGGUCGUACUCUC<br>UUACUUAAAAGAUCGUCCAU GCGCAGUCGAAUAACCAAUU CUCGUAGACAUUACCGAAGA GCCGCCCAUCUAGAGCUUAU GGGGCUUUCGUGGGGAAGGU |
| 300 | 5 | AACUCCAAUUAACCAACGAC GUUGUUGGAUCUCCAUAACU GUGGUCGUAAUUCAUGUUAU UGCAGGGUAGUGCACCGAUG AUCAAACCAUGAUAACGCAG<br>GUUCCCUCUAGGAGCCUGUA UAGAGUUCGUGGUUAUAAGG UAUUGAGGUAGAGGUAAUCC UGCCGGUUGUGUCCGAAAGU AGUUAAGUGGAACUAAGAUC<br>AACAGUGGGGCUAUACCUAC UGGACCCAAAGCAACCCUCA UUGUGUGGUUUGUAUUUACG AAUUUGGAGCCCCAUCCCAA AACCAGUCUUAUACCUUCGU |
| 300 | 6 | CGAACACUGUACAGCACACU UCGUAUUGCUUGACCCGGCG CGGUAGGUUAGUGGGCCCUC GCACGCCUUCUAUGCCGACC CUGGACGGAAAUGUAAGUCG<br>GUCUAUCUAUUGCUCUCACC CAUUCACAUUCCGACACAGG UGCAUCGUGACGAACGAGCG CGCGAGGCGCACUGUUGAUA UACCGUUAACACGGGCCGCG<br>CCCUGACUCAUGAGUUCUCC CAGUAUGGCUAUCUCUUGAU GAUGGCUUGUGGGCUCUGAU GCGCUGAGUCGGUGUAGGGU AAAUAAGGGAAGAUGGUAUU |
| 300 | 7 | UGAAUUACCCGUCUAGAUCA CUGAAUAUGAAAACUUCAAG CACCUGUUUAGGGCGAAUGA UCCGUGUGAGGCAGAAUAAU CAUCACUGCAAGUCUAGUCU<br>AGACUGUAAGUUAUGCUAGA ACUACCUCGGCGACAGUGAG UAUGUGGACACCGGUUCCCG GGAGAGCUGCGAUUAUGUGC UCUACAGCGUUCUAGGUAUU<br>UGGAAGCUUUCGCCCGAGAU CCGUGCGGAGUUGUUGAUGG GCUGCGUUCCGAUUCUGUCC GCUAAUAAAAGGGGUGCUAA CCACAUCUUUACCCCCGACU |
| 300 | 8 | CAGUGUAUAUCCUCCCCUUC UGCCGCACGCUAUAUCGCCA AAGCCUGGGUUCUCCAUCUC UCUCUGACUCUCAUAGUCCC CGUGUCUCAGCAACGAUCCA<br>UAACUGGUCGCUUUUCACGG CCCAGUGUAUUUACACCAAA GCUACGUGUCACGGAGCAUG CCAGAACGCAAUCCUCGAGU UCUAUCGAUCUGAAUAUGUU<br>GCAUGCAAAUCGGAGAGACU GUGCAACACAUCGUGCUGCC CUGGGCGACCAGUUGUGCGA ACUAUUUCCCUCCGCCAUUU GUUGAGGAGGGGGAAUGUCC |
| 300 | 9 | CCGUAUCCCAGUUUAGAGAU GCUUUUAAGACCCUUUGAGG UUCGUGUACCCGUGGCGCUC GAUAACGCUAGCCCACGAGU GAGGGCCGGACCCGAUGUUC<br>UGCGGAAGCUGAAGCUCCGU ACUAGCACGGUAUUUUAGCU CUGAAACGACAAAAGACUGG CCACGCCAUCGCUAGAGGGU CGUCAACGCAGUGAAUUUAC<br>CCACACUUCUCCAGGACGCC UGGAACCCGCGUAUCUUACG AGUCUGGAUUUUGCUAAGUU UGUAGGUGUUAGGCAUUUAG GGGCCUGAGUUGAGUCAAAA |
| 300 | 10 | UAUACUUUGGCAGAGGUAGU UGACCAGCUGUAUGCUCAUC CAUCCGAAACGCCGUUCCGC UUGGCCUCGAGGCAACAGAA CUCUGGUUCGGCAGCUAUCG<br>AGACGCGUCAACAGUGGGUU UCCCUACUAGCUUGGGCAUU CUUGCUCGGACUGUGGCCUA UACUGCUGUCACCUGCAGGA CUCUUCCAGCCUGGGCGGGC<br>GAUGGCGAGGGGCUCCAUGC GGACUACUUGCGAGACGUGG AAACGAGGUAUUAUCGAUCC GUCAAAAGUGGCGACCAGUA UUGGCUAAGAUGGCUCAACC |

Table S3: **Random RNAs taken for our benchmark analysis.** These sequences are generated with probability 1/4 for the characters 'ACGU'.

### 4 Extended Benchmark

#### 4.1 Memory Requirements

We compared the memory requirements for the structure sampling tools `RNAsubopt` (in 'regular' and uniform sampling mode), `RNAlocopt`, `RNAlocmin`, `RNANR`, and the exhaustive enumeration method implemented in `RNASLOpt` against our novel guided sampling strategy `RNAxplorer`. For that purpose, we use the set of artificially created random sequences listed in Table S2 and S3. The observed memory requirements for varying sequence lengths and varying sample sizes can be found in Figure S3.

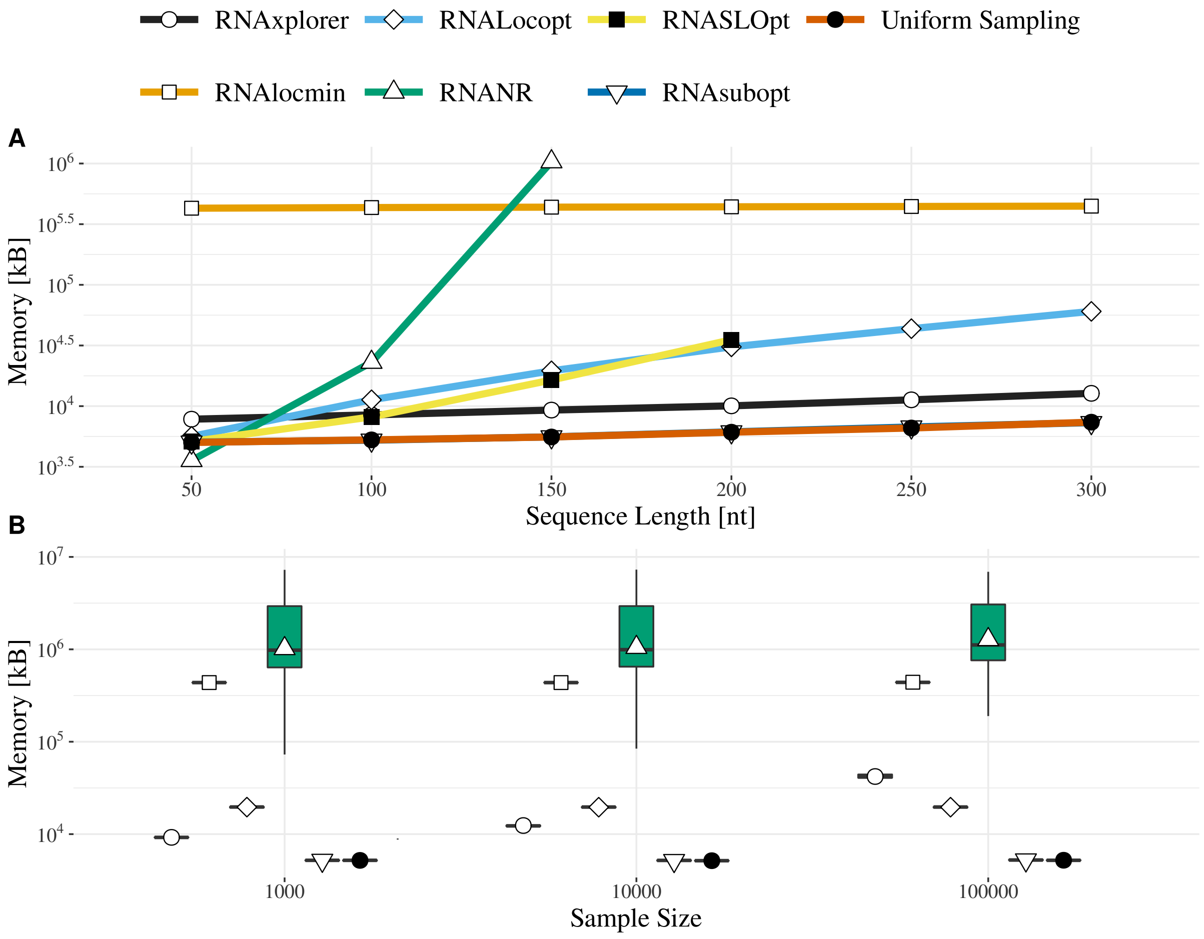

Figure S3: **Memory consumption**. Ten sequences of length 150 per tool and different sample sizes.

#### 4.2 Runtime and memory requirements on sample sizes

Next, we analyzed the runtime and memory requirements for varying sample sizes of $10^3$, $10^4$, $10^5$, and $10^6$ but constant sequence length of 150 nt. The runtime is depicted in figure S4. Here, `RNAlocmin` was the slowest with a steady runtime of about 150 seconds already for $10^3$ structures with only miniscule increase when the sample size was increased. Interestingly, the runtime and memory requirements of `RNANR` not only depend on the sequence length and number of samples, but also the sequence composition, clearly visible in the extent of the corresponding 1st and 3rd quantile. Its average runtime is about 17 and 18 seconds for $10^3$ and $10^4$ structures, respectively, and increases more than 5-fold for sample sizes of $10^5$ structures. Similar to `RNAlocmin`, the runtime of `RNAlocopt` did not significantly vary among different samples sizes with an average of 5 to 8 seconds. This renders it as fast as `RNANR` for sample sets of $10^3$ to $10^4$ structures. For sample sizes up to $10^4$ structures, our novel tool `RNAxplorer` is significantly faster compared to all methods mentioned before with average runtimes of 0.31 ($10^3$) and 1.96 ($10^4$) seconds. Still, even for $10^5$ structures, it is faster than `RNANR` and `RNAlocmin` and only slightly slower than `RNAlocopt` with 15.74. Finally, the fastest program measured is `RNAsubopt` (in both modes) with averages of 0.1, 0.54, and 4.94 seconds for sample sizes of $10^3$, $10^4$, and $10^5$ structures, respectively. Regarding the memory requirements, `RNAsubopt` is the least demanding with just about 5 MB for any sample size. Similarly, `RNAlocopt` and `RNAlocmin` display an almost constant memory requirement of about 20 MB and 440 MB, respectively. The memory consumption for our implementation of `RNAxplorer`, on the other hand, shows an increase from about 9 MB, to

12 MB and 42 MB for sample sizes of $10^3$, $10^4$, and $10^5$ structures, respectively. As mentioned before, even for the same sequence lengths, the resource requirements of `RNANR` also depend on the input sequence composition. This is clearly visible in the variance of the memory demand for each sample size. While still large compared to all the other methods, its memory requirements only slowly grow with the number of samples from an average of about 2.23 GB to 2.24 GB, and 2.4 GB for sample sizes of $10^3$, $10^4$, and $10^5$ structures, respectively.

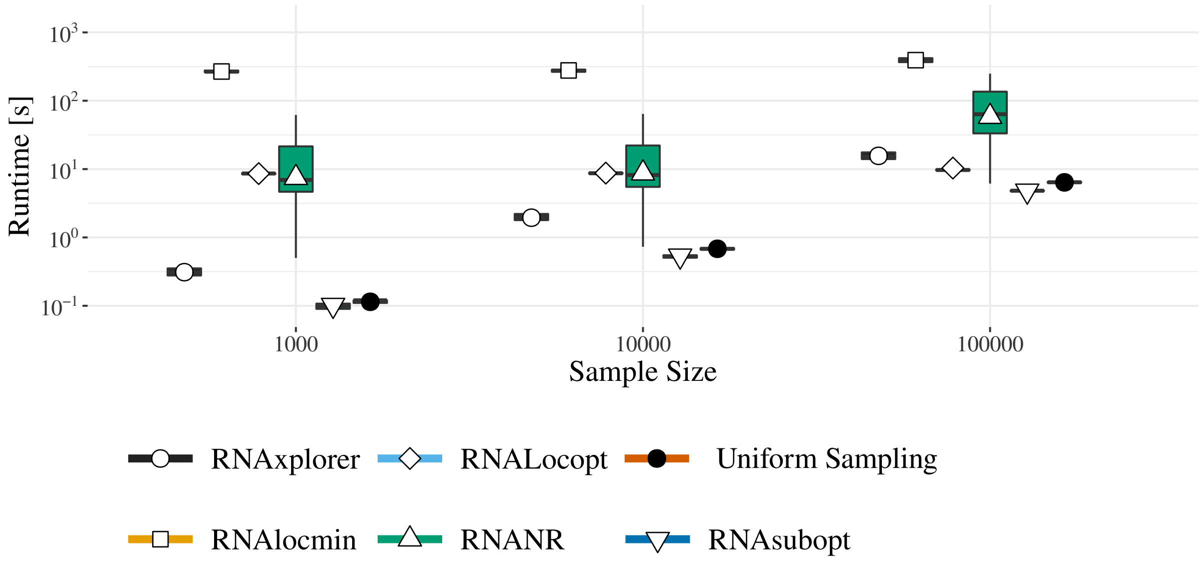

Figure S4: **Runtime comparison**. Ten sequences of length 150 per tool and different sample sizes.

### 5 Supplementary Validations

#### 5.1 Diversity Measures

In contrast to the majority of all methods we compare against, `RNASLOpt` implements an (exhaustive) enumeration approach, thus never yields redundant samples. Similarly, `RNANR` is an explicit non-redundant method but may yield different structures in successive sample rounds. We, therefore, excluded `RNASLOpt` from the below analysis. The remaining tools utilize Boltzmann sampling which is typically prone to over-sampling of low free energy states. Consequently, a sample set of $N$ structures may be highly redundant, i.e. it consists of only a few unique structures. To obtain a common coarse graining for all methods, we map each sample set to the corresponding set of local minima that is accessible through gradient walks.

For the below analysis, we requested sample sets of size $10^6$ per benchmark sequence from each method, mapped the structures in the sample sets to their corresponding local minima, and repeated the experiment 10 times, to obtain average measures.

**Number of Unique Local Minima** The most simple measure of redundancy is obtained from simply counting the number of unique local minima obtained from each method. The individual results of this analysis are shown in Figure S5A. Unsurprisingly, `RNANR` and near uniform sampling yield the least redundancy with values in the range of 96–100% and 51–99%, respectively. For all other tools we observe variations depending on the sequence composition, where the fraction of unique local minima for `RNAxplorer` and `RNAlocmin` are highest with 1–35% and 3–16%, respectively. `RNAlocopt` appears to be much more redundant with just 0–13% unique local minima reachable from the sampled structures. Finally, regular Boltzmann sampling produces the least number of unique structures with values between 0–13%. In decreasing order, the average fraction of unique structures over all 9 RNAs are 98.6% (`RNANR`), 83.6% (near uniform sampling), 14.4% (`RNAxplorer`), 8.5% (`RNAlocmin`), 2.7% (`RNAlocopt`), and 2.2% (`RNAsubopt -p`). To provide insight into the stochasticity effects of the different methods, we also show results for the individual sequences in Table S4. As depicted by the minimum and maximum percentage of the number of unique local minima retrieved from the different methods (last two columns of Table S4), the variation between different input sequences is rather high for all methods except `RNANR`. However, the maximum standard deviation for the 10 repetitions of the sampling procedure among all sequences (column 3 of Table S4) shows, that for a particular sequence the stochastic effect remains low. This demonstrates that the results obtained from the different methods remain stable over repetitions with the same input sequence. Here, `RNAlocmin` shows the largest effect of stochasticity, while `RNANR` turns out to be the most robust.

| Tool | Unique local minima [%] | max stdev | Unique local minima [%] | |
|---|---|---|---|---|
| | | | min | max |
| `RNANR` | 98.6 | 9.40e-03 | 96.3507 | 99.8294 |
| `RNAxplorer` | 14.4 | 4.94e-01 | 0.6061 | 35.1254 |
| `RNAlocmin` | 8.5 | 3.27e+00 | 2.7675 | 15.8947 |
| `RNAlocopt` | 2.7 | 2.43e-02 | 0.0133 | 13.3519 |
| `RNAsubopt` | 2.2 | 2.84e-02 | 0.0120 | 8.6026 |
| `Uniform Sampling` | 83.6 | 3.72e-02 | 51.3254 | 99.1782 |

Table S4: The average fraction of unique structures for all tools, 10 repetitions per tool for 9 sequences. The measure is computed for the unique part of the set of local minima reachable from the structures in each sample set. Standard deviations have been computed over 10 values per sequence. For each tool the maximum standard deviation is shown. The minimal and maximal value of unique structures is shown in the last column. An exception is `RNANR`, which could only be computed for 6 sequences.

**Mean Base Pair Distance** We then analyzed the similarity of structures within the unique parts of the obtained $10 \times 9$ sample sets. To this end, we computed a variation of the commonly used *mean base pair distance* $\langle d \rangle = \sum_{s,t \in \Omega} p(s)p(t)d_{\mathrm{BP}}(s,t)$ for Boltzmann weighted structure ensembles $\Omega$. In particular, we compute $\langle d \rangle$ over the unique part $\mathcal{S}^{\mathrm{u}} \subset \mathcal{S}$ of the sample set $\mathcal{S}$ and we treat the structures $s$ and $t$ equiprobable, i.e.:

$$\langle d' \rangle = \sum_{s,t \in \mathcal{S}^{\mathrm{u}}} f(s)f(t)d_{\mathrm{BP}}(s,t) \tag{17}$$

with $f(s) = \frac{1}{|\mathcal{S}^{\mathrm{u}}|}, \forall s \in \mathcal{S}^{\mathrm{u}}$. The smaller this value, the more similar the structures in $\mathcal{S}^{\mathrm{u}}$ are.

Note, that using Eqn. 3 the above equation can be rewritten without explicit pairwise computation for $s$ and $t$. Instead, one can derived it from the frequencies $f_{ij} = \sum_{s \in \mathcal{S}} f(s)\delta_{ij}(s)$ of the individual base pairs $(i,j)$ as

$$\begin{aligned}
\langle d' \rangle &= \sum_{s,t \in \mathcal{S}^{\mathrm{u}}} f(s)f(t) \sum_{i<j} \left(\delta_{ij}(s) + \delta_{ij}(t) - 2\delta_{ij}(s)\delta_{ij}(t)\right) \\
&= \sum_{i<j} \left( \sum_s f(s)\delta_{ij}(s) \sum_t f(t) + \sum_t f(t)\delta_{ij}(t) \sum_s f(s) - 2 \sum_s f(s)\delta_{ij}(s) \sum_t f(t)\delta_{ij}(t) \right) \\
&= \sum_{i<j} \left(2f_{ij} - 2f_{ij}^2\right) \\
&= 2 \sum_{i<j} f_{ij}(1 - f_{ij}) = \sum_{ij} f_{ij}(1 - f_{ij}).
\end{aligned} \tag{18}$$

Also note, that one can use the same transformation to compute the above mentioned $\langle d \rangle$ for Boltzmann weighted ensembles. The only difference to Eqn. (18) is that structure and base pair frequencies have to be replaced by their corresponding equilibrium ensemble probabilities, which is similar to the distance between two ensembles as shown in Eqn. (14) of [7].

Similar to the fraction of unique structures above, we computed the mean pairwise distance measure for the unique part of the set of local minima reachable from the structures in each sample set. For the sake of comparability, we normalized the distance measure by dividing through the individual sequence lengths, thus the units of this measure are base pairs per nucleotide (bp/nt). The results of this analysis are presented in Figure S5B.

Unfortunately, for this analysis, we were unable to apply `RNANR` to three of our benchmark sequences due to its large memory requirements. Thus, we omit the corresponding data for the lysine riboswitch *lysC* (233 nt), the SAM riboswitch *metE* (134 nt) and the tpp riboswitch *thiamine* (185 nt). The largest average diversity is found in the sample sets generated by `RNANR` (0.45 bp/nt), followed by `RNAxplorer` (0.44 bp/nt), `RNAlocmin` (0.31 bp/nt), `RNAlocopt` (0.26 bp/nt), `RNAsubopt -p` (average: 0.24), and *near uniform sampling* (0.23 bp/nt). Again, we analyzed the individual results that were used for averaging. As shown in the last two columns of Table S5, we find that the lowest variation among the 9 different sequences in our benchmark set was obtained from `RNAxplorer` and `RNANR` with $\min\langle d' \rangle = 0.4\ bp/nt, \max\langle d' \rangle = 0.48\ bp/nt$, and $\min\langle d' \rangle = 0.39\ bp/nt, \max\langle d' \rangle = 0.47\ bp/nt$, respectively. The remaining methods show a larger sequence dependence, which was highest for `RNAlocopt`. On the other hand, the maximum standard deviation (third column of Table S5) for the results obtained from 10 repetitions for the same sequence, again, shows little effect of the stochasticity of the methods. The most robust method in our benchmark turned out to be `RNANR`, followed by *near uniform sampling* with standard deviations in the order of $10^{-5}\ bp/nt$ and $10^{-4}\ bp/nt$. However, `RNAxplorer` and `RNAlocopt` also show only little effect of stochasticity with a maximum standard deviation in the order of $10^{-3}\ bp/nt$ each. The highest effect was found with `RNAlocmin` where the maximum standard deviation among the repetitions for individual benchmark sequences was in the order of $10^{-2}\ bp/nt$. Still, the results demonstrate that the stochastic effect of the different methods tested does not overly contribute to their general results.

| Tool | avg $\langle d' \rangle$ | max stdev | min $\langle d' \rangle$ | max $\langle d' \rangle$ |
|---|---|---|---|---|
| `RNANR` | 0.4531 | 3.69e-05 | 0.3911 | 0.4774 |
| `RNAxplorer` | 0.4426 | 3.82e-03 | 0.4024 | 0.4807 |
| `RNAlocmin` | 0.3075 | 5.74e-02 | 0.2036 | 0.3947 |
| `RNAlocopt` | 0.2554 | 2.90e-03 | 0.1325 | 0.3632 |
| `RNAsubopt` | 0.2441 | 3.72e-03 | 0.1229 | 0.3686 |
| `Uniform Sampling` | 0.2301 | 1.31e-04 | 0.1828 | 0.3194 |

Table S5: The average mean base pair distance (normalized by the sequence length) for all tools, 10 repetitions per tool for 9 sequences. The measure is computed for the unique part of the set of local minima reachable from the structures in each sample set. Standard deviations have been computed over 10 values per sequence. For each tool the maximum standard deviation is shown. The minimal and maximal value of mean base pair distances is shown in the last columns.

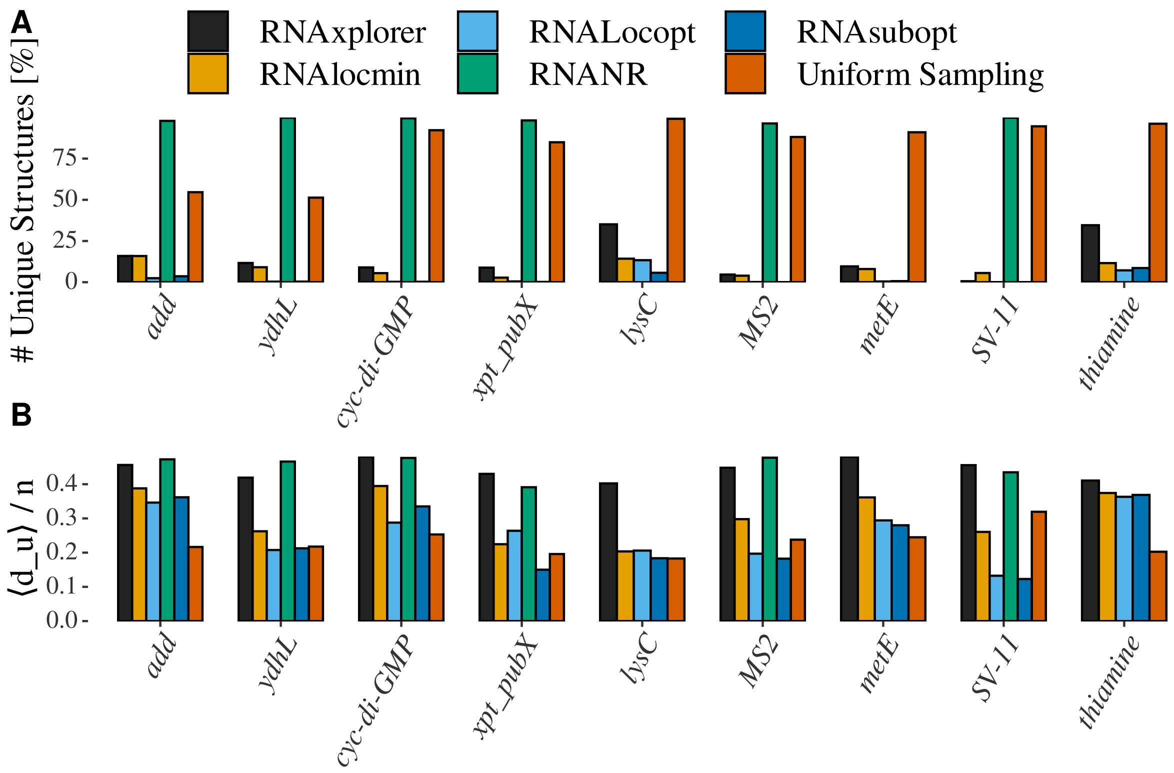

Figure S5: Sample Set Diversity. (A) Sample set redundancy. Shown is the average fraction of unique structures in 10 independently drawn sample sets of $10^6$ structures for the 9 functional RNAs from our benchmark set. (B) Normalized mean base pair distance $\langle d' \rangle / n$. This bar plot depicts the average mean base pair distance normalized by dividing through the individual sequences lengths. The average was taken from 10 independent sample sets of $10^6$ structures each. Results for `RNANR` for *lysC* and *thiamine* are missing due to too large memory requirements. The methods implemented in `RNANR` and `RNAxplorer`, as well as near uniform sampling clearly produce the largest diversity.

#### 5.2 Density of States

To quantify the coverage of the retrieved samples in terms of energy levels of the structures, we computed the density of states (DOS) [2] for each sample set. Here, structures $s \in \Omega$ are classified according to their free energy $E(s)$ to obtain densities, i.e. the number of structures, at particular energy levels. Most sampling methods are prone to over-sample the low free energy regime, but structures at higher energy levels might be important for modeling the dynamic behavior of RNA folding. While this measure does not tell us about the actual diversity within individual levels, it still allows one to investigate if structures at certain energy levels are over- or underrepresented. For that analysis, we chose the 9 benchmark sequences from Table S1 and computed the ground truth, i.e. the actual number of structures at each energy level using the tool `RNAdos` of the `ViennaRNA Package`. We then applied all Boltzmann sampling tools from our comparison, except for uniform sampling, to create sample sets of $10^6$ structures for each benchmark sequence. The deterministic enumeration of local optimal structures from `RNASLOpt` was also left out of this analysis, since we were unable to generate large enough sample sets even for very large $\delta$ values. After removing all duplicates from the samples, the respective remainder of structures was partitioned into their corresponding energy levels. The resulting structure counts were then plotted together with the ground truth to allow for visual inspection, as shown in Figure S6. Note that here, we depict the ground truth as a gray background, while the counts obtained from the sample sets are shown in color.

In this analysis we observed, that the three tools `RNAxplorer`, `RNAlocmin`, and `RNANR` perform best on covering a larger extent of the energy levels for most of the sequences in our benchmark. Due to excessive memory requirements, however, we were not able to obtain large enough sample sets from `RNANR` for 3 of the benchmark sequences, namely *lysC*, *metE* and *thiamine*. In general, the smallest energy range was covered by `RNAsubopt -p` and `RNALocopt`. Both of the former methods perform 'regular' Boltzmann sampling, though from different levels of coarse graining of the energy landscape. Here, our results clearly demonstrate the effect of oversampling of low energy levels. The area under the curve, i.e. the sum of all counts shown in color, reflects the total number of unique structures left over from the initial set of $10^6$ structures after duplicate removal. Thus, the larger the area and the further it is spread of the different energy levels, the more diverse the initial samples have been.

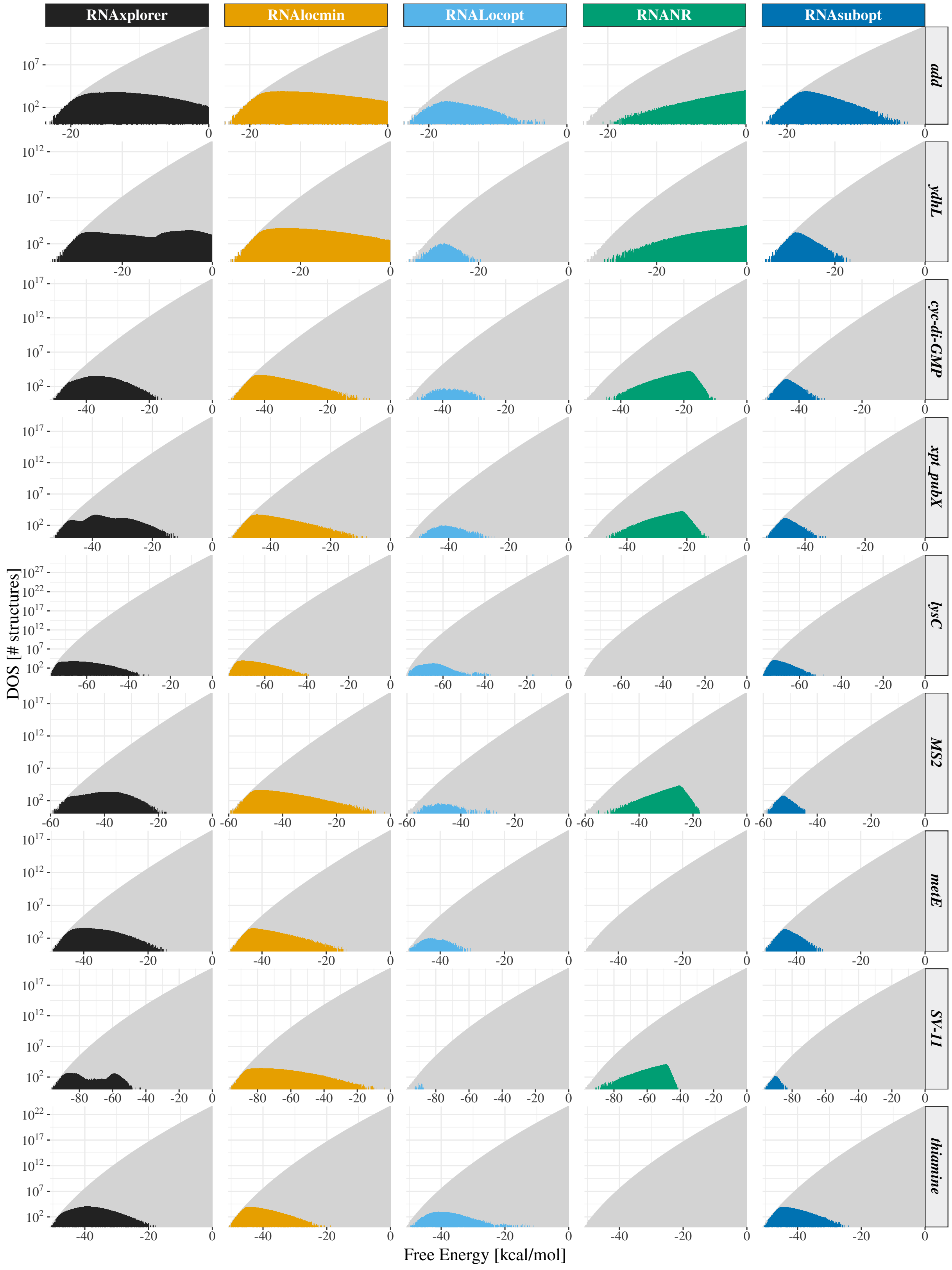

Figure S6: **Density of states for different sampling methods.** Each line shows the density of states (DOS) obtained from the unique part of a sample of $10^6$ secondary structures for one of the 9 benchmark sequences. The ground truth is depicted as gray background. Note, that the larger the area under the colored curves, the more unique samples were produced by the sampling method. Counts for energy levels above 0 kcal/mol are left out for better comparability against methods that oversample low energy levels. Due to its large memory requirement, we were unable to generate large enough sample sets for *lysC*, *metE* and *thiamine* using `RNANR`.

#### 5.3 Barriers Comparison

As proxy for a kinetics comparison, we made a comparison of the largest barriers. This is a good proxy since the larger barriers determine the long term folding behaviour of RNA folding kinetics. A proxy is necessary, because the kinetics can look very different for large RNAs if Arrhenius rates are used, since these rates neither take the neighbourhood of RNA clusters into account nor the cluster size.

A barriers tree was computed (with the tools `RNAsubopt` and `barriers` [4]) as reference landscape. Since the number of structures grows exponential with the sequence length, we used a 15 kcal/mol threshold above the MFE for structure enumeration with `RNAsubopt`. This still required about 200 GB for 100 nt long sequences on our test machine. Furthermore, the coarse graining with `barriers` was increased using a minimum energy barrier of neighboring basin of 3.0 kcal/mol using the `--minh` option. This effectively merges basins separated by lower energy barriers and enabled us to perform all pairwise energy barrier lookups in reasonable time.

For each sampling method, the samples were mapped to the original barriers tree of the connected component with the MFE structure (with –map-struc option of `Barriers` and -c for the connected component). In the next step, we extracted the 100 largest barriers for transitions into lower free energy basins.

The tables S6, S7, S8, S9, S10 list the coverage of the 100 largest barriers for different sample sizes ($10^3$, $10^4$ and $10^5$) and the average time for each sampling method. The coverage of the 100 largest barriers is given in percentage for the barrier trees of 10 random sequences and 10 runs for each tool.

The score for largest barriers is already high for small sample sizes ($10^3$) for all tools except for uniform sampling and `RNANR`. However, the tool `RNANR` finds the most number of local minima that match into the barriers tree for sequences with 70 nt and longer. In terms of largest barriers, the tools (`RNAxplorer`, `RNAlocmin`, `RNAlocopt`, `RNAsubopt`) are almost equally good for 100 nt sequences and $10^5$ samples. However, the number of covered local minima in the barriers tree is the highest with `RNAxplorer` and 14138 minima in contrast to the next best tool `RNAlocmin` with 9834 local minima. With an average time for sampling of 3.72 seconds, `RNAxplorer` produces the relevant samples for barriers kinetics much faster than `RNAlocmin` with 73.61 seconds.

| Tool<br>length: 50 nt<br>max. basins: 562 | sample size | | | | | | | | |
|---|---|---|---|---|---|---|---|---|---|
| | $10^3$ | | | $10^4$ | | | $10^5$ | | |
| | coverage [%] | | | coverage [%] | | | coverage [%] | | |
| | barriers | basins | $\bar{t}$ [$s$] | barriers | basins | $\bar{t}$ [$s$] | barriers | basins | $\bar{t}$ [$s$] |
| RNAxplorer (bp gp) | 88.90 | 34.70 | 0.02 | 98.50 | 71.53 | 0.18 | 99.80 | **95.73** | 1.90 |
| RNAxplorer (bpd gp) | 85.70 | 31.14 | 0.07 | 93.30 | 49.82 | 0.60 | 96.80 | 63.52 | 2.60 |
| RNAlocmin | **95.50** | **41.99** | 32.85 | **99.30** | **83.10** | 34.01 | **100.00** | 94.84 | 38.82 |
| RNAlocopt | 84.70 | 28.29 | 0.12 | 91.00 | 44.31 | 0.16 | 95.00 | 58.54 | 0.33 |
| RNANR | 54.20 | 17.44 | 0.03 | 54.20 | 17.44 | 0.03 | 54.20 | 17.44 | 0.03 |
| RNAsubopt | 85.50 | 30.43 | 0.01 | 91.40 | 45.20 | 0.07 | 96.10 | 62.10 | 0.37 |
| Uniform Sampling | 9.90 | 0.53 | 0.00 | 15.60 | 0.89 | 0.02 | 56.10 | 4.63 | 0.53 |

Table S6: Coverage of the 100 highest saddle points associated to the highest barriers between the deepest left and right minima in the barriers tree for ten 50 nt sequences. All values in the first column represent the best barriers coverage in percent. The middle column represents the sum of covered basins in the barrier trees for all ten sequences in percent. The third column is the average run time for each sampling tool. Two guiding potentials have been used for `RNAxplorer`. The base pair associated guiding potential, described in the main text (section **??** is (bp gp). The base pair distance based guiding potential (described in Eqn. (12)) is (bpd gp).

| | sample size | | | | | | | | |
|---|---|---|---|---|---|---|---|---|---|
| Tool | $10^3$ | | | $10^4$ | | | $10^5$ | | |
| length: 60 nt | coverage [%] | | | coverage [%] | | | coverage [%] | | |
| max. basins: 2407 | barriers | basins | $\bar{t}$ $[s]$ | barriers | basins | $\bar{t}$ $[s]$ | barriers | basins | $\bar{t}$ $[s]$ |
| RNAxplorer (bp gp) | 87.30 | **16.04** | 0.02 | 99.30 | 40.09 | 0.21 | 99.60 | 71.46 | 2.13 |
| RNAxplorer (bpd gp) | 82.60 | 13.09 | 0.14 | 92.60 | 24.47 | 1.16 | 97.50 | 38.10 | 3.37 |
| RNAlocmin | **89.70** | 15.16 | 33.67 | **99.50** | **42.29** | 36.35 | **100.00** | **71.83** | 57.60 |
| RNAlocopt | 80.20 | 11.09 | 0.22 | 90.10 | 19.57 | 0.28 | 97.30 | 31.95 | 0.52 |
| RNANR | 73.90 | 9.14 | 0.20 | 73.90 | 9.14 | 0.26 | 73.90 | 9.14 | 0.26 |
| RNAsubopt | 82.20 | 12.75 | 0.01 | 92.30 | 22.48 | 0.07 | 96.50 | 36.93 | 0.46 |
| Uniform Sampling | 0.00 | 0.00 | 0.00 | 0.00 | 0.00 | 0.00 | 0.00 | 0.00 | 0.00 |

Table S7: Coverage of the 100 highest saddle points associated to the highest barriers between the deepest left and right minima in the barriers tree for ten 60 nt sequences. The sample size is the number of structures that a tool produces in one run per sequence. In addition to the barriers, the basin coverage is shown and the average run time for each tool. Two guiding potentials have been used for `RNAxplorer`. The base pair associated guiding potential, described in the main text (section **??** is (bp gp). The base pair distance based guiding potential (described in Eqn. (12)) is (bpd gp).

| | sample size | | | | | | | | |
|---|---|---|---|---|---|---|---|---|---|
| Tool | $10^3$ | | | $10^4$ | | | $10^5$ | | |
| length: 70 nt | coverage [%] | | | coverage [%] | | | coverage [%] | | |
| max. basins: 5319 | barriers | basins | $\bar{t}$ $[s]$ | barriers | basins | $\bar{t}$ $[s]$ | barriers | basins | $\bar{t}$ $[s]$ |
| RNAxplorer (bp gp) | 84.10 | 7.82 | 0.03 | **94.90** | **27.69** | 0.24 | 97.40 | **63.98** | 2.44 |
| RNAxplorer (bpd gp) | 80.50 | 6.58 | 0.18 | 90.40 | 13.99 | 1.67 | 93.30 | 23.78 | 4.62 |
| RNAlocmin | **84.90** | 6.66 | 34.93 | 94.60 | 22.99 | 37.62 | **98.20** | 57.10 | 65.39 |
| RNAlocopt | 77.90 | 5.88 | 0.38 | 87.50 | 11.49 | 0.43 | 94.00 | 20.38 | 0.72 |
| RNANR | 78.40 | **10.40** | 0.49 | 78.40 | 11.00 | 4.54 | 78.40 | 11.00 | 6.66 |
| RNAsubopt | 80.60 | 6.32 | 0.03 | 88.00 | 12.15 | 0.07 | 93.30 | 22.88 | 0.50 |
| Uniform Sampling | 0.00 | 0.00 | 0.00 | 0.00 | 0.00 | 0.00 | 0.00 | 0.00 | 0.00 |

Table S8: Coverage of the 100 highest saddle points associated to the highest barriers between the deepest left and right minima in the barriers tree for ten 70 nt sequences. The sample size is the number of structures that a tool produces in one run per sequence. In addition to the barriers, the basin coverage is shown and the average run time for each tool. Two guiding potentials have been used for `RNAxplorer`. The base pair associated guiding potential, described in the main text (section **??** is (bp gp). The base pair distance based guiding potential (described in Eqn. (12)) is (bpd gp).

| | sample size | | | | | | | | |
|---|---|---|---|---|---|---|---|---|---|
| Tool | $10^3$ | | | $10^4$ | | | $10^5$ | | |
| length: 80 nt | coverage [%] | | | coverage [%] | | | coverage [%] | | |
| max. basins: 12872 | barriers | basins | $\bar{t}$ $[s]$ | barriers | basins | $\bar{t}$ $[s]$ | barriers | basins | $\bar{t}$ $[s]$ |
| RNAxplorer (bp gp) | **90.70** | 4.27 | 0.04 | 94.40 | **18.75** | 0.29 | **96.60** | **50.05** | 2.60 |
| RNAxplorer (bpd gp) | 86.70 | 3.48 | 0.35 | 92.20 | 8.86 | 3.36 | 93.80 | 17.79 | 8.16 |
| RNAlocmin | 88.40 | 3.48 | 36.43 | **94.50** | 14.76 | 39.36 | 94.60 | 39.29 | 67.81 |
| RNAlocopt | 81.90 | 2.92 | 0.59 | 90.80 | 6.32 | 0.65 | 92.90 | 12.78 | 1.04 |
| RNANR | 86.20 | **7.38** | 0.85 | 86.20 | 8.54 | 11.45 | 86.20 | 8.54 | 41.61 |
| RNAsubopt | 85.90 | 3.05 | 0.03 | 92.40 | 6.98 | 0.09 | 94.20 | 14.54 | 0.59 |
| Uniform Sampling | 0.00 | 0.00 | 0.00 | 0.00 | 0.00 | 0.00 | 0.00 | 0.00 | 0.00 |

Table S9: Coverage of the 100 highest saddle points associated to the highest barriers between the deepest left and right minima in the barriers tree for ten 80 nt sequences. The sample size is the number of structures that a tool produces in one run per sequence. In addition to the barriers, the basin coverage is shown and the average run time for each tool. Two guiding potentials have been used for `RNAxplorer`. The base pair associated guiding potential, described in the main text (section **??** is (bp gp). The base pair distance based guiding potential (described in Eqn. (12)) is (bpd gp).

| | sample size | | | | | | | | |
|---|---|---|---|---|---|---|---|---|---|
| Tool | $10^3$ | | | $10^4$ | | | $10^5$ | | |
| length: 90 nt | coverage [%] | | | coverage [%] | | | coverage [%] | | |
| max. basins: 34561 | barriers | basins | $\bar{t}$ [$s$] | barriers | basins | $\bar{t}$ [$s$] | barriers | basins | $\bar{t}$ [$s$] |
| RNAxplorer (bp gp) | **81.60** | 2.29 | 0.05 | **91.00** | **9.47** | 0.33 | **98.60** | **31.12** | 2.83 |
| RNAxplorer (bpd gp) | 76.50 | 2.00 | 0.32 | 88.60 | 5.47 | 2.75 | 91.50 | 12.79 | 9.31 |
| RNAlocmin | 78.30 | 1.95 | 38.14 | 90.10 | 7.31 | 41.01 | 95.20 | 24.43 | 68.91 |
| RNAlocopt | 76.40 | 1.66 | 0.86 | 86.90 | 3.88 | 0.93 | 90.70 | 8.06 | 1.36 |
| RNANR | 67.30 | **3.87** | 1.53 | 67.40 | 6.47 | 17.02 | 67.40 | 6.48 | 273.03 |
| RNAsubopt | 79.80 | 1.86 | 0.03 | 87.40 | 5.01 | 0.10 | 92.10 | 11.95 | 0.61 |
| Uniform Sampling | 0.00 | 0.00 | 0.00 | 0.00 | 0.00 | 0.00 | 0.00 | 0.00 | 0.00 |

Table S10: Coverage of the 100 highest saddle points associated to the highest barriers between the deepest left and right minima in the barriers tree for ten 90 nt sequences. The sample size is the number of structures that a tool produces in one run per sequence. In addition to the barriers, the basin coverage is shown and the average run time for each tool. Two guiding potentials have been used for `RNAxplorer`. The base pair associated guiding potential, described in the main text (section **??** is (bp gp). The base pair distance based guiding potential (described in Eqn. (12)) is (bpd gp).

| | sample size | | | | | | | | |
|---|---|---|---|---|---|---|---|---|---|
| Tool | $10^3$ | | | $10^4$ | | | $10^5$ | | |
| length: 100 nt | coverage [%] | | | coverage [%] | | | coverage [%] | | |
| max. basins: 63536 | barriers | basins | $\bar{t}$ [$s$] | barriers | basins | $\bar{t}$ [$s$] | barriers | basins | $\bar{t}$ [$s$] |
| RNAxplorer (bp gp) | **78.60** | 1.67 | 0.06 | **87.40** | **6.64** | 0.41 | **93.50** | **22.25** | 3.72 |
| RNAxplorer (bpd gp) | 74.20 | 1.48 | 0.38 | 84.90 | 4.07 | 3.63 | 88.50 | 9.85 | 10.80 |
| RNAlocmin | 74.80 | 1.46 | 41.19 | 84.30 | 4.61 | 44.11 | 90.20 | 15.48 | 73.61 |
| RNAlocopt | 75.00 | 1.03 | 1.20 | 83.70 | 2.82 | 1.30 | 85.30 | 6.21 | 1.92 |
| RNANR | 70.40 | **2.07** | 2.17 | 70.80 | 3.31 | 18.30 | 70.80 | 3.32 | 562.75 |
| RNAsubopt | 72.70 | 1.40 | 0.02 | 84.60 | 4.00 | 0.11 | 89.00 | 9.56 | 0.70 |
| Uniform Sampling | 0.00 | 0.00 | 0.00 | 0.00 | 0.00 | 0.00 | 0.00 | 0.00 | 0.00 |

Table S11: Coverage of the 100 highest saddle points associated to the highest barriers between the deepest left and right minima in the barriers tree for ten 100 nt sequences. The sample size is the number of structures that a tool produces in one run per sequence. In addition to the barriers, the basin coverage is shown and the average run time for each tool. Two guiding potentials have been used for `RNAxplorer`. The base pair associated guiding potential, described in the main text (section **??** is (bp gp). The base pair distance based guiding potential (described in Eqn. (12)) is (bpd gp).

#### 5.4 Visual assessment of the quality of generated samples

It is not trivial to visualize the conformation space in a Cartesian coordinate system, since secondary structures are not a regular graph. However, it is possible to create projections on a 2-dimensional grid. A classified dynamic programming algorithm, implemented in the tool `RNA2Dfold` can be used to compute either the partition function or the MFE structure for each cell in the 2-dimensional grid of base pair distances to two reference structures. With this projection as ground truth, different sampling methods can be compared and missing important minima in the landscape can be identified.

Figure S14 depicts the local minima (obtained via gradient walks) of $10^6$ structures, for different sampling methods. The corners at the x- and y-axis, represent the ground and the metastable state of the 'sv11' RNA. Only `RNAxplorer` is able to identify the metastable state with the given number of samples (and even with less than $10^6$ samples).

With each tool $10^6$ structures were generated, except for `RNASLopt`. `RNASLopt` does not have any option to set the number of structures. Instead, a parameter $\delta$ is used to adjust the allowed energy range above the MFE. This parameter was increased from 0 to 1000. If the sample size did not change for the next iteration, we took the sample set with the largest size. For the sequences *tpp.* and *lys.* the $\delta$ parameter was set to 30 and 20, respectively. Computations with higher $\delta$ would take longer than 2 days (which does not make any sense to compare with the other tools that create samples within a few seconds). All other sequences could be computed with $\delta$ 1000 within a few seconds but with low sample sizes. Only 1,523 up to 332,357 structures could be generated with `RNASLopt`.

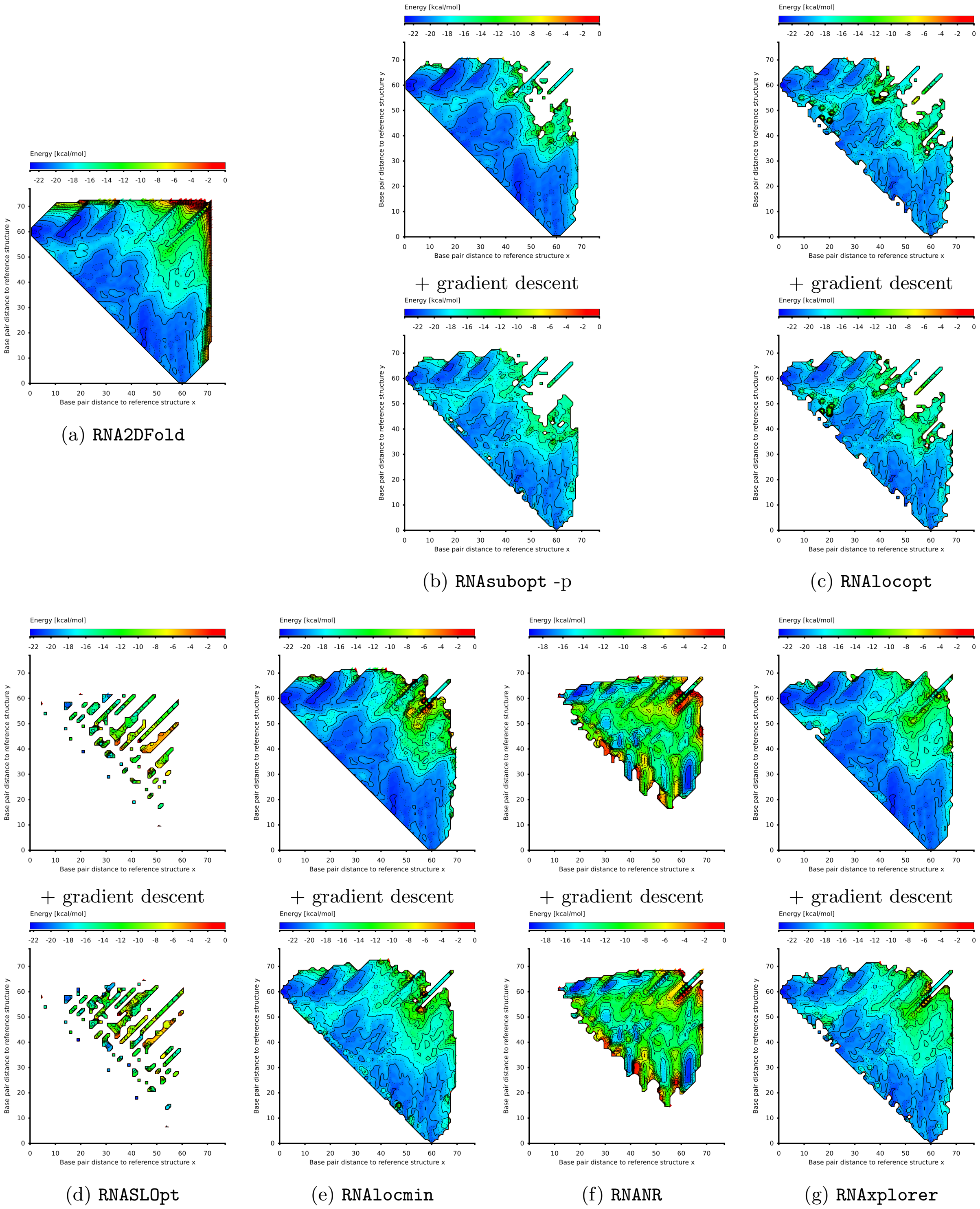

Figure S7: 2D projections of samples and local minima from different methods for the RNA sequence add. The sample size for each tool is $10^6$ (except `RNASLOpt`, for which we have the maximum number of structures with $\delta$ energy up to 1000). (a) Complete Enumeration (`RNA2DFold`), (b) Boltzmann Sampling (`RNAsubopt -p`), (c) Local Optima Sampling (`RNAlocopt`), (d) `RNASLOpt`, (e) Variable Temperature Sampling (`RNAlocmin`), (f) Non-redundant Sampling (`RNANR`), (g) Repellant Sampling (`RNAxplorer`).
The run times of the sampling tools in seconds are: (a) 3.93, (b) 7.21, (c) 33.56, (d) 2.16, (e) 656.49, (f) 34114.19, (g) 51.54

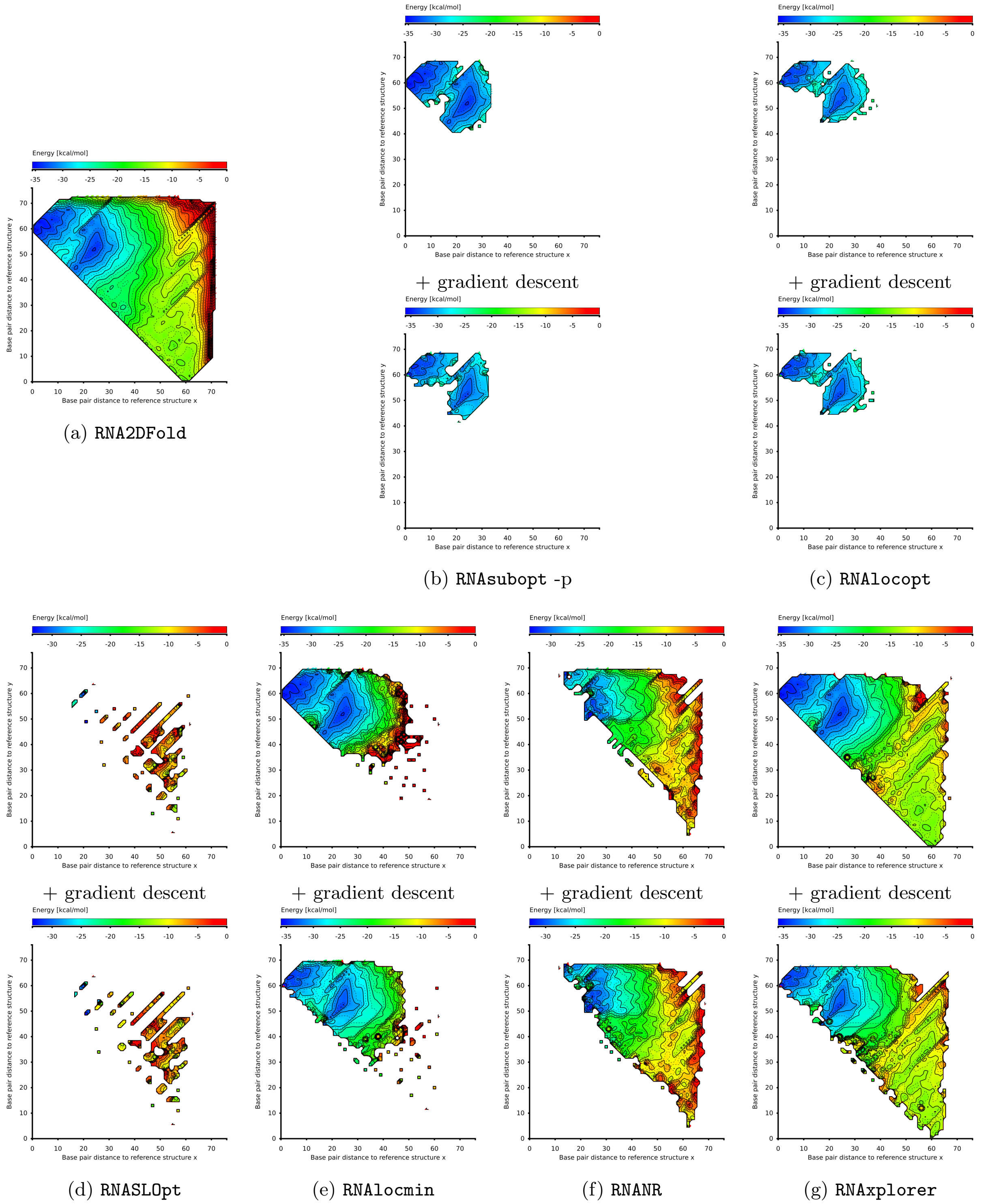

(a) `RNA2DFold` (b) `RNAsubopt -p` (c) `RNAlocopt`

(d) `RNASLOpt` (e) `RNAlocmin` (f) `RNANR` (g) `RNAxplorer`

Figure S8: 2D projections of samples and local minima from different methods for the RNA sequence ydhL. The sample size for each tool is $10^6$ (except `RNASLOpt`, for which we have the maximum number of structures with $\delta$ energy up to 1000). (a) Complete Enumeration (`RNA2DFold`), (b) Boltzmann Sampling (`RNAsubopt -p`), (c) Local Optima Sampling (`RNAlocopt`), (d) `RNASLOpt`, (e) Variable Temperature Sampling (`RNAlocmin`), (f) Non-redundant Sampling (`RNANR`), (g) Repellant Sampling (`RNAxplorer`).
The run times of the sampling tools in seconds are: (a) 3.81, (b) 6.23, (c) 26.32, (d) 3.49, (e) 612.46, (f) 156506.66, (g) 80.32

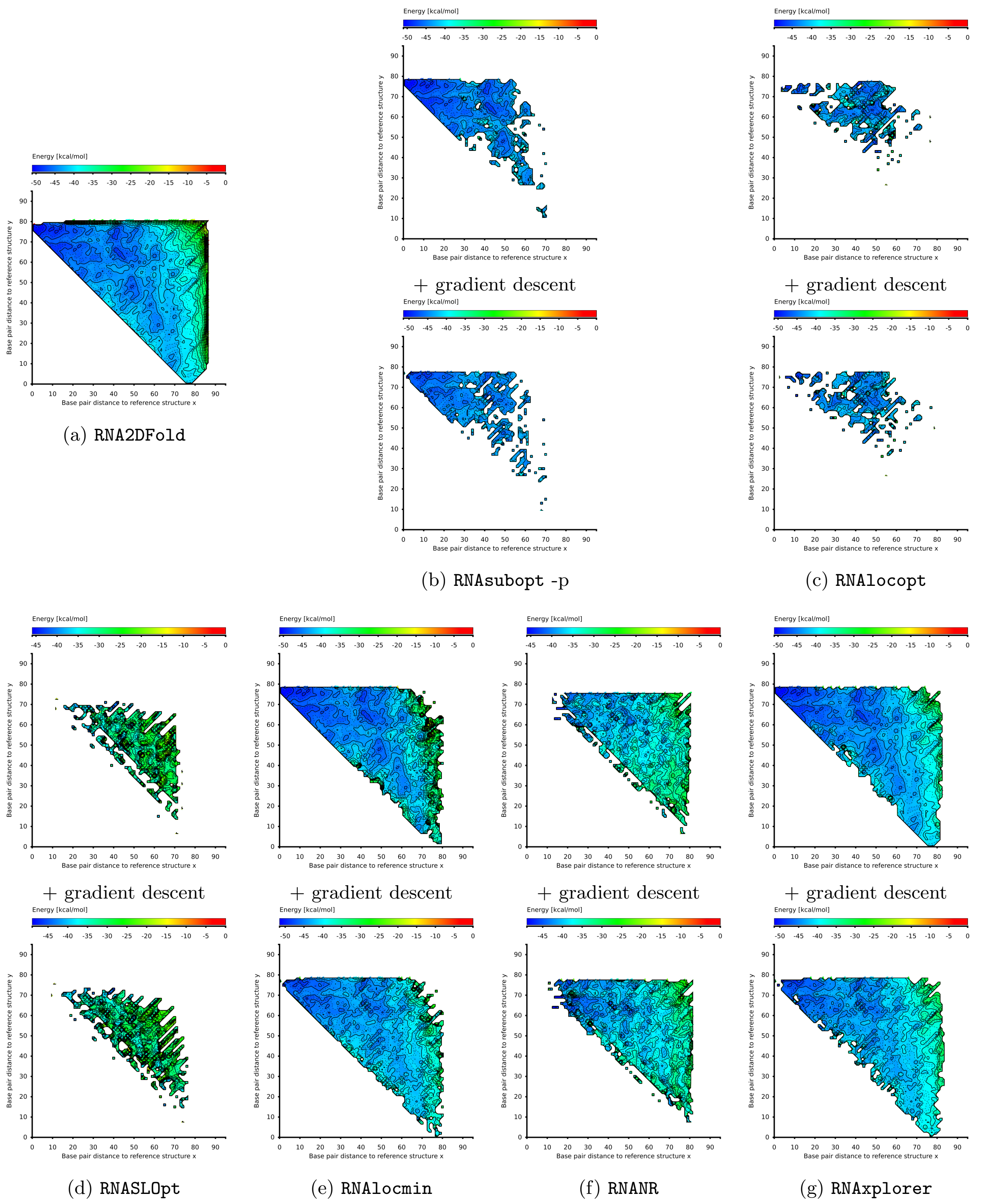

Figure S9: 2D projections of samples and local minima from different methods for the RNA sequence cyc-di-GMP. The sample size for each tool is $10^6$ (except RNASLOpt, for which we have the maximum number of structures with $\delta$ energy up to 1000). (a) Complete Enumeration (RNA2DFold), (b) Boltzmann Sampling (RNAsubopt -p), (c) Local Optima Sampling (RNAlocopt), (d) RNASLOpt, (e) Variable Temperature Sampling (RNAlocmin), (f) Non-redundant Sampling (RNANR), (g) Repellant Sampling (RNAxplorer).
The run times of the sampling tools in seconds are: (a) 8.16, (b) 9.02, (c) 44.36, (d) 38.01, (e) 483.67, (f) 59782.88, (g) 33.89

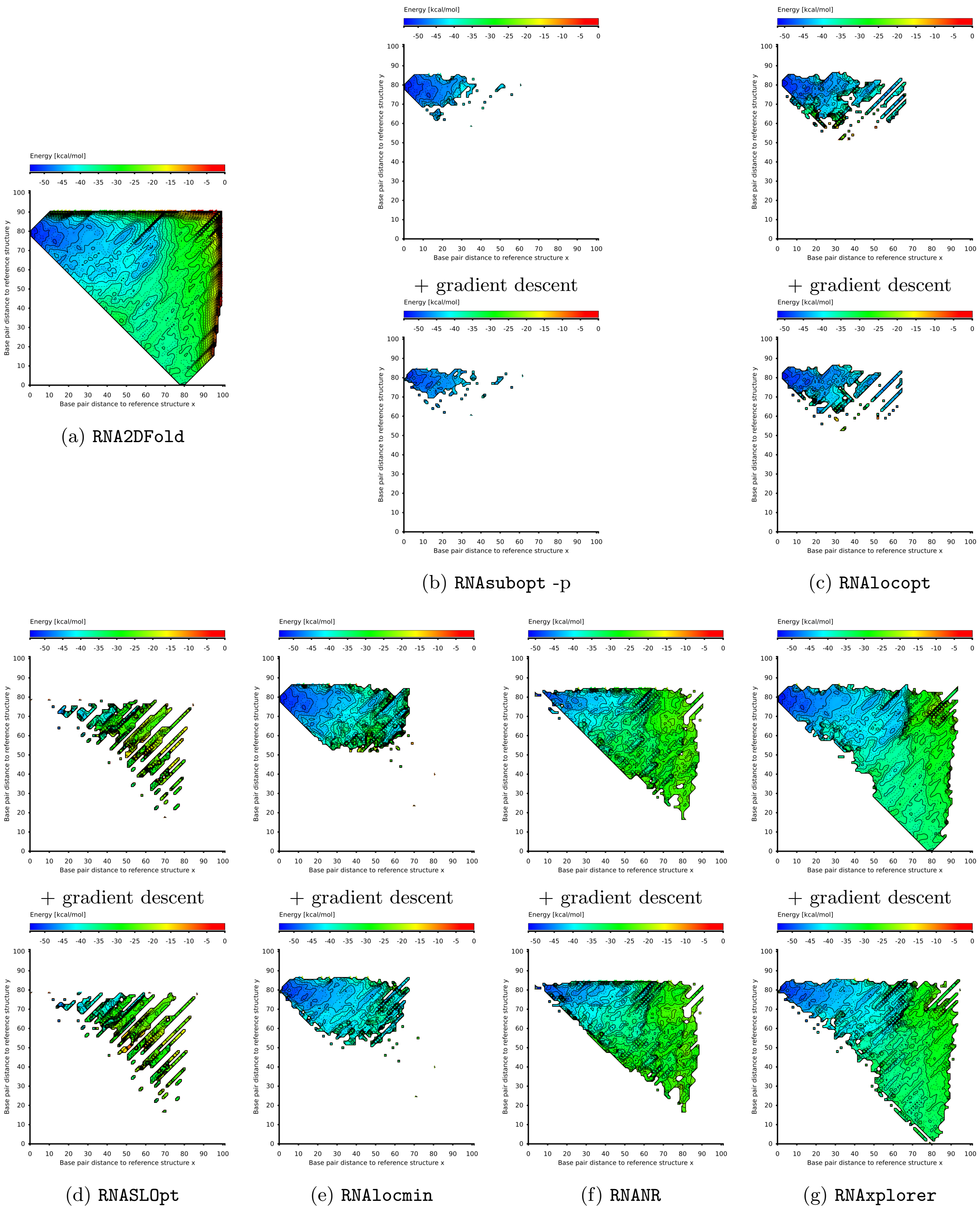

Figure S10: 2D projections of samples and local minima from different methods for the RNA sequence xpt-pubX. The sample size for each tool is $10^6$ (except `RNASLOpt`, for which we have the maximum number of structures with $\delta$ energy up to 1000). (a) Complete Enumeration (`RNA2DFold`), (b) Boltzmann Sampling (`RNAsubopt -p`), (c) Local Optima Sampling (`RNAlocopt`), (d) `RNASLOpt`, (e) Variable Temperature Sampling (`RNAlocmin`), (f) Non-redundant Sampling (`RNANR`), (g) Repellant Sampling (`RNAxplorer`).
The run times of the sampling tools in seconds are: (a) 26.32, (b) 9.11, (c) 82.81, (d) 235.91, (e) 560.27, (f) 38885.86, (g) 52.21

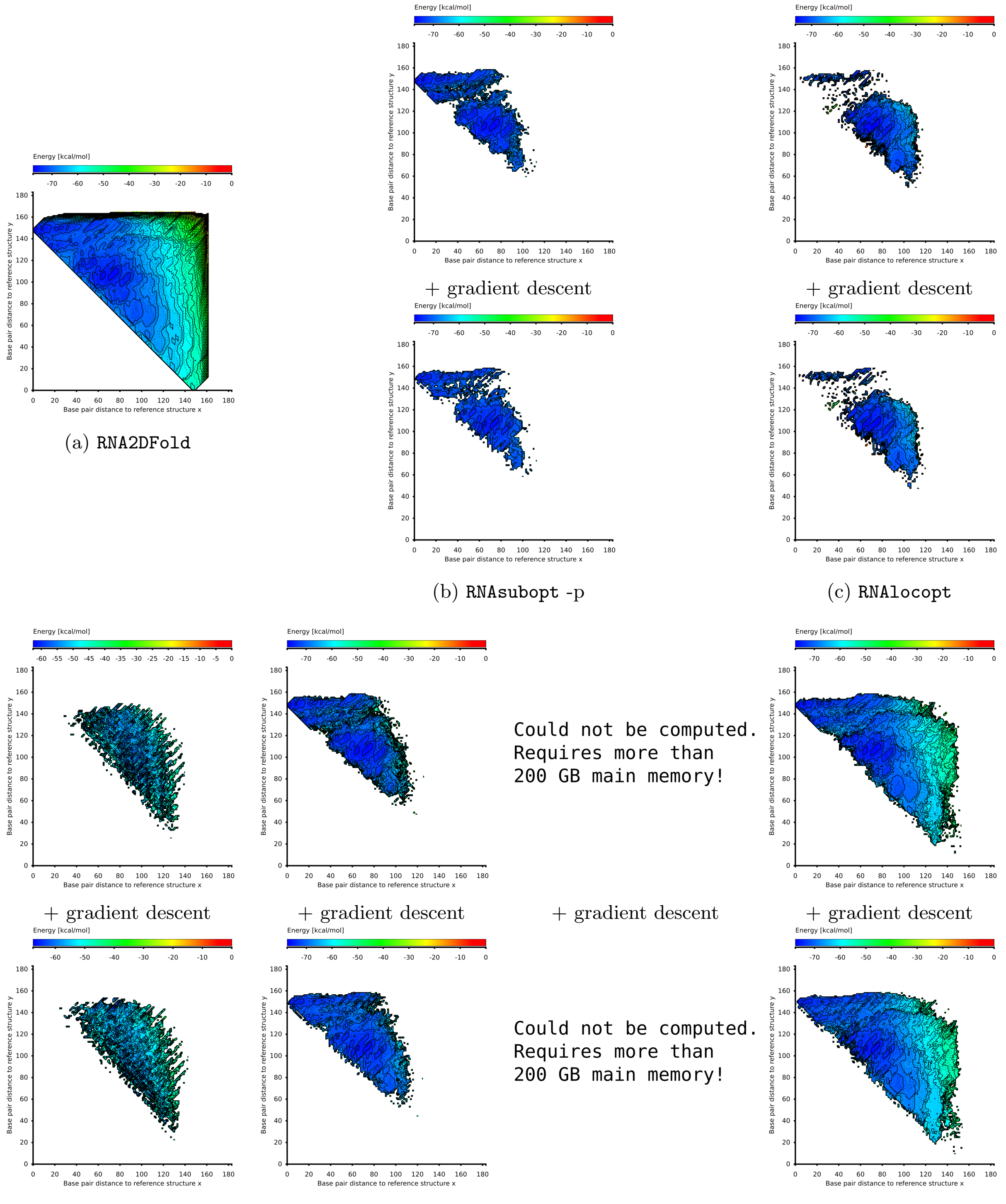

(a) RNA2DFold (b) RNAsubopt -p (c) RNAlocopt (d) RNASLOpt (e) RNAlocmin (f) RNANR (g) RNAxplorer

Figure S11: 2D projections of samples and local minima from different methods for the RNA sequence lysC. The sample size for each tool is $10^6$ (except RNASLOpt, for which we have the maximum number of structures with $\delta$ energy up to 1000). (a) Complete Enumeration (RNA2DFold), (b) Boltzmann Sampling (RNAsubopt -p), (c) Local Optima Sampling (RNAlocopt), (d) RNASLOpt, (e) Variable Temperature Sampling (RNAlocmin), (f) Nonredundant Sampling (RNANR), (g) Repellant Sampling (RNAxplorer).
The run times of the sampling tools in seconds are: (a) 731.10, (b) 14.25, (c) 216.43, (d) 31913.72, (e) 841.54, (f) N.A., (g) 81.08

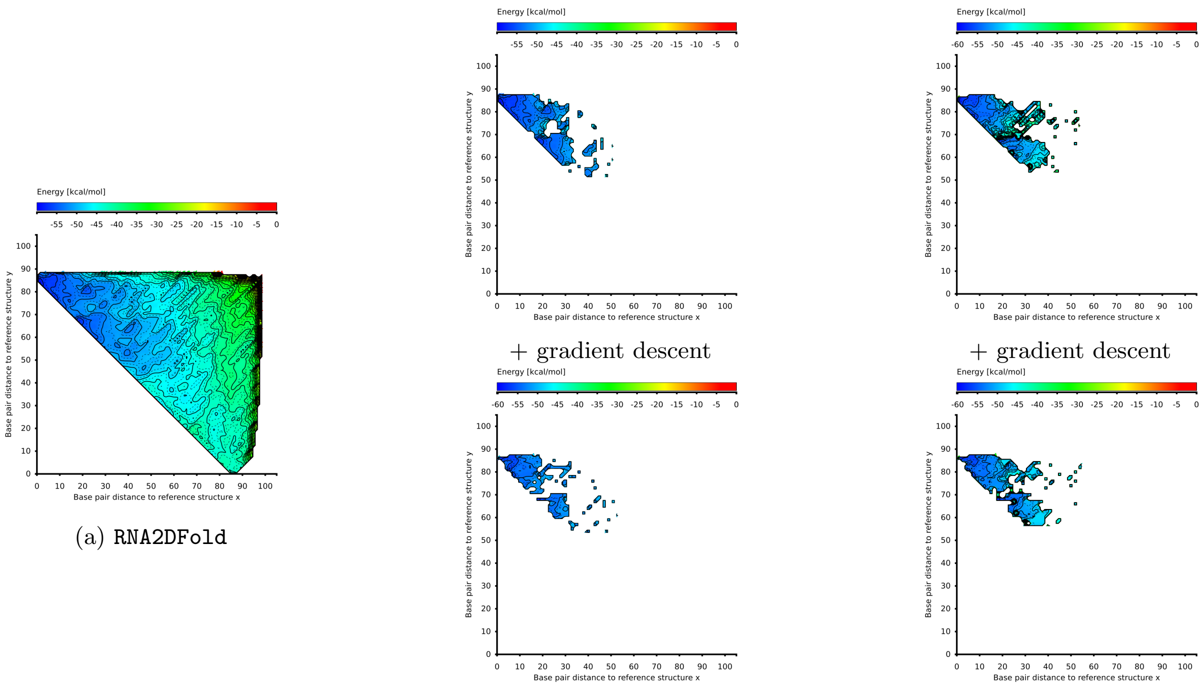

(a) RNA2DFold (b) RNAsubopt -p (c) RNAlocopt

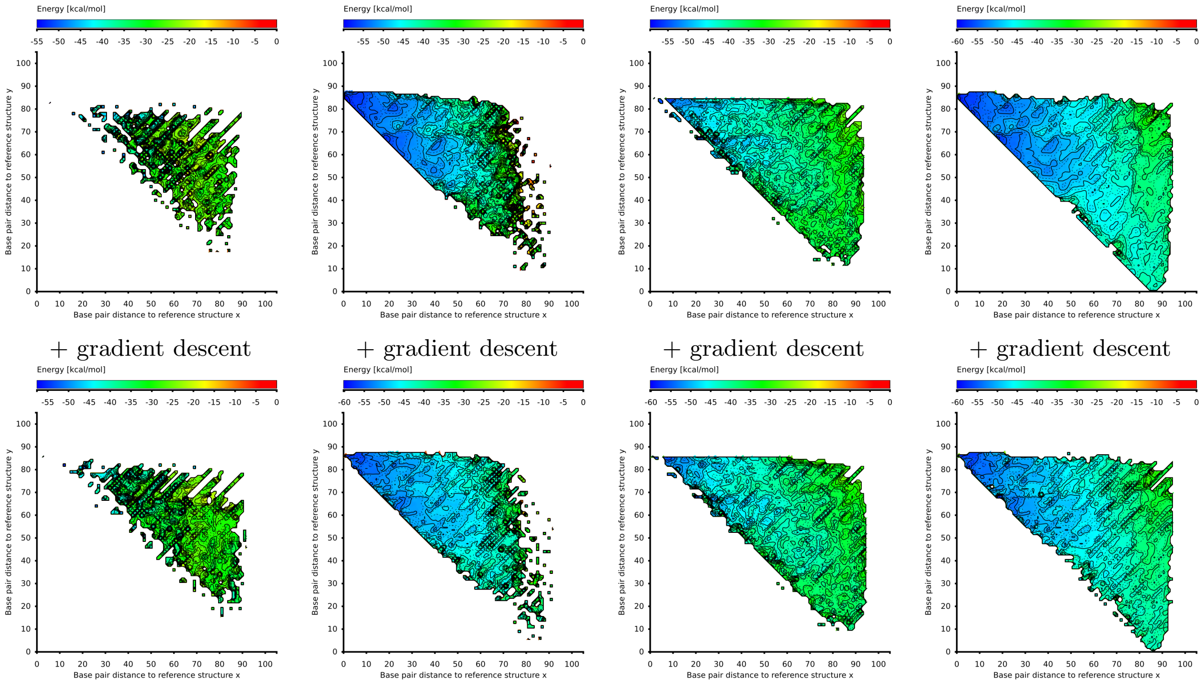

(d) RNASLOpt (e) RNAlocmin (f) RNANR (g) RNAxplorer

Figure S12: 2D projections of samples and local minima from different methods for the RNA sequence MS2. The sample size for each tool is $10^6$ (except RNASLOpt, for which we have the maximum number of structures with $\delta$ energy up to 1000). (a) Complete Enumeration (RNA2DFold), (b) Boltzmann Sampling (RNAsubopt -p), (c) Local Optima Sampling (RNAlocopt), (d) RNASLOpt, (e) Variable Temperature Sampling (RNAlocmin), (f) Non-redundant Sampling (RNANR), (g) Repellant Sampling (RNAxplorer).
The run times of the sampling tools in seconds are: (a) 15.98, (b) 8.98, (c) 130.72, (d) 994.66, (e) 515.99, (f) 116889.91, (g) 48.78

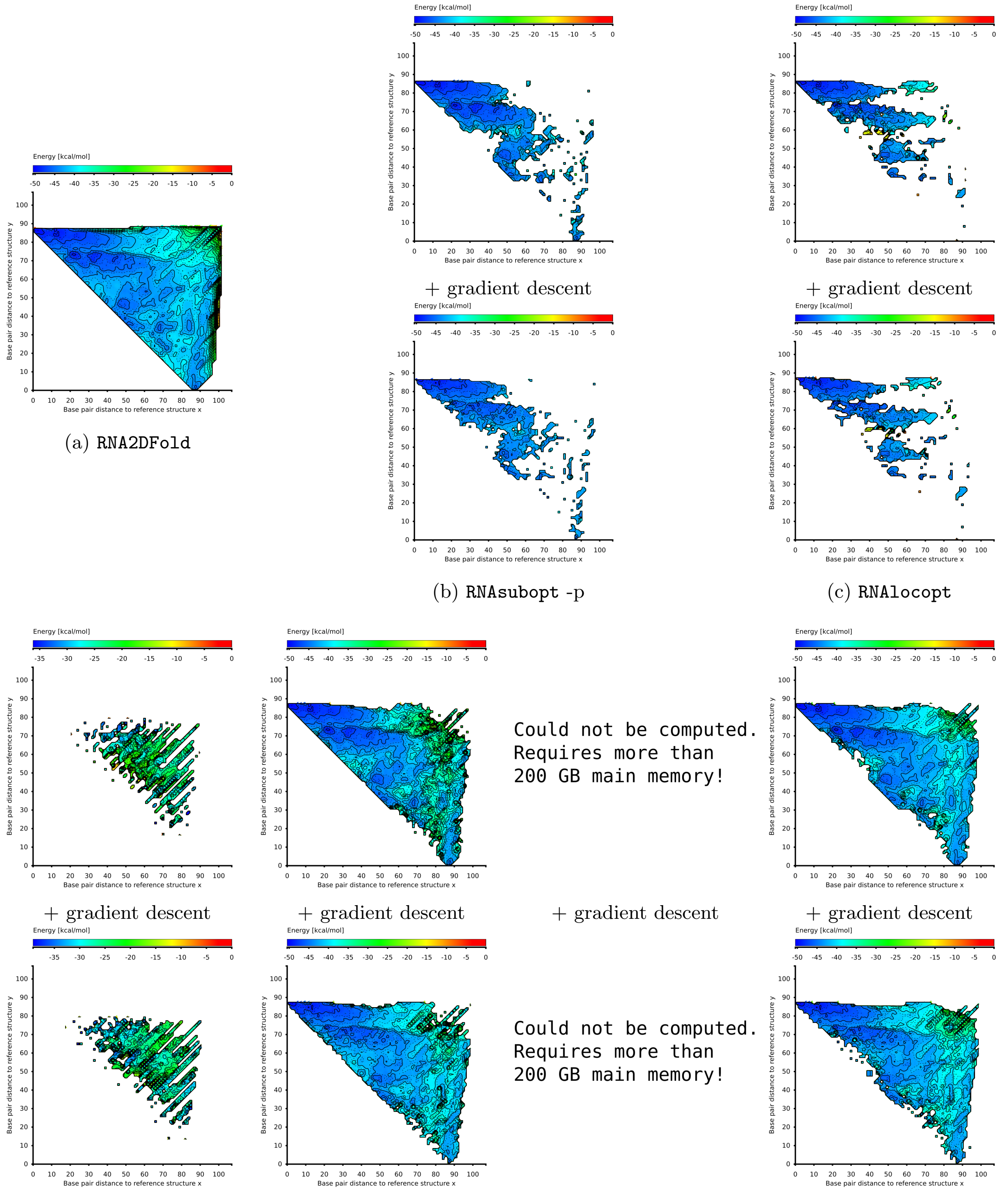

(a) `RNA2DFold` (b) `RNAsubopt -p` (c) `RNAlocopt`

(d) `RNASLOpt` (e) `RNAlocmin` (f) `RNANR` (g) `RNAxplorer`

Figure S13: 2D projections of samples and local minima from different methods for the RNA sequence metE. The sample size for each tool is $10^6$ (except `RNASLOpt`, for which we have the maximum number of structures with $\delta$ energy up to 1000). (a) Complete Enumeration (`RNA2DFold`), (b) Boltzmann Sampling (`RNAsubopt -p`), (c) Local Optima Sampling (`RNAlocopt`), (d) `RNASLOpt`, (e) Variable Temperature Sampling (`RNAlocmin`), (f) Non-redundant Sampling (`RNANR`), (g) Repellant Sampling (`RNAxplorer`).
The run times of the sampling tools in seconds are: (a) 14.32, (b) 8.02, (c) 46.90, (d) 864.23, (e) 546.96, (f) N.A., (g) 49.83

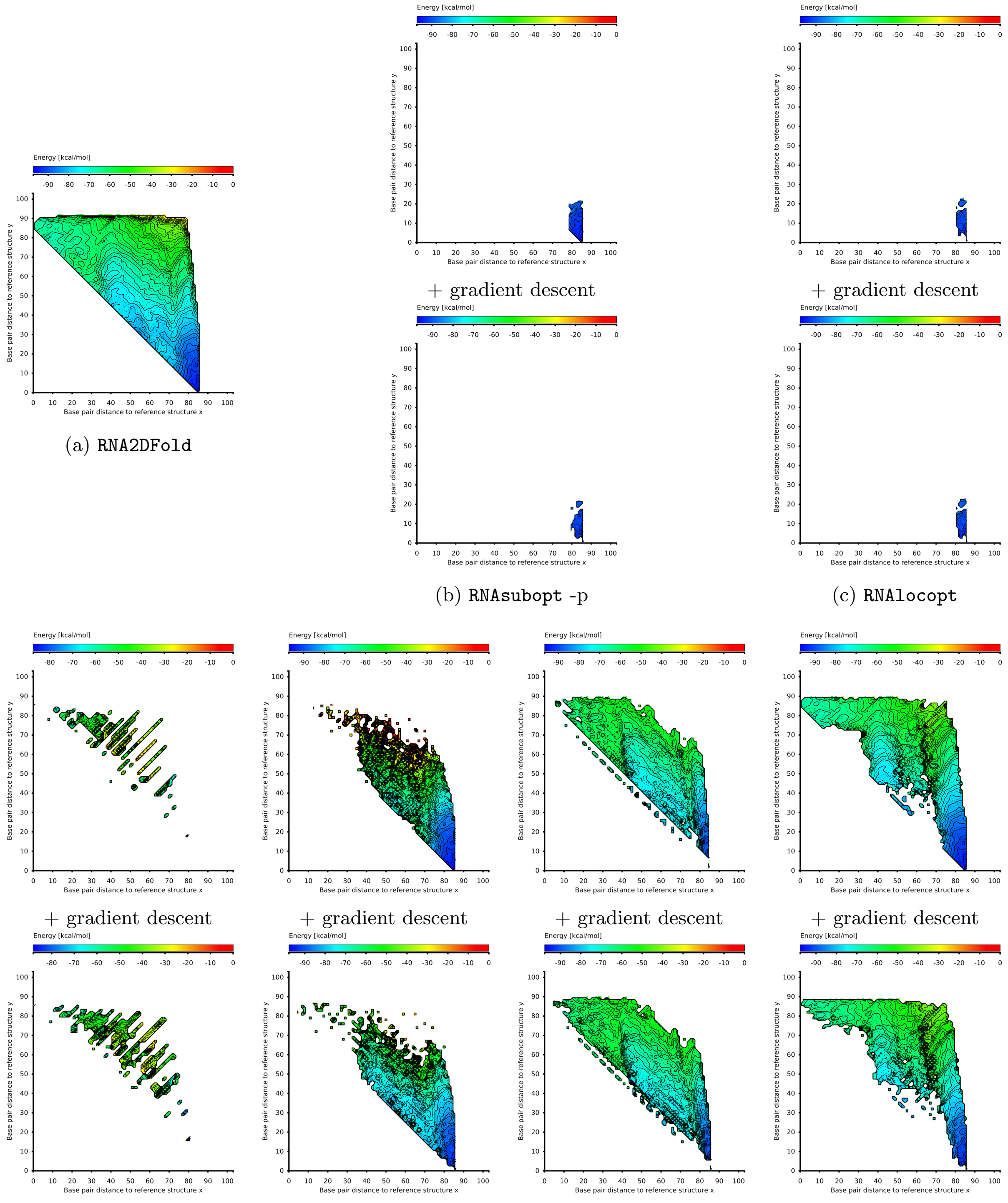

(a) `RNA2DFold`

(b) `RNAsubopt -p`

(c) `RNAlocopt`

(d) `RNASLOpt`

(e) `RNAlocmin`

(f) `RNANR`

(g) `RNAxplorer`

Figure S14: 2D projections of samples and local minima from different methods for the RNA sequence SV-11. The sample size for each tool is $10^6$ (except `RNASLOpt`, for which we have the maximum number of structures with $\delta$ energy up to 1000). (a) Complete Enumeration (`RNA2DFold`), (b) Boltzmann Sampling (`RNAsubopt -p`), (c) Local Optima Sampling (`RNAlocopt`), (d) `RNASLOpt`, (e) Variable Temperature Sampling (`RNAlocmin`), (f) Non-redundant Sampling (`RNANR`), (g) Repellant Sampling (`RNAxplorer`).
The run times of the sampling tools in seconds are: (a) 4.16, (b) 6.75, (c) 21.81, (d) 115.99, (e) 487.73, (f) 4285.81, (g) 27.87

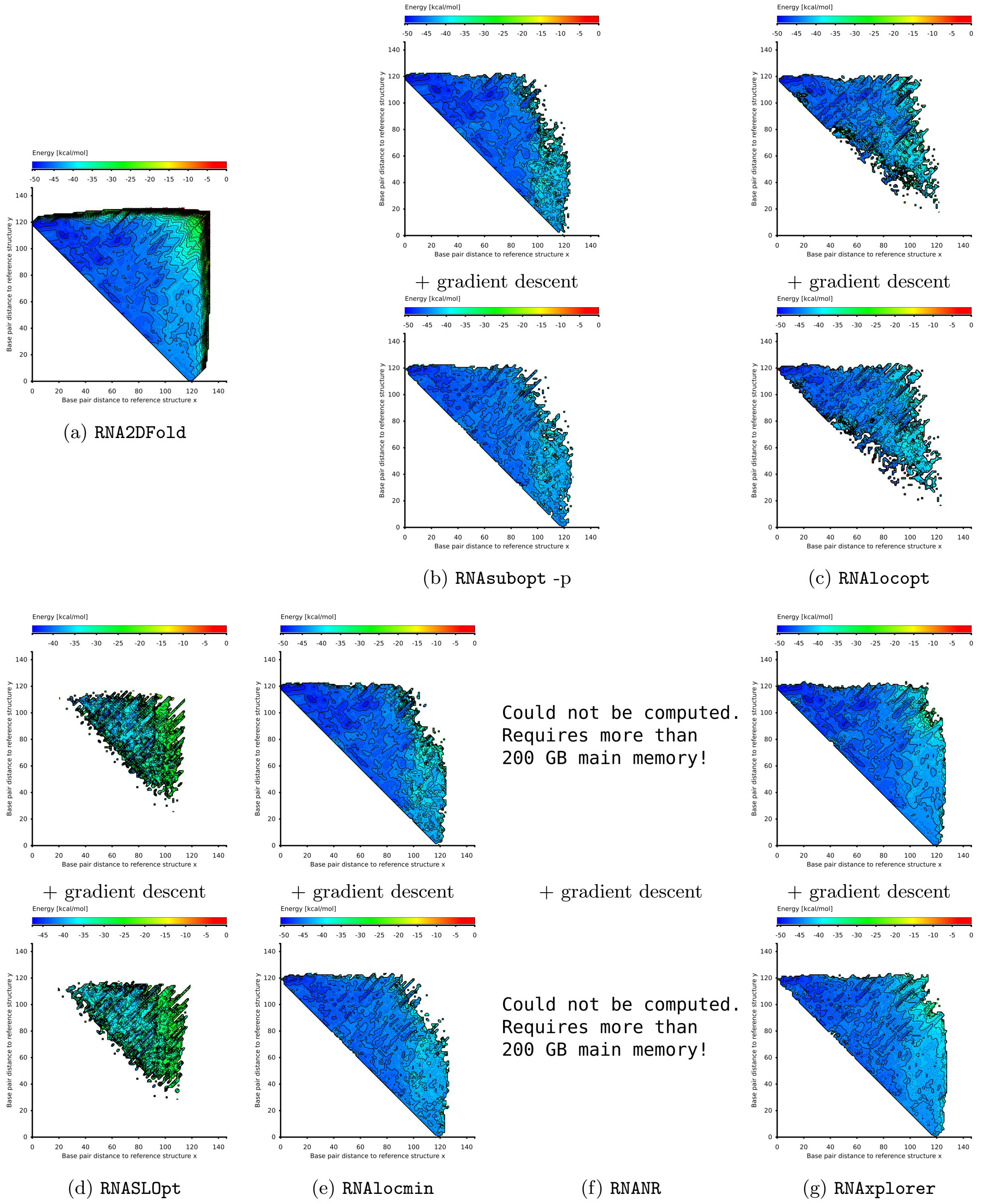

Figure S15: 2D projections of samples and local minima from different methods for the RNA sequence thiamine. The sample size for each tool is $10^6$ (except RNASLOpt, for which we have the maximum number of structures with $\delta$ energy up to 1000). (a) Complete Enumeration (RNA2DFold), (b) Boltzmann Sampling (RNAsubopt -p), (c) Local Optima Sampling (RNAlocopt), (d) RNASLOpt, (e) Variable Temperature Sampling (RNAlocmin), (f) Non-redundant Sampling (RNANR), (g) Repellant Sampling (RNAxplorer).
The run times of the sampling tools in seconds are: (a) 182.52, (b) 14.76, (c) 154.47, (d) 91944.85, (e) 663.17, (f) N.A., (g) 69.59

##### 5.5 Coverage of Distance Classes

Since metrics such as the number of distinct structures or local minima only partially capture the quality of samples, we extended the concept of distance classes in 2D projections for our benchmarks.

This notion of diversity relies on the distance of the samples to two reference structures and the quality part of the measure is either the minimum free energy or the fraction of the partition function. The following analysis attempt to evaluate the number of cells for which the sample achieves good quality, i.e. represents a good approximation of the true value.

Before we define our metrics, let us define $maxC$ as the number of distance classes populated by two reference structures $\hat{s}_1$ and $\hat{s}_2$ for a sequence $\sigma$, such that

$$maxC(\sigma) = |\{(d_1, d_2) \in \mathbb{N}^2 \mid \mathcal{C}^{d_1,d_2} \neq \varnothing\}|.$$

If several sequences are used with this measure, the number $n\mathcal{C}(\sigma)$ of satisfactory classes can be largely heterogeneous, biasing our analysis towards large structures. We thus normalize the contribution of each sequence, dividing it by $max\mathcal{C}(\sigma)$. In other words, we average in our computation the proportion of classes for which our the true MFE is correctly approximated. Overall, for a set of sequences $\mathcal{B}$, our final quality measure $qm$ (used in the main paper in Figure 4) is:

$$qm = \sum_{\sigma \in \mathcal{B}} \frac{n\mathcal{C}_\vartheta(\sigma)}{max\mathcal{C}(\sigma)} / |\mathcal{B}| \tag{19}$$

###### 5.5.1 With respect to the cell MFE

In order to compare sampling methods with different move sets and energies, we compute a unified coarse-graining of each sample (via gradient walks). Moreover, in order to allow some level of flexibility, we set thresholds $\vartheta$ of 0 and 5 $kcal/mol$ above the MFE for each class are used, to compare sampling methods that use different energies in a unified manner.

Let $\mathcal{S}$ be a sampled set of structures and $\sigma$ a sequence, the number of cells $n\mathcal{C}_\vartheta(\sigma)$ having an MFE comparable to the optimal one (up to a threshold $\vartheta$), can be formally defined as

$$n\mathcal{C}_\vartheta(\sigma) = \left| \left\{ d_1, d_2 \mid \min_{s \in \mathcal{S} \cap \mathcal{C}^{d_1,d_2}} E(s) - MFE^{d_1,d_2} \leq \vartheta \right\} \right| \tag{20}$$

Figure S16 shows the number of cells having an MFE comparable with the true MFE as a function of sample size for all tools an sequences. This corresponds to the measure described in equation 20. Here, we applied two different thresholds of $\vartheta_1 = 0$ kcal/mol and $\vartheta_2 = 5$ kcal/mol. Figure S17 shows the corresponding average of the quality measure as described in Eqn. 19.

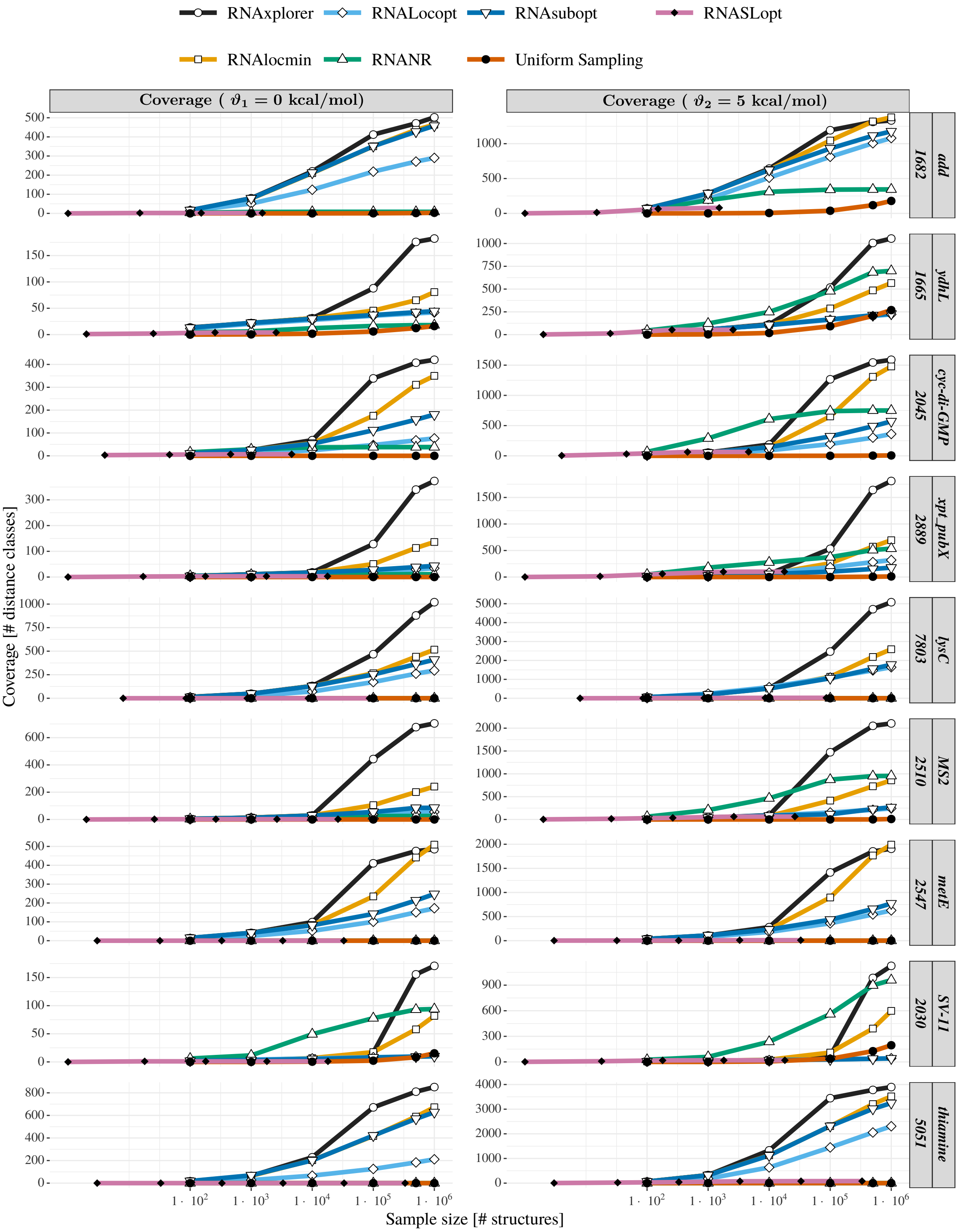

Figure S16: MFE distance class coverage of landscape projections in absolute numbers. Each line corresponds to one sequence. The right most gray boxes contain the sequence names. The number next to the name is the maximal number of distance classes for this sequence. The left column contains the measure with exact MFEs ($\vartheta_1 = 0$ kcal/mol), the right column allows for a threshold $\vartheta_2 = 5$ kcal/mol above the true MFE.

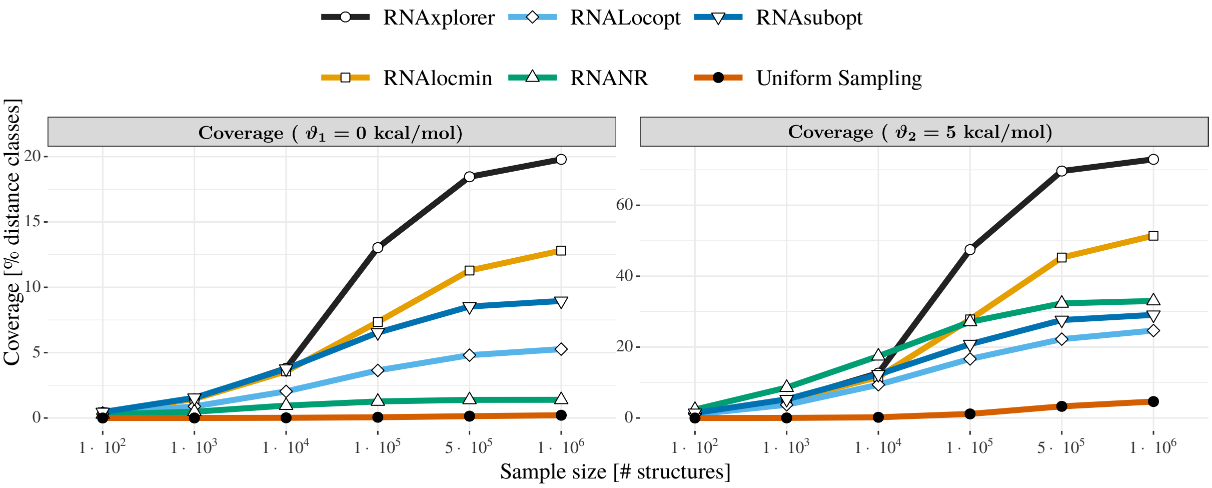

Figure S17: Coverage of distance classes with the MFE measure averaged over the 9 benchmark sequences (10 runs per tool and sequence). Detailed results per sequence and tool are listed in the supplementary material (figure S16).

##### 5.5.2 With respect to the partition function of cells

In this metric, we compare the partition functions for all spots in the projection to its true value. In particular, we report the number of spots where the partition function of the samples is higher than a certain fraction of the partition function for all structures on this spot.

Let $\mathcal{S}$ be a sampled set of structures and $\sigma$ a sequence, we consider the number $n\mathcal{C}_\vartheta(\sigma)$ having a partition function in $\mathcal{C}$ that is comparable to the optimal one (up to a threshold $\vartheta$). It can be formally defined as

$$n\mathcal{C}_\vartheta(\sigma) = \left| \left\{ d_1, d_2 \mid \frac{\sum_{\substack{s \in \mathcal{S} \text{ s.t.} \\ s \in \mathcal{C}^{d_1,d_2}}} e^{-\beta E(s)}}{Z^{d_1,d_2}} \geq \vartheta \right\} \right| \tag{21}$$

In Figure S18 we show the respective results for all 6 tools in our comparison for each of the 9 sequences of the benchmark set (cf. Eqn 21). Here, we applied thresholds of 10% ($\vartheta_1 = 0.1$) and 50% ($\vartheta_2 = 0.5$). An average with the same thresholds over all benchmark sequences is shown in Figure S19 (cf. Eqn 19).

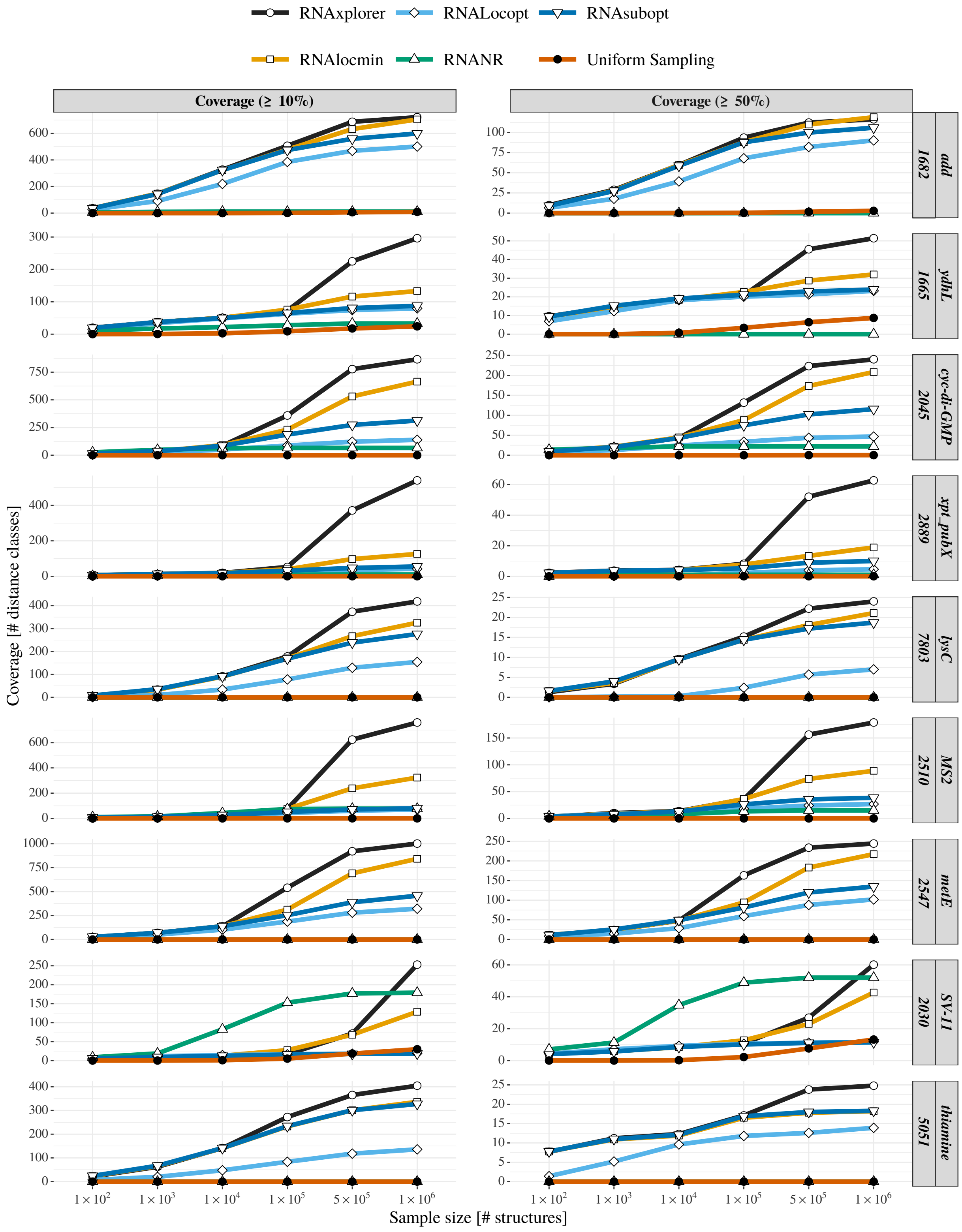

Figure S18: Partition function distance class coverage of landscape projections in absolute numbers. Each line corresponds to one sequence. The right most gray boxes contain the sequence names. The number next to the name is the maximal number of distance classes for this sequence. The left column contains the measure with at least 10% ($\vartheta_1 = 0.1$) partition function coverage, the right column is restricted to a threshold of at least 50% ($\vartheta_2 = 0.5$) of the true partition function.

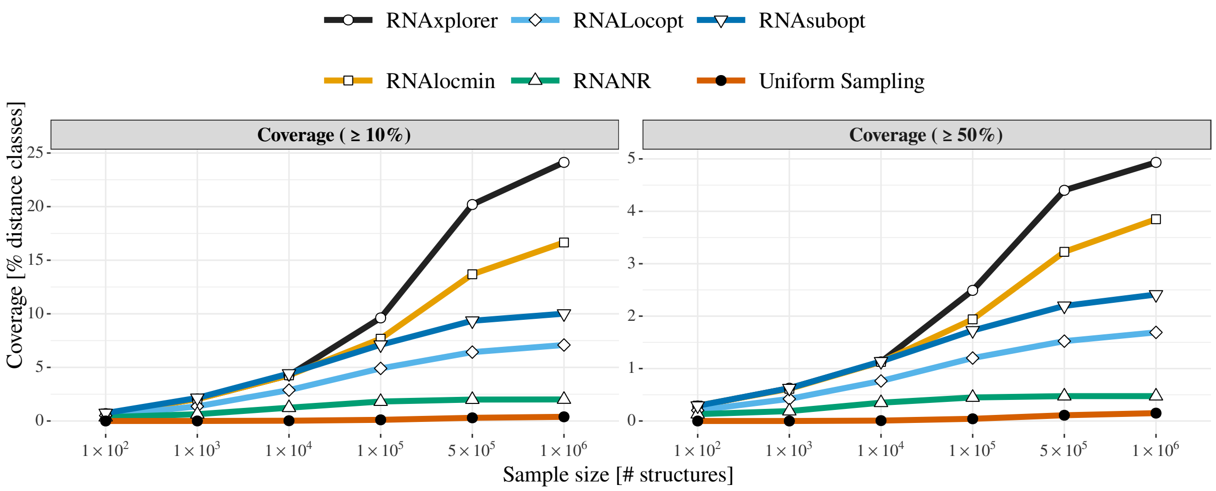

Figure S19: Coverage of distance classes measure with partition function thresholds of 10% ($\vartheta_1 = 0.1$) and 50% ($\vartheta_2 = 0.5$), averaged over the 9 benchmark sequences (10 runs per tool and sequence). Detailed results per sequence and tool are listed in figure S18.
